# Coexisting Y haplotypes reveal cyclic sex chromosome differentiation around a shared sex-determining region

**DOI:** 10.64898/2025.12.31.697208

**Authors:** Fantin Carpentier, Antoine Houtain, Ezgi Unal, Kelsey L. Doucette, Ricard Fontserè, Alan Brelsford, Melissa A. Toups, Nicolas Perrin, Paris Veltsos, Wen-Juan Ma

## Abstract

Sex chromosome evolution is often viewed as a unidirectional process in which recombination suppression expands around a sex-determining locus, leading eventually to Y-chromosome degeneration. Yet many vertebrates, including frogs, retain homomorphic sex chromosomes and polymorphic differentiation despite long-term recombination suppression, which is quite puzzling. Here, we combine pooled whole-genome resequencing, doubled-haploid YY genomes, and RNA-seq to understand the dynamics of homomorphic sex chromosome evolution in a Swiss Alpine population of the common frog, *Rana temporaria,* where multiple Y haplotypes coexist. We show that two fully differentiated Y haplotypes carry exceptionally large nonrecombining regions spanning ∼90% of the sex chromosome (∼618–625 Mb), whereas semi-differentiated Y haplotype retains only a 4.64 Mb *Dmrt1*-linked sex-determining region, and XX males are genetically indistinguishable from XX females. The large NRRs show limited degeneration, with no evidence of gene loss, Y-linked copy decay, faster-X evolution or enrichment of sex-biased genes, and only modest repeat accumulation. Surprisingly, the coexisting Y haplotypes are neither independent sex-determining systems nor successive stages of a single degeneration trajectory. Instead, they share an ancestral *Dmrt1*-linked region whose X–Y divergence dates to ∼3.47–5.70 Mya, while present-day Y copies of this region remain nearly identical and the large NRRs have accumulated private mutations independently. We thus propose that the Y chromosome undergoes cyclic dynamics of differentiation: extreme heterochiasmy promotes rapid large NRRs formation, sex-reversed XY females recurrently restore X–Y recombination, gene conversion and/or rare Y-Y recombination may homogenize Y copies at the *Dmrt1*-linked region, and renewed male transmission drives subsequent Y-specific differentiation. The findings provide a genome-wide resolution to a central paradox in sex chromosome evolution: how sex chromosomes can experience extensive recombination suppression yet remain homomorphic and only weakly degenerated. By showing that heterochiasmy and sex reversal uncouple the physical extent, age, and degeneration of sex-linked regions, our study reframes homomorphic sex chromosomes as dynamic, repeatedly regenerated systems shaped by alternating phases of recombination suppression, restored X–Y recombination and renewed Y differentiation.

**Significance statement:** Why many vertebrate sex chromosomes remain homomorphic despite long-term recombination suppression remains unresolved. Using whole-genome population genomics, doubled-haploid YY genomes, and transcriptomics in the common frog, we show that multiple coexisting Y haplotypes in a single pond share an old *Dmrt1-*linked sex-determining region but carry younger, independently differentiated Y chromosomes. This genome-wide resolution reveals that extreme heterochiasmy, where recombination is largely restricted to chromosome ends in males, can rapidly generate very large nonrecombining regions, while sex-reversed XY females can restore X–Y recombination and reduce divergence outside the *Dmrt1*-linked region. These opposing processes generate cyclic dynamics of Y differentiation rather than a directional path towards degeneration. Our study provides a mechanistic explanation for the persistence of co-existing multiple Y haplotypes in one population, maintenance of homomorphic sex chromosomes, and a broadly relevant framework for sex chromosome evolution in lineages with heterochiasmy and sex reversal.

## Introduction

Sex chromosomes have originated repeatedly across the Tree of Life (1–3), and their evolution is central in models of sexual selection, sex-specific dispersal, genomic conflict, adaptation, and speciation (4–7). Sex chromosomes demonstrate a remarkable diversity in their rate and level of genetic differentiation (1, 8–10). The classical model of sex chromosome evolution begins with the acquisition of a sex-determining locus on one member of an autosomal pair via mutation. Recombination suppression can then be favoured around this locus, particularly when linkage preserves favourable combinations of the sex-determining locus and sexually antagonistic alleles (11, 12). This causes the sex-limited Y/W chromosome to differentiate and progressively diverge in structure and function from the still-recombining X/Z chromosome (13–16). In this framework, the nonrecombining region expands in steps, generating evolutionary strata and eventually exposing the sex-limited Y or W chromosome to degeneration through repeat accumulation, reduced effective population size, gene loss and dosage imbalance (Fig. 1a) (17, 18). This can result in highly degenerated Y/W chromosomes which have lost most of their original gene content, as observed in most mammals (16–18), most birds (19, 20), *Drosophila* (6, 21, 22) and other insect lineages (23, 24).

**Fig. 1.**
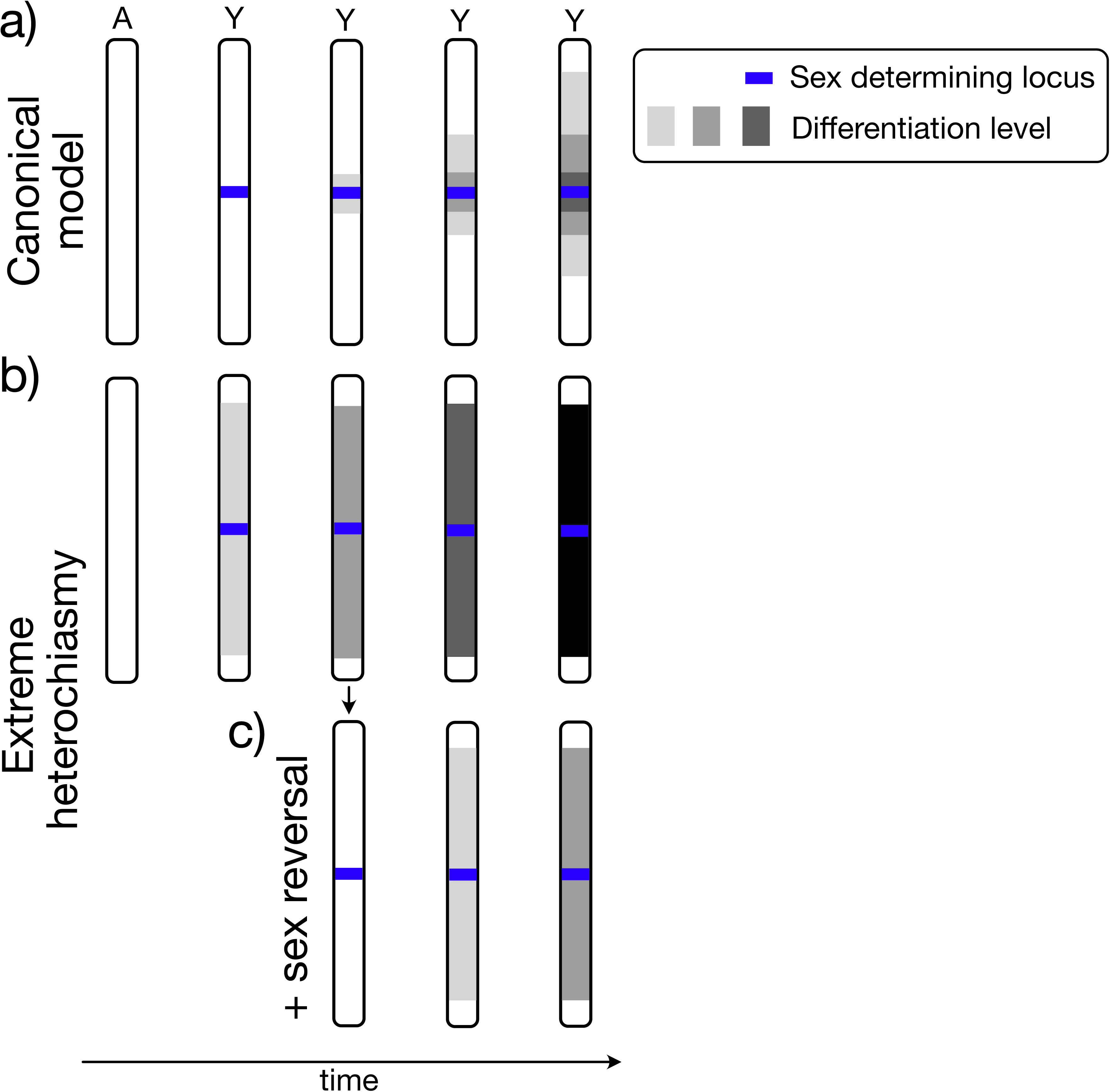
Divergence trajectories of the Y chromosome under the canonical model with chiasmata versus extreme heterochiasmy. a) In the canonical model, the origin of a sex-determining allele on the proto-Y initiates localized X–Y divergence through reduced recombination and the build-up of sexually antagonistic alleles, with recombination suppression subsequently expanding to form progressively differentiated strata. b) Under extreme heterochiasmy, where male recombination is confined to telomeric sites, the establishment of a sex-determining locus produces immediate, chromosome-wide X–Y differentiation except at the recombining ends. XY sex-reversal events transiently restore X–Y recombination and reset differentiation outside the sex-determining region; the resulting recombinant Y haplotype then accumulates independent mutations and follows a novel divergence trajectory.

This classical model has been highly influential. Its hallmarks are the inevitable gene loss and degeneration of the Y/W chromosomes over evolutionary time (15, 17, 21), and the appeared correlation of Y degeneration with evolutionary time (17, 25). However, recent empirical studies have revealed substantial variation in the extent of recombination suppression and the rate of sex chromosome differentiation and degeneration across the Tree of Life (26–31). Gane gains (along with loses) have been found, for example, on the differentiated neo-Y of *Drosophila miranda* (32). There is also strong heterogeneity in the rate and differentiation of sex chromosomes both among closely related and within species (19, 33–38). It seems that the age of sex chromosomes does not correlate well with the degree of recombination suppression, as sex chromosomes can remain homomorphic or maintain a large recombining region (the pseudoautosomal region, PAR) despite long divergence times (e.g. 25-150 million years in ratites) (33, 34, 36, 38–42).

The classical model has been refined by studies of young and homomorphic sex chromosomes. Recent syntheses emphasize that recombination often persists over much of a sex chromosome, that PAR boundaries can be polymorphic or shift over time, and that the physical size, evolutionary age, and degree of molecular differentiation of a sex-linked region need not covary (43–45). Forces other than sexually antagonistic selection have been proposed to shape early sex chromosome evolution (46, 47). They include the pre-existing recombination landscape (43, 45), differentiation due to neutral accumulation of sequence divergence adjacent to the sex-determining locus (48–50); *cis* and *trans* gene expression regulators maintaining recombination suppression and sex chromosome differentiation (51–53).

Pre-existing recombination landscapes may strongly shape the early stage of sex chromosome evolution, particularly in taxa with pronounced heterochiasmy. Across animals and plants, recombination often differs between the sexes (heterochiasmy). In the most extreme case, one sex lacks genome-wide meiotic recombination (achiasmy), as in XY males in *Drosophila* and ZW females of butterflies and moths (54, 55). In other systems, recombination is restricted to chromosome ends (extreme heterochiasmy) in one sex (34, 56–61). Under this architecture, a newly evolved male-determining locus located away from chromosome ends can immediately become embedded within a large NRR, without requiring other recombination modifiers such as stepwise inversions, which are commonly invoked in classical models. Thus, recombination architecture itself may determine the initial physical scale of sex linkage (Fig. 1b). This mechanism does not rely on sexually antagonistic selection and predicts that large NRRs can arise rapidly and therefore need not be old or highly degenerated (57, 58, 60).

The evolutionary consequences of such recombination landscapes depend on whether suppressed recombination is stable or can be restored. The “fountain-of-youth” model proposes that, when recombination follows phenotypic rather than genotypic sex, sex-reversed XY females can recombine X and Y chromosomes during female meiosis, purging deleterious mutations and eroding Y-linked differentiation (62). Rapid sex chromosome turnover, where closely related species use different chromosome pairs to determine sex, results in low sex chromosome differentiation if it occurs faster than sex chromosomes can differentiate as it restores homomorphy to the ancestral sex chromosome pair (XX or YY) (34, 63–66). Together, sex-reversal-mediated recombination and/or sex chromosome turnover may help explain why many amphibians (67, 68), fishes (36, 69–72), many lizards (35), and flowering plants (9, 53, 73, 74) retain largely homomorphic sex chromosomes with limited degeneration. More broadly, these processes align with recent views of young sex chromosomes as dynamic regions in which fully sex-linked regions, partially sex-linked regions, and PARs can shift through time rather than progressing along a single irreversible path towards degeneration (43, 45).

Anurans (frogs and toads) provide a powerful system for testing how pre-existing recombination landscapes interact with “fountain of youth” processes to shape the sex chromosome differentiation and dynamics (Fig. 1b). Many anuran lineages combine extreme heterochiasmy, labile sex determination, and over 78% of studied anurans possess largely homomorphic sex chromosomes (30, 47, 67, 75, 76). Specifically, in anurans, male recombination is often concentrated near chromosome ends, whereas female recombination is more broadly distributed (34, 59, 60, 67). A male-determining locus that arises in the central portion of a chromosome could therefore rapidly define a large Y-linked NRR in male meiosis, which would be eroded outside the sex-determining region via XY sex reversals (Fig. 1b). Repeated cycles of Y inheritance through XY males and sex-reversed XY females would therefore generate cycles of expansion, recombination restoration, and renewed differentiation.

Sex chromosomes can show substantial intraspecific heterogeneity in X–Y differentiation and the European common frog, *Rana temporaria*, is a striking example of this (34, 77–79). *R. temporaria* has extensive intraspecific variation in (homomorphic) X–Y differentiation, including fully differentiated, semi-differentiated, and apparently undifferentiated Y haplotypes (78–83). In certain populations, multiple Y haplotypes coexist within the same pond (78–81, 84). Previous work also demonstrated that X–Y recombination in *R. temporaria* depends on phenotypic sex: rare XY females can recombine X and Y chromosomes, and XY females appear to have no detectable reproductive disadvantage (75, 79, 84). These studies relied largely on a small number of sex-linked markers, leaving unresolved what the coexistence of multiple Y haplotypes reveals about the tempo and reversibility of sex chromosome evolution. The central question is whether these haplotypes represent successive stages of Y differentiation, independent origins of sex determination, or divergent Y backgrounds that share an ancestral sex-determining region which differ in history of recombination suppression, X–Y recombination, and mutation accumulation. Resolving these alternatives requires distinguishing divergence between X and Y from divergence among Y haplotypes, while also defining NRR boundaries and ages, assessing the extent of degeneration, and identifying the master sex-determining locus. Resolving these alternatives allows us to understand the broader question in sex chromosome evolution of whether young sex chromosomes evolve as integrated, irreversible units or as dynamic genomic mosaics shaped by shared ancestry, recombination suppression, and recombination restoration.

Here, we sequence pools of different Y haplotypes and doubled-haploid YY genomes, as well as transcriptomes from a single natural population of *R. temporaria* to examine how extreme heterochiasmy and recurrent sex reversal shape Y-chromosome evolution. By analysing X–Y differentiation and divergence, Y–Y divergence, NRR boundaries, degeneration, the sex-determining region and its age, this study evaluates the genomic consequences of extreme heterochiasmy and “fountain-of-youth” dynamics at high resolution. We infer the evolutionary history of multiple coexisting Y haplotypes and discuss their maintenance in a single population. This work advances our understanding of how young sex chromosomes can follow non-linear paths of differentiation and degeneration.

## Results

### Whole genome sequencing of multiple Y haplotypes coexisting in a single pond

To characterise the sex-chromosome differentiation at high resolution and understand the coexistence of multiple Y haplotypes in one population, we combined pool-seq of phenotypically and genetically defined (using 12-17 sex-linked markers (79, 84)) male and female pools, and whole-genome sequencing of doubled-haploid YY individuals generated by androgenesis. We generated five pools: two fully differentiated male pools (top two most frequent haplotypes Y_B1_^a^, Y ^a^), one semi-differentiated male pool (Y ^0^), one undifferentiated male pool (XX males), and one XX female pool (Table1; Supplementary Table S1). Pool sizes ranged from 48 to 120 individuals (Table 1). To obtain further insights from Y-specific features and avoid the challenges of phasing X and Y sequences of homomorphic sex chromosomes, we further sequenced six doubled-haploid YY individuals, which were generated with an androgenesis approach (Supplementary Table S2 and S3, see details in M&M). We sequenced six doubled haploid YY individuals (Table S2, S3): two fully differentiated Y (Y ^a^) and four semi-differentiated Y (Y_B1_ ^0^, Y ^0^) (Table 1), as well as mate-pair sequencing libraries of the same fully differentiated YY male (Y ^a^).

**Table 1.** Phenotypic and genotypic information on samples of pool-seq and doubled haploid YY individuals. Doubled haploid individuals refers to the YY individuals generated through androgenesis. Family number refers to the sequenced YY individual offspring generated from a given clutch from Meitreile in the same year. The frequency of each male genotype was calculated from (84), based on sampling 842 males in Meitreile. Y differentiation level was inferred from 12-17 sex-linked markers (79, 84).

| Sex | Y haplotype | Frequency of each male genotype | Y differentiation | Pool-seq |  | WGS of YY individuals |  |
| --- | --- | --- | --- | --- | --- | --- | --- |
|  |  |  |  | Pool size | Coverage | Family # | Coverage |
| Female | NA | NA | NA | 120 | 197.10 | NA | NA |
| Male | X | 40.6% | Undifferentiated | 120 | 147.81 | NA | NA |
| Male | $Y_{B1}^0$ | 6.5% | Semi-differentiated | NA | NA | 40 | 15.40 |
|  |  |  |  |  |  | 46 | 11.87 |
| Male | $Y_{B1}^a$ | 21.4% | Fully differentiated | 120 | 146.88 | 18 | 12.87 |
|  |  |  |  |  |  | 50 | 12.30 |
| Male | $Y_{B2}^0$ | 7.8% | Semi-differentiated | 50 | 108.08 | 33 | 11.20 |
|  |  |  |  |  |  | 4 | 12.76 |
| Male | $Y_{B2}^a$ | 11.6% | Fully differentiated | 48 | 145.37 | NA | NA |

The five pools yielded 548.8 - 744.1Gb of sequence data (Supplementary Table S4), corresponding to 1.2 - 3.0x coverage per individual, and the doubled-haploid YY individuals yielded 46.0 – 63.3 Gb per genome and mate-pair libraries with 1.50 – 21.1G bp (Table 1; Supplementary Table S4). Reads were mapped to the chromosome-level female reference genome (85), in which 99.5% of the 4.11-Gb assembly is assigned to 13 chromosomes and has 90.7% BUSCO completeness (86). Most previously published sex-linked markers mapped a region spanning approximately two-thirds of the sex chromosome (chromosome 1), overall confirming the correspondence between earlier marker-defined Y haplotypes and the chromosome-scale assembly used here (Supplementary Table S5). One marker’s (Bfg191) order and position were sightly changed in the assembly (*Kank1, Dmrt1, Dmrt3, Bfg191,* compared to the published genetic map of *Kank1*, *Bfg191*, *Dmrt1* and *Dmrt3*). The observed minor discrepancy may reflect population-specific variation or local structural differences between the Swiss samples and the UK reference genome.

### Genome-wide scans reveal three discrete levels of sex chromosome differentiation

We first assessed X-Y differentiation by comparing each male pool with the XX female pool using male-to-female read-depth ratio, SNP-density ratio, nucleotide diversity and F_ST_. We use X-Y differentiation to refer broadly to SNP accumulation, allele-frequency differences, indels, and repeat accumulation, whereas we use X-Y divergence/degeneration to refer to differentiation responsible for reduced Y mapping and sequence loss, including in genes. In young or largely homomorphic systems, X and Y sequences are still similar enough that male X- and Y-derived reads map equally well to the X reference; therefore, sex linkage is mainly detected through elevated male heterozygosity, SNP density, and F_ST_ (87, 88). In older or more degenerated regions, Y-linked reads map less efficiently to the X reference because of accumulated divergence or sequence loss, producing reduced male-to-female read-depth ratios (88). Under substantial Y-linked sequence loss, the male-to-female read-depth ratio is expected to approach a log_2_ ratio of –1, corresponding to approximately half the female coverage (89).

We found no read-depth ratio difference on autosomes in any male-female pool comparison (Supplementary Fig. S1). On the sex chromosome, XX males and Y_B2_^0^ males did not deviate from the autosomal 95% confidence interval (CI), indicating no detectable large-scale gene loss or heteromorphic regions (Fig. 2a1, a2). The fully differentiated Y_B1_^a^ and Y_B2_^a^ male pools showed only modest, localized reductions in coverage compared to the female pool, with log2 ratios near -0.05 rather than the expected -1 under hemizygous gene loss (Fig. 2a3, a4). Consistently, genes with zero mapped reads, a proxy for gene loss, in double-haploid Y_B1_^a^ genomes were more frequent on autosomes than on the sex chromosome (Fisher’s exact tests: *P* < 0.001; Supplementary Table S6) across the length of which they were randomly distributed (Supplementary Fig. S2). Given the modest and uneven coverage of these samples, 9.2–13×, and the use of a UK reference genome for Swiss resequencing samples, such zero-read genes are likely dominated by stochastic coverage variation, mapping ability differences, and possible population-specific sequence or structural variation rather than true Y-linked gene loss. Together with the read-depth ratio results, none of the Y haplotypes shows evidence of extensive Y-linked gene loss; instead, read-depth ratio reduction is limited and broadly consistent with the degree of Y differentiation.

**Fig. 2.**
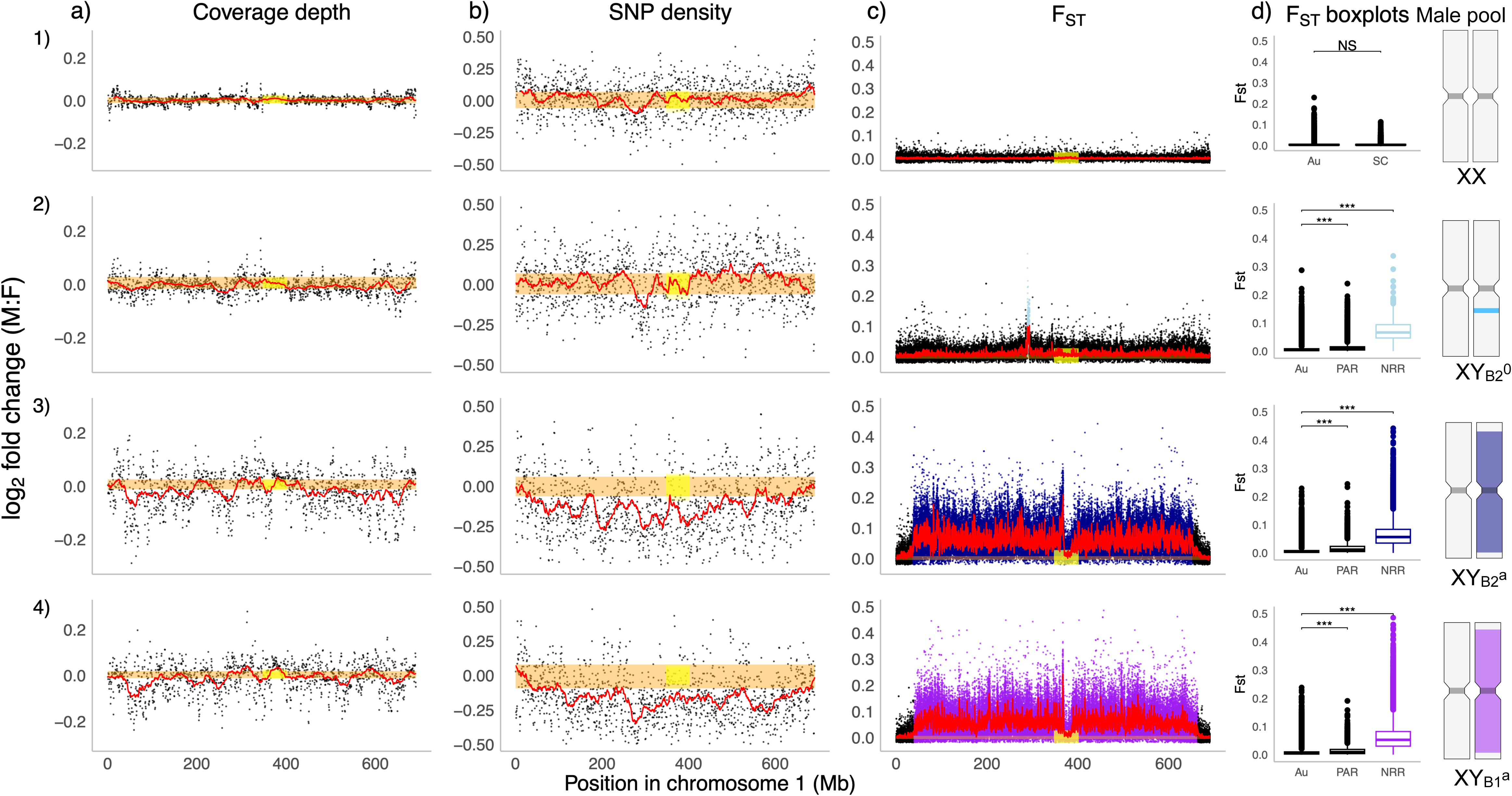
Genomic signatures of sex-chromosome differentiation across male pools. Read-coverage ratio (a), SNP-density ratio (b), F_ST_ across the sex chromosome (c), and F_ST_ boxplots across genomic compartment (autosome, PAR, NRR) (d) are shown for four male pools relative to the female pool: an undifferentiated XX male pool (1), a semi-differentiated Y_B2_ ^0^ pool (2), and two fully differentiated Y_B2_^a^ (3) and Y ^a^ (4) pools. The red horizontal line denotes the rolling mean (window size = 50) along the sex chromosome, and the orange band marks the central 95% of autosomal rolling-mean values. Yellow shaded regions indicate putative centromeres. For panels (a) and (b), each point represents the male: female ratio of mean read coverage or SNP density calculated in 1-Mb windows with 500-kb step size. For panel (c), F_ST_ values are averaged within 10-kb windows with 5-kb step size. Au: autosome, PAR: pseudoautosome region, NRR: non-recombining region.

Strong population genetic signal of sex linkage was on SNP-density ratio and nucleotide diversity. Suppressed recombination is expected to increase X-Y divergence leading to increased male heterozygosity (89). In pooled data, however, SNP density reflects allele frequencies within the pool and does not simply measure pairwise X-Y divergence. If Y haplotypes are largely shared within a male pool, Y-linked variants contribute little to segregating SNP density, whereas the female pool contains two sets of recombining X chromosomes and can retain higher X-linked polymorphism. Thus, regions with strong Y differentiation may show limited nucleotide diversity in XY pools compare to XX pools, and SNP-density ratio is expected to be higher in XX pools. Consistently with this expectation, π on the sex chromosome was significantly reduced in the fully differentiated Y_B1_^a^ pool, followed by Y_B2_^a^ and then the semi-differentiated Y_B2_^0^ pool, relative to the XX female pool (Supplementary Table S7; Fig. S3a; all *P* < 0.0001). In contrast, the XX male pool showed only a negligible effect despite statistical significance (Cohen’s *d* =0.007), with π values nearly identical to the XX female pool (Supplementary Fig. S3a,c). Autosomal comparisons on π showed no dramatic differences across male pools, as expected (Supplementary Fig. S3b,c; Table S7). SNP density ratio did not differ across autosomes in any pool comparison (Supplementary Fig. S4), nor along the sex chromosome in the XX male versus XX female pool comparison (Fig. 2b1). The semi-differentiated Y_B2_^0^ pool also remained within the autosomal 95% CI (Fig. 2b2). By contrast, the fully differentiated Y_B1_^a^ and Y_B2_^a^ pools had a significant female-biased SNP-density ratio across most of the sex chromosome, except for the chromosome ends (Fig. 2b3, b4), consistent with the strongest π reduction of the fully differentiated Y pools.

Window-based F_ST_ provided the clearest definition of the nonrecombining regions (NRRs). F_ST_ was near zero and homogeneous across the 12 autosomes in all comparisons (median: 0.00104 – 0.00167; Supplementary Fig. S5). Consistently, comparison of the XX male and female pools showed no significant F_ST_ elevation on the sex chromosome relative to autosomes (permutation test *P* = 0.054, Fig. 2c1, d1). In contrast, the semi-differentiated Y_B2_^0^ pool showed a single narrow F_ST_ peak of 4.64 Mb, approximately 0.67% of the chromosome, which we define as the small NRR (median F_ST_ = 0.0663; permutation test *P* = 0.001; Fig. 2c2, d2). The remainder of the sex chromosome had low F_ST_, although slightly above the autosomal background (median = 0.00656 versus 0.00153; permutation test *P* = 0.001), consistent with a recombining pseudo-autosomal region (PAR). In the fully differentiated Y_B1_^a^ or Y_B2_^a^ pools, F_ST_ was elevated across most of the sex chromosome (median = 0.0511 and 0.0558, respectively), except near both chromosome ends, and was significantly higher than autosomes in both comparisons (permutation test, *P* = 0.001; Fig. 2c3, d3, c4, d4; Supplementary Table S8). Changepoint analysis identified broadly similar boundaries of these large NRRs in the two Y haplotypes: 39.91-664.61 Mb for Y_B1_^a^ and 36,39 Mb-654.10 Mb for Y_B2_^a^. The terminal PARs of the fully differentiated haplotypes also showed slightly elevated F_ST_ relative to autosomes (median = 0.00241 and 0.00645; permutation test, *P* = 0.001 respectively) but with much smaller differences. Together with the read-depth ratio, nucleotide diversity, and SNP-density ratio analyses, these results confirm three distinct levels of sex chromosome differentiation within the same pond: no detectable NRR in XX males, a 4.64 Mb small NRR (0.65%) in Y_B2_^0^, and chromosome-scale large NRRs (∼90%) in Y_B1_^a^ and Y_B2_^a^.

**Fig. 3.**
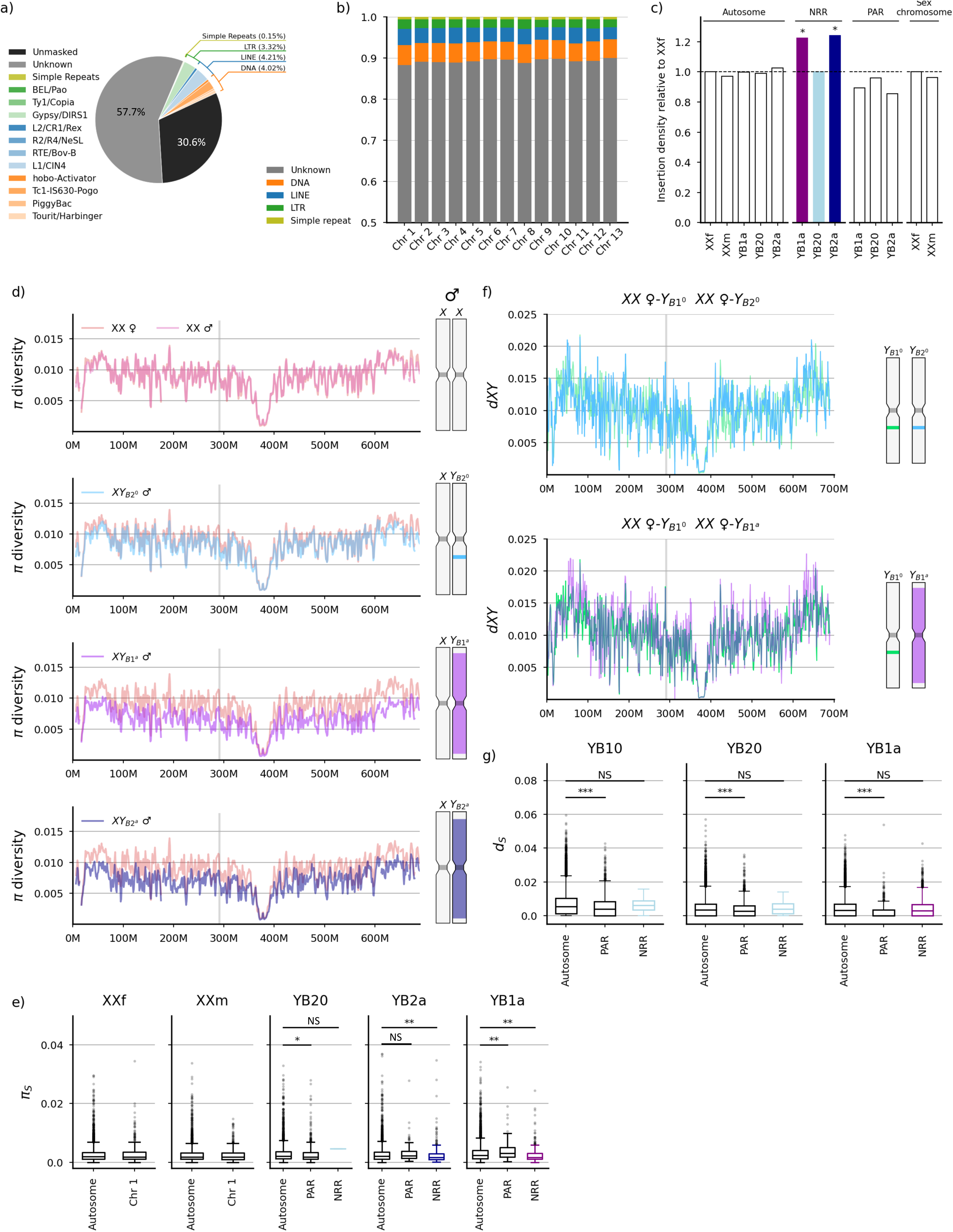
Sequence divergence and degeneration of the sex chromosome. (a) Proportion of major transposable element (TE) classes and simple repeats in the reference genome, based on a *de novo* species-specific TE library. (b) Proportion of major TE classes across the 13 chromosomes. (c) Relative TE insertion rates for the 4 male pools comparing to the female pool. (d) Rolling mean of nucleotide diversity π between each of the four male pool and the female pool. (e) π_S_ comparison across different genomic compartments within each Y haplotypes. (f) Pairwise sequence divergence (Dxy) along the sex chromosome between the reference X chromosome of two semi-differentiated Y haplotypes (Y_B1_ ^0^, Y ^0^) and one fully differentiated Y haplotype (Y ^a^). (g) d_S_ comparison across different genomic compartments (A: autosome, PAR, and NRR) within each Y haplotypes. Grey shading denotes the small NRR.

**Fig. 4.**
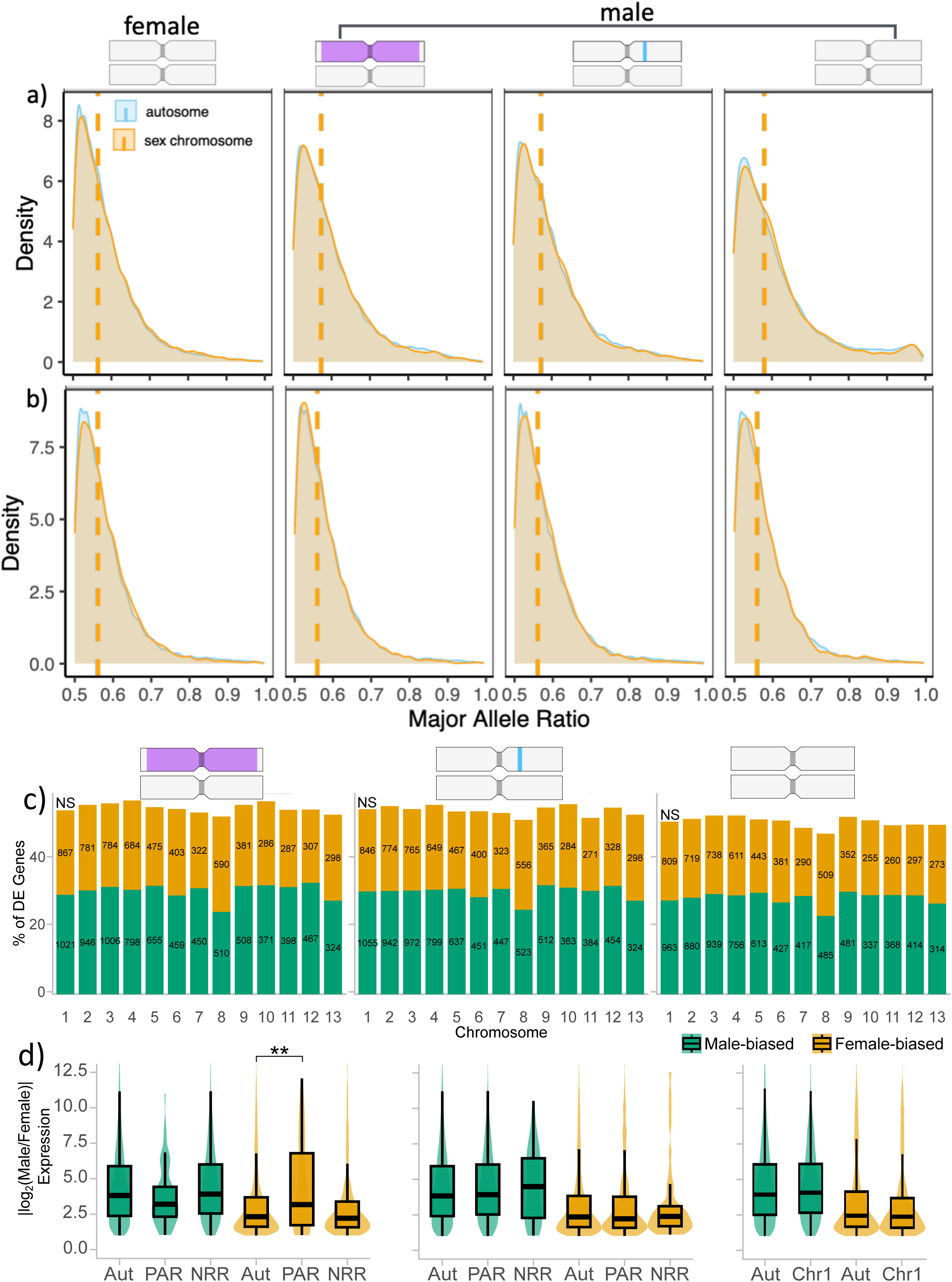
Distributions of major allele ratios and sex-biased gene expression across males with three distinct levels of Y-chromosome differentiation. , illustrated by the cartoon where purple indicates a large NRR and blue a small NRR, ordered from fully-differentiated Y to semi-differentiated Y, to undifferentiated Y. Density plots show the distribution of major allele ratio (MAR) for autosomal (blue) and sex-chromosomal (orange) genes in gonad (a) and brain (b) tissues of females and males. Vertical dotted lines denote median MAR values. (c) Proportion of female-, and male-biased genes per chromosome among three types of males. (d) Boxplot of differential gene expression (in log2(male/female)) among different genomic compartments among three types of males (Aut: autosome, NRR: non-recombining region and Chr1: sex chromosome).

### Large NRRs show high repeat accumulation but limited coding-sequence degeneration

To further assess levels of sex chromosome divergence and degeneration, we quantified TE insertions, nucleotide and synonymous/nonsynonymous diversity (π, π_N_, π_S_) using pool-seq data and estimated pairwise sequence and synonymous/nonsynonymous divergence (Dxy, d_N_ and d_S_) using doubled haploid YY genomic datasets. A *de novo* TE library built from the XX reference genome showed repeats comprising 69.4% of the genome, with most (80.9%) elements unclassified, while Kimura divergence indicated a recent burst of TE activity (Fig. 3a; Supplementary Fig. S6). However, no specific TE class was enriched on the X chromosome relative to autosomes (Fig. 3b). Using PopoolationTE2, we found a significantly elevated TE insertion density in the large NRRs of both Y_B1_^a^ and Y_B2_^a^ pools relative to autosomes (Glm, *P* < 0.003), but not in the NRR of the Y_B2_^0^ pool (*P* = 0.504, Fig. 3b). As expected, there was no significant increase in male-specific TE insertion density across autosomes among male pools (Glm, *P* = 0.524; Fig. 3c).

Patterns of nucleotide and coding-sequence variation indicated limited functional degeneration. Nucleotide diversity (π) was reduced across the large NRRs of Y_B1_^a^ (Cohen’s *d* = -0.708) and Y_B2_^a^ (*d* = -0.581), whereas XX males (*d* = 0.007) resembled XX females and Y_B2_^0^ (*d* = -0.189) showed only a localized reduction around the small NRR (Fig. 3d; Supplementary Fig. S3). Consistently with this pattern, synonymous diversity (π_S_) was significantly reduced on the NRRs of Y_B1_^a^ and Y_B2_^a^ male pools (permutation test *P* < 0.0035, Supplementary Table S9a), but π_S_ did not differ between sex chromosome compartments and autosomes in the XX male or Y_B2_^0^ male pools (permutation test, all *P* > 0.23, Fig. 3e, Supplementary Table S9a, Fig. S7). π_N_/π_S_ did not differ between sex chromosome compartments and autosomes in any male pool (permutation test, all *P* > 0.29, Supplementary Table S9b, Fig. S8-10).

Doubled-haploid YY genomes provided an independent test of direct pairwise sequence divergence between Y haplotypes and the X reference. Absolute sequence divergence (Dxy) showed elevated nucleotide divergence for the large NRR of Y_B1_^a^, and locally elevated divergence at the small NRRs of Y_B1_^0^ and Y_B2_^0^ (Fig. 3f). Consistently, autosomal d_S_ values were uniformly low (median = 0.0034-0.0055, Supplementary Table S10a, Fig. S11). d_S_ values on the sex chromosome NRRs were similarly low (median = 0.0026-0.0061), remained within the autosomal 95% CI, and did not differ significantly from autosomes in Y_B1_^0^, Y_B2_^0^ or Y_B1_^a^ (permutation test, *P* = 0.79, 0.79, 0.10 respectively; Fig. 3g). Changepoint analysis of d_S_ identified no additional internal strata along the large and small NRR respectively. Similarly, d_N_/d_S_ did not differ significantly between sex chromosome compartments and autosomes (permutation test, all *P >* 0.82; Supplementary Fig. S12-13, Table S10b), except for an elevated value in the PAR of fully differentiated Y_B1_^a^ (permutation test, *P* = 0.0006), which was not located in the NRR and was not shared across haplotypes. Thus, we find evidence for moderate pairwise sequence divergence between X and Y for NRRs, but these likely involve non-coding regions, as there was no evidence for elevated nonsynonymous divergence or relaxed purifying selection in the NRRs. These results indicate that the large NRRs of the differentiated Y haplotypes have, compared to the X chromosome, accumulated significantly more TE insertions, show reduced nucleotide diversity and moderate sequence divergence, but they have no detectable gene loss, no faster sex-linked gene evolution and no evidence of relaxed purifying selection.

### Differentiated Y haplotypes have no expression decay or sex-biased gene enrichment

To test whether Y differentiation is associated with reduced Y-linked transcription, we estimated the major-allele ratio (MAR) for expressed heterozygous sites using gonad and brain RNA-seq data from Meitreile. If Y-linked alleles were broadly transcriptionally degraded, MAR should increase on sex-linked genes relative to autosomal genes. Males carrying different Y haplotypes showed no significant elevation of sex-linked MAR relative to autosomes in gonad (median= 0.58-0.59, permutation test, all *P* > 0.20), or brain (permutation test, all *P* > 0.21, 0.56-0.57) (Fig. 4a, b), and rolling-mean MAR profiles were similar across females and all male genotypes (Supplementary Fig. S14a, b). Pairwise comparisons among male genotypes detected almost no differentially expressed genes in either brain or testis (|log_2_FC| >= 1, FDR < 0.05; Supplementary Fig. S15-16, Table S11). As expected, females showed no difference between autosomal and sex-linked MAR in either gonad or brain (permutation test, all *P* = 0.15 and 0.13 respectively, Fig. 4a, b). These results provide no evidence for Y-linked expression decay in any Y haplotype.

We next tested whether sex-biased genes are enriched on the NRRs across different Y haplotypes. Brains showed only 1 sex-biased gene, located on Chr13 (an autosome), so enrichment analyses focused on gonadal tissues. Female-versus-male comparisons with XY_B_^a^, XY_B_^0^ and XX males identified 6598, 6449 and 6057 female-biased genes, and 8106, 8032, and 7554 male-biased genes respectively (|log_2_FC| ≥ 1, FDR < 0.05; Supplementary Fig. S17). Across these contrasts, neither the magnitude of sex-biased expression nor the number of sex-biased genes differed between the sex chromosome and autosomes, except for one case described below (permutation test, all *P* > 0.3; gene number permutation test, all *P* > 0.33; Fig. 4c; Supplementary Fig. S18, S19; Supplementary Table S12-17). The same result held when the NRRs and PARs were analysed separately in semi- and fully differentiated males (permutation test, all *P* > 0.12; Supplementary Fig. S19, Table S16, S17) for all contrasts.

The only exception was stronger female-biased expression in the PAR of XY_B_^a^ male testes relative to autosomes (*P* = 0.00045), although the number of female-biased genes was not increased (*P* = 0.120). Because the PAR lies largely in gene-rich telomeric regions and shows reduced π and πs (Fig. 3g, 4d), we tested whether this signal reflected a broader telomeric pattern. Indeed, similar enrichment of female-biased expression magnitude occurred in distal regions of autosomes 3, 6 and 7 in both fully and semi-differentiated males (permutation test, all *P* < 0.005). Applying the Y_B_^a^ - defined PAR boundaries to all male genotypes also recovered significantly higher female-biased expression magnitude in this region in XY_B_^a^, XY_B_^0^, XX males (all P < 0.005; Supplementary Table S18). We therefore interpret the PAR signal as a feature of distal, gene-rich telomeric regions rather than evidence for NRR-specific sexualization in terms of gene expression.

### *Dmrt1* is the strongest candidate sex-determining gene

The narrow 4.64 Mb small NRR in Y_B2_^0^ provided a high-resolution entry point for identifying the candidate master sex-determining gene. Changepoint analysis on F_ST_ values separated this interval into a 3.25 Mb highly differentiated stratum containing 19 annotated genes, including *Dmrt1*, and a 1.39 Mb intermediate stratum with 10 annotated genes (Wilcoxon test, *P* < 8.4 × 10-11; Fig. 5a, 5b; Supplementary Table S19). Among these 29 genes, *Dmrt1* and *Spata6l* have known roles in sex determination/testis development and spermatogenesis, respectively (Supplementary Table S19; Fig. 5b). Adult RNA-seq from Meitreile further identified two genes with almost exclusive gonadal-specific expression in the region: *Dmrt1*, which showed testis-specific expression, and *LOC120943612*, an ovary-specific ortholog of anuran class-A G protein-coupled receptors, potentially involved in steroidogenesis and ovarian signalling (Fig. 5a, b, c). *Spata6l*, however, did not show significant gene expression between sexes (Fig. 5b). Because *Dmrt1* is a deeply conserved regulator of male cell fate and testis development across vertebrates, its location within the most strongly differentiated stratum and its testis-specific expression make it the strongest functional candidate in the region. Published developmental RNA-seq from whole tadpoles in other *R. temporaria* populations (Ammarnäs with fully differentiated XY, Tvedöra with semi-differentiated XY^0^) (76, 90), showed low or undetectable *Dmrt1* and *LOC120943612* expression across early development, although the latter was strongly female-biased at the froglet stage (Supplementary Fig. S20, S21). Because whole-tadpole transcriptomes dilute the (very tiny) gonadal signal present at early development (91), these data do not exclude an early sex-determining role for *Dmrt1.* Targeted gonadal transcriptomics during the thermosensitive and early sex-determining period are required to confirm a role of *Dmrt1* in sex determination during early development.

**Fig. 5.**
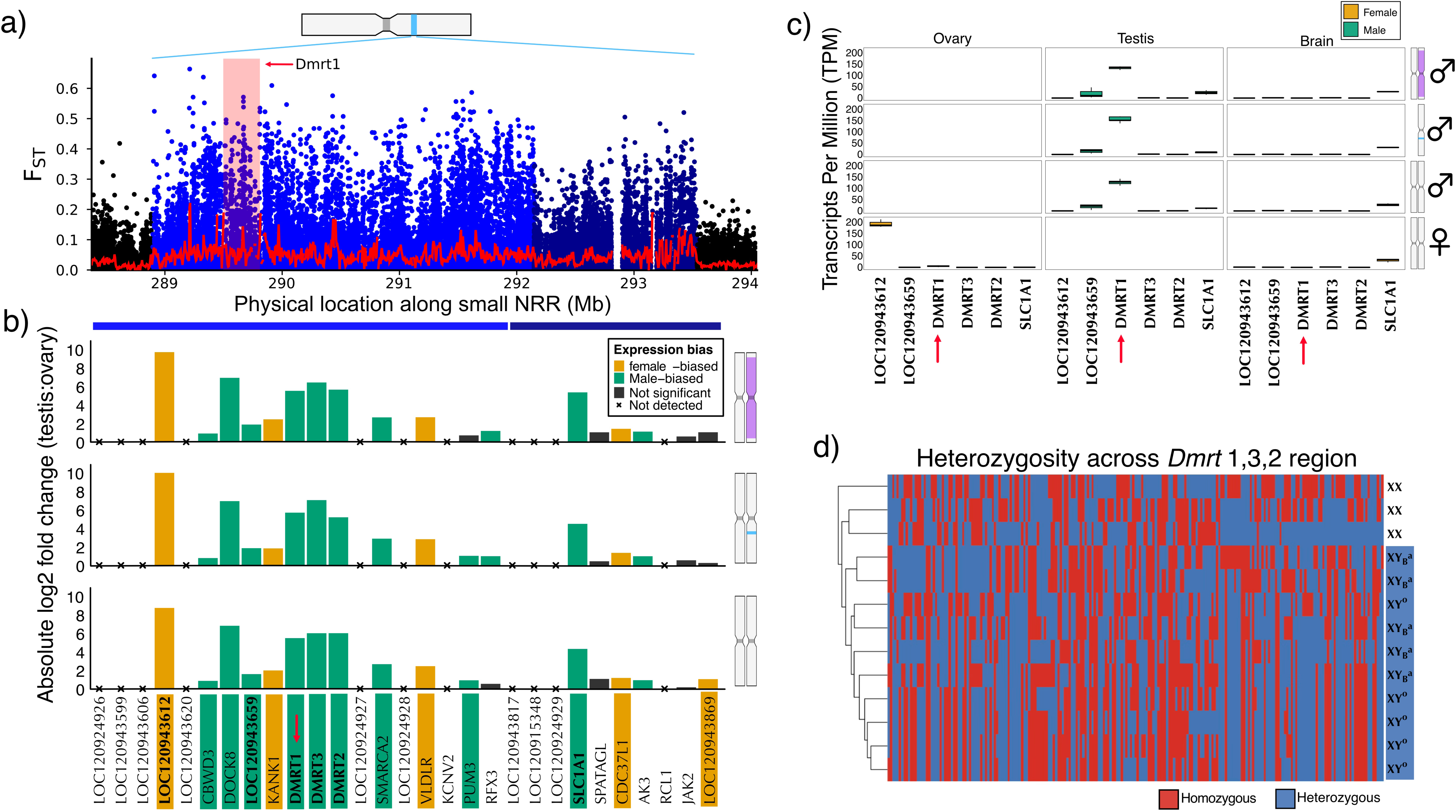
Sex-biased gene expression and RNA-seq SNP clustering within the small NRR. (a) The elevated F_ST_ peak between males carrying the semi-differentiated Y ^0^ haplotype and females is subdivided into two strata, a light-blue and a dark-blue region. (b) Differential expression analyses identify male- and female-specific or biased genes in adult testis versus ovary across three male types representing distinct stages of Y-chromosome differentiation. (c) Expression profiles of the top six candidate genes, including *Dmrt1,* display pronounced sex-specific or sex-biased expression. (d) A clustering dendrogram based on homozygosity in XX individuals and heterozygosity in XY individuals including both fully and semi-differentiated Y haplotypes was constructed from SNPs derived from the RNA-seq datasets.

Genotype patterns across expressed SNPs provided independent support for *Dmrt1* as the best candidate master sex-determining gene. Hierarchical clustering of SNPs from Meitreile gonadal RNA-seq reads revealed male-specific clusters only for *Dmrt1, Dmrt3,* and *Dmrt2* among genes in the small NRR (or sex-determining region) (Fig. 5d). The strongest signal occurred around *Dmrt1-Dmrt3*: 21/25 SNPs homozygous in XX males but heterozygous in both XY_B_^a^, XY_B_^0^ males localised in *Dmrt1* or between *Dmrt1* and *Dmrt3* (Supplementary Tables S20, S21). A stricter scan for SNPs homozygous in all XX individuals but heterozygous in both semi- and fully differentiated XY_B_^a^ males identified one SNP located between *Kank1* and *Dmrt1* (Supplementary Table S22). Together, the combined data on the highest male-female F_ST_ peak, candidate gene function, testis-specific expression, and XY-specific heterozygosity identify the *Dmrt1*-linked region as the most plausible sex-determining region, with *Dmrt1* as the leading candidate master sex-determining gene.

### The *Dmrt1*-linked region is shared among all Y haplotypes, whereas larger NRRs are more recent and haplotype-specific

We next tested whether coexisting multiple Y haplotypes represent independent origins of sex determination, a linear differentiation trajectory, or shared ancestry with subsequent haplotype-specific differentiation. To do so, we compared XX, XY_B2_^0^, XY_B1_^a^ or XY_B2_^a^ male pools using pairwise F_ST_, nucleotide diversity π, pairwise divergence Dxy from the doubled haploid YY data, and SNP clustering from RNA-seq data. Autosomal F_ST_ among male pools was uniformly near zero, providing a baseline of minimal genome-wide differentiation (Supplementary Fig. S22). On the sex chromosome, F_ST_ between XX male and Y_B1_^a^ or Y_B2_^a^ pools showed consistently elevated F_ST_ across the large NRRs, including the 4.64 Mb *Dmrt1*-linked region (Wilcoxon test, FDR-corrected *P* < 0.0001; Fig. 6a). The Y_B2_^0^ versus XX male comparison showed a sharp F_ST_ peak restricted to the *Dmrt1*-linked region (*P* < 0.0001; Fig. 6a). Thus, all Y haplotypes differ from the X in the *Dmrt1*-linked region, whereas only Y_B1_^a^ or Y_B2_^a^ show broad differentiation across most of the sex chromosome (Fig. 2c, Fig. 6a). Nucleotide diversity along the sex chromosome showed the same contrast: XX and Y_B2_^0^ males were nearly identical across most of the sex chromosome, except near and within the *Dmrt1*-linked region (Fig. 6d). In sharp contrast, along the large NRR, π of both Y_B1_^a^ and Y_B2_^a^ was dramatically reduced (Fig. 6d).

**Fig. 6.**
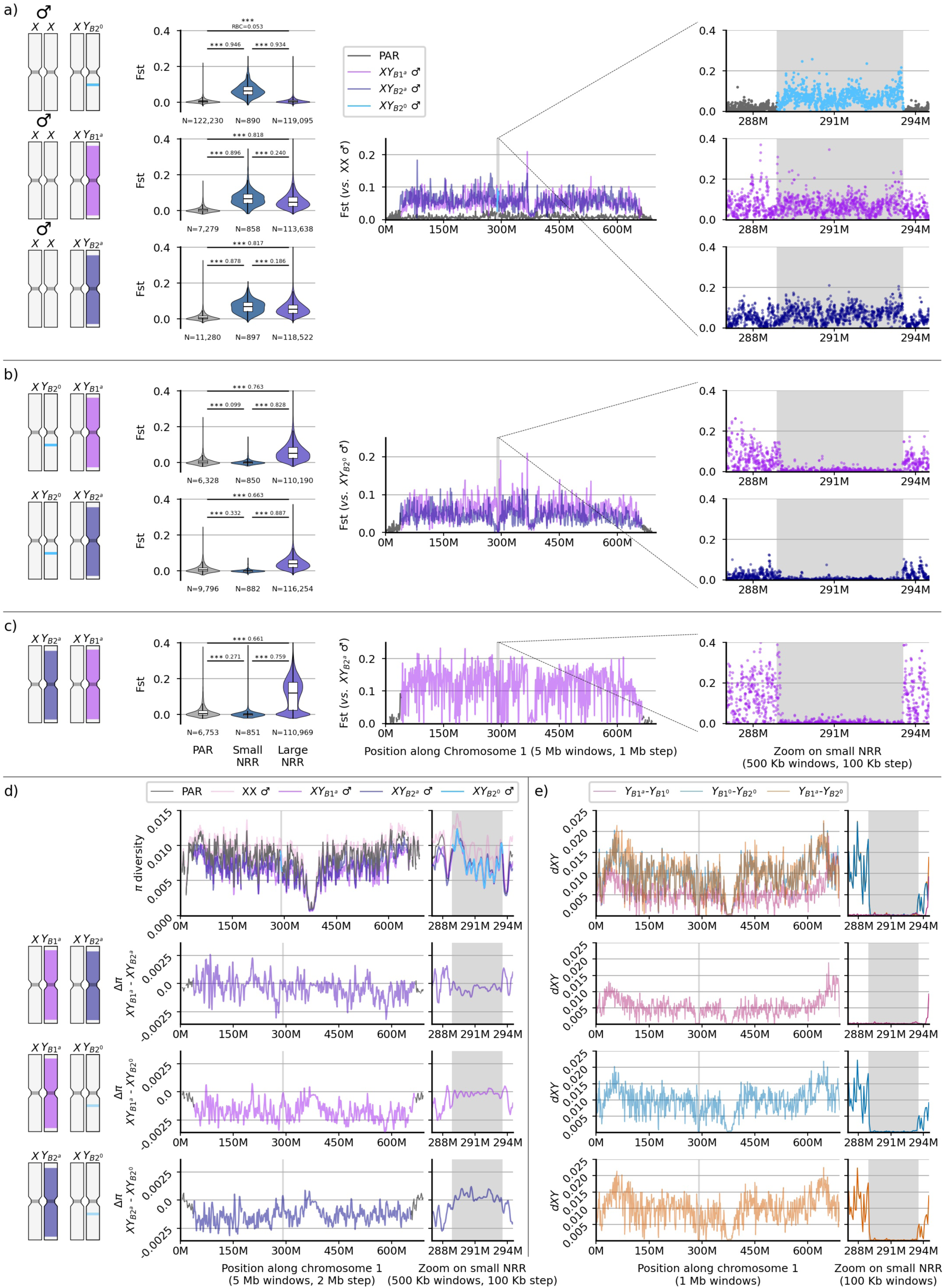
Genomic divergence reveals a shared sex-determining region across various Y haplotypes. (a–c), Windowed F_ST_ across sex chromosome for pairwise comparisons among XX individuals and males carrying the Y_B2_ ^0^, Y_B1_^a^ or Y ^a^ haplotypes. Violin plots summarize differentiation in the PAR, small NRR and large NRR, and right panels show zoomed views of the small NRR (4.64 Mb *Dmrt1*-linked region). (d) Nucleotide diversity (π) and pairwise Δπ profiles highlight haplotype-specific reductions in diversity across the NRR regions across all 4 male pools. (e) Pairwise sequence divergence (Dxy) among Y haplotypes along sex chromosome and zoomed in view of the small NRR. Grey shading denotes the small NRR; window sizes are indicated below each panel.

Pairwise comparisons among Y-bearing male pools revealed that the 4.64 Mb *Dmrt1*-linked region is shared across all Y haplotypes, whereas large NRRs accumulated mutations independently. Across the 4.64 Mb region, F_ST_ among XY_B2_^0^, XY_B1_^a^ and XY_B2_^a^ male pools was close to zero and significantly lower than in the PARs or the rest of the large NRRs (all FDR-corrected *P* > 0.29; Fig. 6b, c; Supplementary Fig. S23). The boundaries of this low-F_ST_ region matched the small NRR detected in the Y_B2_^0^-versus-female comparison and encompassed strata defined by changepoint analysis (Fig. 2c2; Fig. 6a, b, c). Outside the *Dmrt1*-linked region but within the large NRR, differentiation among Y haplotypes increased markedly and was heterogeneous: F_ST_ was highest between Y_B1_^a^ and Y_B2_^a^ (Fig. 6c), intermediate between Y_B1_^a^ and Y_B2_^0^ (Fig. 6b), and lowest between Y_B2_^a^ and Y_B2_^0^ (Fig. 6b; all FDR-corrected *P* < 0.0001).

Patterns of nucleotide diversity and pairwise sequence divergence independently supported a shared sex-determining region. Across most of the sex chromosome, π differed among XX males and the three Y-bearing pools, consistent with haplotype-specific differentiation outside the sex-determining region. Within the 4.64-Mb *Dmrt1*-linked region, however, π was nearly identical among Y_B1_^0^, Y_B2_^0^ and Y_B1_^a^ pools, and pairwise π differences approached zero (Fig. 6d). Similarly, pairwise D_XY_ varied among doubled-haploid YY genomes across the broader sex chromosome but dropped to zero for all Y–Y comparisons within the 4.64 Mb region (Fig. 6e). Notably, Y_B1_^0^ and Y_B1_^a^, which share the *Dmrt1–Dmrt3* marker haplotype, also showed markedly reduced divergence across the large NRR relative to other Y–Y comparisons (i.e. largely 0-0.01 vs 0.01-0.02; Fig. 6e). This pattern provides direct evidence that Y_B1_^0^ and Y_B1_^a^ derive from the same ancestral Y background, with the semi-differentiated Y_B1_^0^ haplotype likely resulting from erosion of a formerly differentiated Y through sex-reversal-mediated X–Y recombination.

As an independent test of the shared sex-determining region, we performed sliding-window RNA-seq-based SNP clustering that followed the expected pattern of XX-specific homozygosity and XY heterozygosity (Fig. 7; Supplementary Figs. S24–S28). In PAR1 and PAR2, clustering was not based on sex, consistent with ongoing recombination. Across the large NRR, fully differentiated Y males generally clustered together (Fig. 7: tree #1-#5, and #9-#15), as expected from their recombination suppression, whereas XX males grouped with females. Strikingly, only in the *Dmrt1–Dmrt3–Dmrt2* region did all Y-bearing males cluster together regardless of Y haplotype (Fig. 7: tree #6-#8). Together, the male–male F_ST_, π, D_XY_, and SNP clustering show that all Y haplotypes share the *Dmrt1*-linked region, the broader large NRRs have record haplotype-specific histories of divergence, erosion, and re-differentiation.

**Fig. 7.**
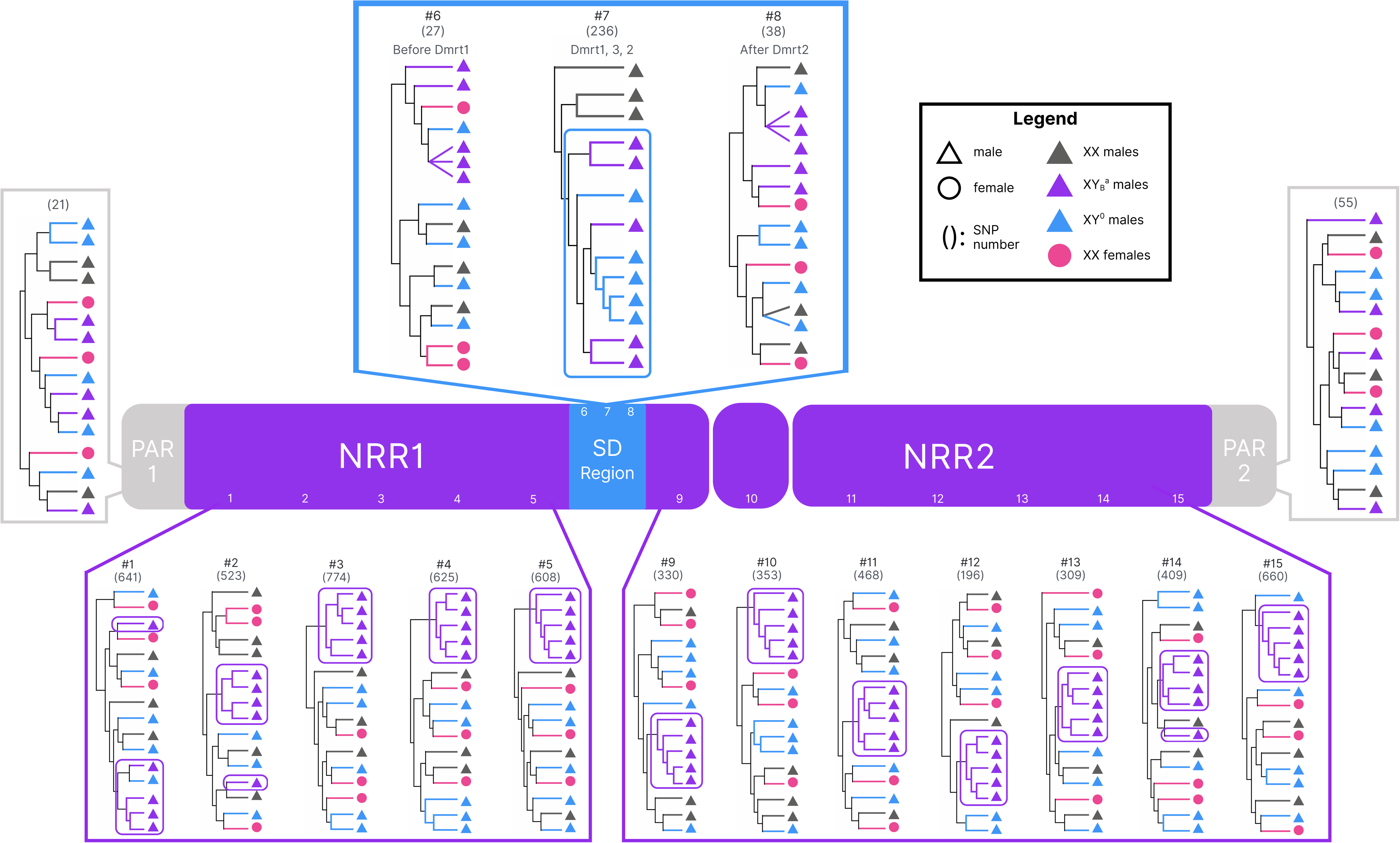
Clustering dendrograms based on SNPs. derived from RNA-seq read mappings of male and female gonad and brain tissues to the reference genome, illustrating patterns of homozygosity and heterozygosity across discrete compartments of the sex chromosome (PAR1, large NRR1, sex-determining (SD) region, large NRR2 and PAR2). Each dendrogram was generated from SNP sets extracted using a 10-Mb sliding window across PAR1 and PAR2, a 50-Mb sliding window across the NRR, and three subdivided windows spanning small NRR. Number of SNPs used in each clustering analysis is indicated in parentheses. Dendrograms are numbered and ordered according to their relative genomic positions along the sex chromosome.

We then estimated relative ages of the shared *Dmrt1*-linked region and the broad large NRRs using synonymous divergence (d_S_) from doubled-haploid YY sequences and a published mutation-rate estimate calibrated from *Nanorana parkeri* (92) (Table 2). The shared *Dmrt1*-linked region was estimated to have originated approximately 3.47–5.70 Mya. In contrast, d_S_-based estimates for the larger NRRs were younger: 2.94–3.32 Mya for Y_B1_^a^ and 0.82–1.08 Mya for Y_B2_^0^. Together, these results indicate that the *Dmrt1*-linked region predates the expansion of the larger NRRs.

**Table 2.** Estimated age of small and large NRR regions reflected by the highly elevated F_ST_ region from Fig.2. The large NRR was subdivided into small NRR and the rest of large NRR. The age estimation and the confidence interval was based on the d_S_ between the YY doubled haploid males with Y_B2_^0^ or Y_B1_^a^ haplotypes and the reference X chromosome sequence. Mutation rate per synonymous site per year (0.776 × 10^−9^) was based on the Tibetan frog *Nanorana parkeri* (92).

| Y haplotype | Small NRR | $d_s$ (mean ± se) | Age (Myr) | Age with 95%CI | Rest of large NRR | $d_s$ (mean ± se) | Age (Myr) | Age with 95%CI |
| --- | --- | --- | --- | --- | --- | --- | --- | --- |
| $Y_{B2}^0$ | 100% | $(26.9 \pm 7.6) \times 10^{-4}$ | 3.47 | 1.55-5.39 | 100% | $(7.4 \pm 0.5) \times 10^{-4}$ | 0.95 | 0.82-1.08 |
| $Y_{B1}^a$ | 100% | $(44.3 \pm 8.2) \times 10^{-4}$ | 5.70 | 3.62-7.78 | 100% | $(26.9 \pm 7.6) \times 10^{-4}$ | 3.13 | 2.94-3.32 |

Finally, our data support the Y-chromosome evolutionary history summarized in Fig. 8. The coexisting Y haplotypes do not reflect independent origins of sex determination or a single linear series of Y degeneration. Instead, they share an ancestral *Dmrt1*-linked sex-determining region, estimated at ∼3.47–5.70 Mya, while the larger sex-linked regions represent younger, divergent Y backgrounds with haplotype-specific histories of recombination suppression, mutation accumulation, and X–Y differentiation. In Fig. 8a, Y_B1_^a^ and Y_B2_^a^ provide two examples of independently differentiated Y haplotypes, shared the same ancestral sex-determining region, each carrying large NRRs spanning most of the sex chromosome. By contrast, Y_B1_^0^ and Y_B2_^0^ retain only the shared *Dmrt1*-linked region after erosion of broader X–Y divergence, likely through sex-reversal-mediated recombination. Fig. 8b summarizes the cyclic dynamics of Y differentiation. Under extreme male heterochiasmy, a newly established sex-determining locus can rapidly become associated with a large NRR. Sex-reversed XY females can then restore X–Y recombination and reduce divergence outside the sex-determining region, whereas subsequent transmission through males can again promote broad recombination suppression and renewed Y-specific differentiation. Thus, in *R. temporaria*, an older shared sex-determining region is surrounded by younger Y backgrounds that have accumulated private mutations independently, allowing multiple Y haplotypes with different NRR sizes, ages, and degrees of differentiation to coexist within a single population.

**Fig. 8.**
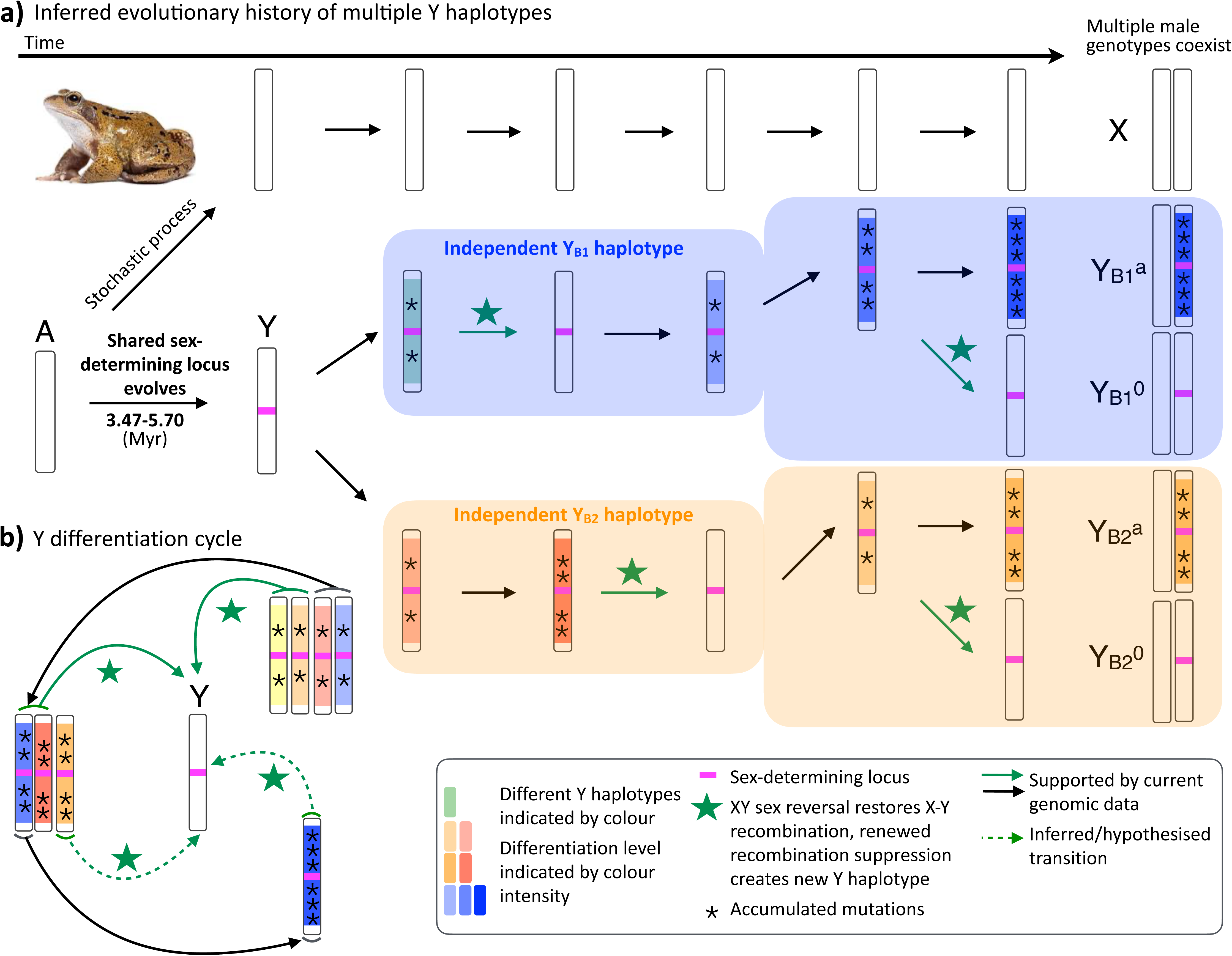
Proposed evolutionary history of multiple Y haplotypes. (a), Ancestral sex determination was likely stochastic, allowing XX males to arise in the absence of detectable sex-linked differentiation. A shared *Dmrt1*-linked small NRR then evolved approximately 3.47–5.70 Mya, establishing an ancestral Y chromosome. Subsequent, independent episodes of recombination suppression generated the Y_B1_^a^ and Y_B2_^a^ haplotype lineages, followed by progressive mutation accumulation and expansion of differentiated regions. Within each lineage, inferred XY sex-reversal events restored X–Y recombination and reset Y differentiation, giving rise to less differentiated haplotypes such as Y_B1_^0^ and Y_B2_^0^, whereas continued recombination suppression produced more differentiated haplotypes such as Y_B1_^a^ and Y_B2_^a^. This process results in the present-day coexistence of multiple male genotypes carrying Y haplotypes at different stages of differentiation. (b), Schematic of the proposed Y-differentiation cycle, in which recombination suppression, mutation accumulation, XY sex reversal and renewed suppression repeatedly generate new Y haplotypes. Chromosome colour indicates haplotype identity, and colour intensity indicates the degree of differentiation. Magenta marks the sex-determining locus, asterisks indicate accumulated mutations, and green stars denote inferred XY sex-reversal events. Solid arrows indicate transitions supported by genomic data; dashed arrows indicate hypothesised transitions.

## Discussion

Here, we use whole-genome sequencing to test how homomorphic sex chromosomes evolve under extreme heterochiasmy and recurrent X–Y recombination mediated by XY sex reversal. We ask whether coexisting Y haplotypes in a single pond represent independent sex-determining systems, successive steps in a single differentiation trajectory, or shared ancestry followed by haplotype-specific differentiation. Our results support shared ancestry with recurrent remodelling: all four Y haplotypes retain an ancient 4.64 Mb *Dmrt1*-linked sex-determining region with nearly identical present-day Y sequences, whereas the surrounding large NRRs in Y_B1_^a^ and Y_B2_^a^ are younger, haplotype-specific and span 618–625 Mb. This pattern shows that the age, size and degeneration of sex-linked regions can be uncoupled, supporting cyclic Y differentiation rather than a linear path toward irreversible Y degeneration. Independent Y haplotypes share a single ancestral *Dmrt1*-linked sex-determining region.

### Independent Y haplotypes share a single ancestral *Dmrt1*-linked sex-determining region

A central result of this study is the temporal decoupling between the sex-determining region and the larger NRRs. The shared *Dmrt1*-linked small NRR is estimated to have originated ∼3.47–5.70 Mya, broadly overlapping independent RAD-seq estimates for the origin of major Y-haplotype lineages across *R. temporaria* (Gradel et al., in prep). By contrast, d_S_-based estimates for the larger NRRs are younger, at ∼2.94–3.22 Mya for Y_B1_^a^ and ∼0.82–1.08 Mya for Y_B2_^0^. These absolute values should be interpreted cautiously, because young sex-linked regions may retain ancestral polymorphism, occasionally recombine through sex reversal, and vary locally in mutation and recombination history. Nevertheless, their relative order is informative: the *Dmrt1*-linked sex-determining region predates the broader expansions, and various evidence shows they are shared among all Y backgrounds.

This pattern is precisely the kind of architecture that recent work on young sex chromosomes predicts. Rather than representing a single monotonic progression from an undifferentiated chromosome to an increasingly degenerated Y, the *R. temporaria* Y chromosomes appear to record a layered history: an older shared region linked to sex determination, surrounded by younger peripheral regions that have expanded, persisted, or been eroded on different haplotype backgrounds. This interpretation aligns with work by (43, 44) showing that fully sex-linked regions may remain small, that partially sex-linked regions and PAR boundaries can be historically mobile, and that the extent of recombination suppression should not be inferred directly from either physical size or differentiation alone. Plant systems such as *Silene* have already shown that PAR boundaries can be polymorphic or shift over time (94); our data extend this logic to a vertebrate system in which multiple Y backgrounds co-occur within a single natural population.

The shared 4.64 Mb region also focuses attention on *Dmrt1* as the most plausible candidate master sex-determining gene. *Dmrt1* has been repeatedly recruited into sex-determining pathways across vertebrates, including fishes, birds, amphibians, and reptiles (95–99). In *R. temporaria*, the absence of fixed X–Y coding differences in *Dmrt1* suggests the sex-determining mechanism may depend on allelic diversification resulting in allele-specific expression between the X and Y copy during the thermally sensitive early embryonic sex-determining period. The observed adult near testis-restricted expression of *Dmrt1* is consistent with a role in male determination and testis development, while failure to detect its expression in whole-tadpole RNA-seq is not decisive due to tissue allometry issue because early gonads represent only a small fraction of whole tadpole (91). Resolving the causal mechanism of sex determination will require haplotype-resolved long-read assemblies of the *Dmrt1* region, early-stage gonad-enriched transcriptomics, allele-specific expression analyses, chromatin or promoter assays, and functional validation of *Dmrt1* and linked candidate regulatory elements.

The shared 4.64 Mb sex-determining region poses a puzzle. Although its points to a ∼3.47–5.70 Mya X–Y divergence, male–male F_ST_, male–male nucleotide diversity and pairwise Y–Y divergence within the same region are nearly zero. Present-day Y haplotypes therefore carry the same ancestral Y-linked sex-determining sequence, while their surrounding Y backgrounds have accumulated private mutations independently. This contrast suggests that the sex-determining region and large NRRs have different population-genetic histories. Gene conversion, and/or rare Y–Y recombination between Y haplotypes may contribute to the nearly identical Y sequences, but they do not fully explain why a 4.64 Mb region is preserved as a sharply bounded unit across divergent Y backgrounds. This pattern points to an additional mechanism maintaining the sex-determining block, such as an inversion, a non-inversion supergene-like architecture maintained by selection or linked sexually antagonistic polymorphisms. The persistent female-biased expression of LOC120943612 and CDC37L1 near the small-NRR boundaries identifies candidate loci for testing the latter possibility. Long-read, haplotype-resolved sequencing will be essential to distinguish among these mechanisms and resolve how the shared sex-determining core is maintained.

### Extreme heterochiasmy as a rapid route to large NRRs

The large NRRs on Y_B1_^a^ and Y_B2_^a^ encompass ∼618–625 Mb, or ∼90% of the sex chromosome, and appear to be among the largest sex-linked NRR reported in vertebrates. Strikingly, the large NRR Y haplotypes coexist in the same population with the 4.64 Mb small NRR of Y_B2_^0^. The broad NRRs are consistent with the extreme heterochiasmy of anurans, in which male recombination is largely restricted to chromosome ends. Under this recombination regime, a male-determining locus located away from the terminal recombination zones can immediately behave as part of an extensive nonrecombining block, even in the absence of a series of stepwise inversions. Thus, in taxa with strong heterochiasmy, the recombination landscape itself may shape the initial physical scale of sex linkage. By resolving NRR boundaries at the whole genome scale, our resequencing data provide direct empirical support for the causal link of extreme heterochiasmy and large NRR size (Fig. 1b, Fig. 8).

This pre-existing recombination landscape complements rather than replaces classical explanations based on selection for tighter linkage between sex-determining and sexually antagonistic loci (11). Sexually antagonistic selection may still shape PAR boundaries, favour modifiers of recombination, or contribute to the maintenance of particular Y backgrounds (100). Our data show that a very large NRR can arise as a consequence of extreme heterochiasmy without requiring sexually antagonistic loci necessarily. However, this mechanism of recombination suppression would only be applicable for the sex-determining system in the heterogametic sex of which also has extreme heterochiasmy, i.e. the XY/XX species. ZW/ZZ species would need alternative mechanism of recombination restriction on the W, unless they can evolve female extreme heterochiasmy where recombination restricting to chromosomal ends occurs in ZW females. Comparisons with ZW anurans, where characterized NRRs are often smaller, are consistent with only XY systems developing large NRRs, however, there are insufficient data to determine if this is a general rule. For example, NRR sizes range from 10.7 Mb (7.7% of Chr7) in *Xenopus tropicalis* (88) to 54.1 Mb (44.2% of Chr8L) in *X. borealis* (101). Systematic sampling of XY and ZW species across anurans, combined with sex-specific linkage maps and phased sex-chromosome assemblies will be essential for testing whether extreme heterochiasmy consistently predicts NRR size.

In this respect, comparison with the Emei moustache toad (*Leptobrachium boringii*) is informative. That species also has an extensive NRR (∼394 Mb, ∼60% of sex chromosome) and documented XY sex reversal, yet phased X and Y assemblies are largely colinear, suggesting that very large NRRs need not always be produced by large inversions (102). *R. temporaria* provides an additional layer by showing that different Y haplotypes within one species can retain the same ancestral sex-determining core while differing dramatically in the extent of surrounding NRRs.

### Large NRRs without extensive Y degeneration

The large physical size of the NRR does not predict high Y degeneration in *R. temporaria*. The most differentiated Y haplotypes show the expected early signals of reduced nucleotide diversity, elevated X–Y divergence, and modest TEs and repeats enrichment, but not the later hallmarks typical of old heteromorphic sex chromosomes such as gene loss, reduced Y-linked expression, faster-X evolution or sex-biased gene expression enrichment within the NRRs. Our data suggest that the large NRRs are young or have experienced intermittent recombination, or both.

The pattern is consistent with observations from other young sex-chromosome systems in which repeat accumulation can precede major gene decay, dosage-compensation evolution, or broad-scale sexualization of gene expression (43, 69, 103–105). Young sex chromosomes in fishes and plants often show heterogeneous degeneration, weak dosage effects, and limited accumulation of sex-biased genes despite detectable X–Y or Z–W differentiation. In *R. temporaria*, the enrichment of female-biased expression in the PAR and on several autosomes, rather than specifically in the NRR, further argues against a simple feminization or masculinization model driven by sex-linked inheritance. Instead, PAR regions close to NRR boundaries remain plausible locations for sexually antagonistic polymorphisms because recombination is locally reduced (100), but this hypothesis requires fitness assays and functional tests of candidate genes.

The limited degeneration of exceptionally large NRRs provides genome-wide evidence that sex reversal can decouple NRR size from Y degeneration. Previous marker-based studies revealed multiple Y haplotypes in *R. temporaria* but lacked the resolution to distinguish among independent sex-determining origins, a linear series of Y differentiation, and shared ancestry of the sex-determining region. By comparing X–Y differentiation with Y–Y divergence across the sex chromosome, we show that the coexisting haplotypes share an old *Dmrt1*-linked sex-determining core while carrying younger, independently differentiated Y backgrounds. This genome-wide pattern reveals how two opposing processes—expansion of recombination suppression under extreme heterochiasmy and recombination-mediated homogenization through sex-reversed XY females—can act within the same population. Together, these processes generate cyclic dynamics of Y differentiation, allowing an old, shared sex-determining region, younger large NRRs, and semi-differentiated Y haplotypes to coexist.

### Sex reversal and the renewal of Y-haplotype diversity

We find heterogeneous sex-chromosome differentiation within the same population. Sex reversal is the plausible mechanism for the erosion of large NRRs and the generation of semi-differentiated Y haplotypes (62). When the resulting Y chromosomes pass again through males with chromosome end-restricted recombination, large blocks can become nonrecombining and begin differentiating anew.

The new perspective on this system brought about by genomic analysis is not the incidence of sex reversal, mobile PAR boundaries, or large NRRs; each has precedents in the sex-chromosome literature. Instead, it is the shared ancestral *Dmrt1*-linked sex-determining region, multiple younger peripheral NRRs of different sizes and differentiation levels, and a biologically plausible mechanism for both expansion and erosion of the NRR boundaries. The coexisting Y haplotypes are unlikely to represent successive stages along one inevitable path of differentiation. Instead, they appear to represent repeated, background-specific changes in the recombination landscape surrounding a conserved sex-determining region (Fig. 8b).

It is unclear how multiple Y haplotypes can coexist. Drift and sex-ratio selection would remove Y variation, especially if XX genotypes produce an approximately equal sex ratio as they do in other populations. Gene flow from neighbouring populations may reintroduce lost Y haplotypes. Temporally fluctuating selection, local adaptation, or sexual selection on male traits may also maintain Y diversity, although fitness differences among male types were not detected (79, 84). The broader geographic patterns in Switzerland and Sweden, where fully differentiated Y haplotypes, semi-differentiated Y haplotypes, and XX males co-occur at variable frequencies (78–83), suggest that the Meitreile pond is not an isolated anomaly. Instead, the coexistence of multiple male genotypes may be a recurrent and potentially long-lived feature of *R. temporaria* populations. Distinguishing neutral turnover, metapopulation extinction and recolonisation dynamics of Y haplotypes, and selection will require range-wide haplotype-resolved population genomics data integrated with environmental data, sex-ratio estimates, and direct fitness measurements across years.

## Conclusion and future directions

Our findings refine current models of young and homomorphic sex chromosome evolution by showing how recombination suppression, restored recombination, and renewed differentiation can occur within the same population. Classical models explain why recombination suppression may evolve near sex-determining loci and why nonrecombining chromosomes can degenerate over long timescales. In *R. temporaria*, however, whole-genome resolution reveals a different dynamic: an older *Dmrt1*-linked sex-determining region is shared across all Y haplotypes, whereas the broad NRR/sex-linked regions are younger and have accumulated private mutations on distinct Y backgrounds. Thus, the unit of evolutionary continuity is not an entire Y chromosome, but a conserved sex-determining region surrounded by haplotype-specific histories of recombination suppression and X–Y differentiation. Under extreme heterochiasmy, large NRRs can arise rapidly; through sex-reversed XY females, X–Y recombination can be restored outside the sex-determining region; and subsequent transmission through males can again promote Y-specific differentiation. These cyclic dynamics of Y differentiation (Fig. 8b) help explain how frog sex chromosomes can remain homomorphic and evolutionarily labile despite episodes of extensive recombination suppression.

This model makes several testable predictions. First, phased long-read assemblies should determine whether the shared 4.64 Mb region is maintained by an inversion, a supergene-like architecture, local recombination suppression without major structural rearrangement, or a combination of these. Second, controlled crosses involving sex-reversed XY females should test whether X–Y recombination is restored outside the *Dmrt1*-linked region and whether this exchange erodes Y-linked divergence. Third, early gonad-specific transcriptomic, epigenomic, and regulatory profiling should determine whether *Dmrt1* acts through allele-specific regulatory divergence, dosage effects, structural variation, or linked regulatory elements. Fourth, population-genetic models incorporating sex reversal, extreme heterochiasmy, drift, migration, and sex-ratio selection should identify the conditions under which multiple Y haplotypes are transient versus maintained over longer timescales. Finally, comparative analyses across XY and ZW anurans, and across other taxa with strong heterochiasmy, should test whether heterochiasmy generally decouples the physical extent of recombination suppression from the age and degeneration of sex-linked regions.

Overall, our results support a view of young sex chromosomes as dynamic population-genetic states rather than fixed evolutionary endpoints. In *R. temporaria*, coexisting Y haplotypes are not independent sex-determining systems and not successive steps along a single degeneration trajectory. They are divergent Y backgrounds anchored by a shared ancestral *Dmrt1*-linked region. This architecture links classical theory on recombination suppression and Y degeneration with models emphasizing recurrent X–Y recombination, shifting sex-linked boundaries, and reversible components of sex chromosome differentiation. It also provides a genome-wide framework for understanding how homomorphic sex chromosomes can persist for long periods in frogs and other lineages with labile sex determination, recurrent sex reversal, and extreme heterochiasmy.

## Materials & Methods

### Field sampling

All sampling was performed over three consecutive years (2014–2016) in Meitreile (46°22’4.9’’N, 7°9’53.1’’E), from a breeding pond located at 1,798m at lower subalpine area in the Western Swiss Alps. The exact sampling method has described previously (79, 84). Briefly, all adult samples were sampled during the short breeding season (late March to late April). Females and males were distinguished by the amplexus status and/or the presence of nuptial pad on the thumbs of the males. Toeclips or buccal cells were sampled from each individual with sterile cotton swab (59), which were taken back to the lab for DNA extractions. The sampled frogs were immediately released after swabbing in the field (84).

Approximately 10 amplexus mating pairs were caught during 2015 breeding season and brought back to the lab for egg spawning at the University of Lausanne for androgenesis experiment. Each amplexus was put in a separate tank (11 L plastic tank) in a climatic room at condition to best mimic the natural environment they were collected from (16°C, with 12:12 light: dark cycle and approximately 50% humidity). A small amount of natural pond water was brought back to the lab, together with natural vegetation from the pond, in order to minimise animal stress.

### Androgenesis experiment

Experiment protocols were approved by the ethics committee for research on animals at the University of Lausanne. The care and treatment of animals used in this research was conducted in accordance with policies on animal care provided by regulations of the Swiss government. Frogs or tadpoles were killed by immersion in 2 g/L or 0.2 g/L bicarbonate-buffered tricaine methanesulfonate (MS-222, Sigma-Aldrich) respectively.

We performed an androgenesis experiment to generate doubled haploid individuals with Y chromosomes (YY individuals) to facilitate X-Y divergence and related analyses. Viable embryos derived solely from the paternal genome were generated using protocol described previously (106). Briefly, females (wild-caught in amplexus) were injected with human chorionic gonadotropin (Sigma-Aldrich) to ovulate with the protocol described earlier (106, 107). Eggs were collected manually in individual petri dishes to inhibit premature activation. Testes were dissected from freshly killed males (with known Y chromosome haplotype) and were crushed in 1x PBS. The collected eggs were irradiated with UV (50-70,000 mJ) for 3 min and incubated with sperm for 40 min by slowly pouring the sperm suspension. These eggs then were cold shocked in an ice bath for 7.5 min and returned to 16°C for further development. We then periodically checked embryonic and tadpole development and collected individuals for DNA extraction and genotyping using sex-linked markers to detect YY individuals for further genome sequencing (Supplementary Table S2, S3).

### DNA extraction and genotyping

DNA was extracted from adult swabs or toeclips or tails of tadpoles, with an overnight treatment of 10% proteinase K (Qiagen) at 56°C. A Qiagen DNeasy kit and BioSprint 96 workstation (Qiagen) were used to obtain 100-200 µl DNA elution in buffer AE (Qiagen). For identifying the different haplotypes of Y chromosomes, sex-linked markers at the candidate sex-determining gene *Dmrt1* (*Dmrt1_1, Dmrt1_2, Dmrt1_5 and Dmrt3*) (78), and nine to 13 diagnostic sex-linked microsatellite loci (*Bfg092, Bfg131, Bfg172, Bfg053, Kank1, Bfg191, Bfg093, RtuB, Bfg266, Btemp5, Bfg021, Bfg072* and *Bfg147*) over the entire length of sex chromosome (Chr01) were used to perform genotyping analysis and protocol was described previously (78, 79, 81, 82) (Supplementary Table S1). Briefly, DNA fragments were amplified with the primers of these sex-linked markers. Primers and PCR protocol are available in these publications respectively. Afterwards, genotyping was performed with four-colour fluorescent capillary electrophoresis using an Applied Bosystems Prism 3100 sequencer (Applied Biosystems), and the generated alleles were scored using GENEMAPPER v4.0.

Haplotypes were characterized based both on the presence of Y-specific *Dmrt* alleles and the rest of sex-linked markers. Three categories of X-Y differentiation were previously detected earlier: i) XY males with Y-specific alleles fixed at all *Dmrt* region markers and 13 sex-linked markers, and different Y haplotypes were described as Y_B1_^a^, Y_B2_^a^ and so on; ii) XY^0^ (semi-differentiated Y) with Y-specific alleles fixed only at all *Dmrt* region markers but not for other five sex-linked markers; iii) XX with no Y-specific alleles fixed at any of the 13 sex-linked markers. Finally, we considered individuals with nine Y-linked markers being homozygous as (likely doubled) haploid (Y)Y individuals (Supplementary Table S2, S3).

### Genome-wide sequencing of pooled samples (pool-seq) and YY individuals

Whole-genome sequencing of both Pool-seq samples and YY individuals was performed at the University of Lausanne Genomic Sequencing Centre using an Illumina HiSeq 2500 platform with 101 bp paired-end reads. From the genotyping results, we selected male samples with three varying levels of X-Y differentiation (XY_B1_^a^, XY_B2_^a^; XY_B2_^0^; XX) and females (XX). For the fully differentiated Y haplotypes, we selected the two most abundant ones (XY_B1_^a^ and XY_B2_^a^). Given the number of collected field samples, for each group we selected several individuals to make a pool, and the sequencing coverage was roughly equal to the number of individuals per pool (detailed sample number and read coverage see Supplementary Table S4). After sample selection, we performed DNA extraction for each selected sample separately and checked their precise DNA concentration using Qubit. We then equalized DNA concentration across samples by either diluting or evaporating excessive DNA solution, to avoid possible bias in the DNA representation within each pool. The 2*µl* DNA samples with similar concentration were taken individually to each pool group, and total pool DNA was carefully mixed thoroughly by pipetting up and down 20 times. For each pool, we used two technical replicates for DNA library preparation and genome sequencing.

Doubled-haploid YY individuals were selected based on the homozygosity of nine different sex-linked markers along the sex-linked linkage group (Supplementary Table S2). We first conducted genotyping of fathers from all collected amplexus pairs and only selected those having Y haplotypes for the androgenesis experiment. Since only the XY fathers were selected, all sex-linked markers were expected to be heterozygous, thus all homozygous individuals with Y haplotypes were considered as doubled haploid YY individuals (Supplementary Table S3). The successful rate of androgenesis was rather low and many tadpoles were selected for genotyping before the final six pools of YY individuals were detected. Since all genotypes of their fathers were obtained first, we only conducted genotyping of four markers located at the *Dmrt* region. Both YY individuals with fully differentiated Y chromosome (Y_B1_^a^, Y_B1_^a^) and proto-Y were selected for sequencing (Y_B1_^0^, Y_B1_^0^, Y_B2_^0^, Y_B2_^0^) with standard paired-end short reads. For the (Y_B1_^a^) YY individual, we further conducted mate-pair sequencing with insertion sizes 3kb, 5kb, and 15kb to facilitate the (essentially) haploid genome assembly.

### Mapping and variant calling

For each pool of the five pool-seq datasets, before and after read quality trimming, we assessed paired-end reads quality using FASTQC v0.11.9 (108) with default parameters and retained reads passing all the read quality criteria. We trimmed the raw reads to remove sequencing adaptors and low-quality base pairs using Trimmomatic 0.39 (109) with the following parameters: “ILLUMINACLIP: 2:30:10 SLIDINGWINDOW:4:20 MINLEN:36. The trimmed reads were mapped to the published chromosome-level genome assembly of one XX female individual using bowtie2 v2.4.5 (110) with the following parameters: “--very-sensitive”. We filtered out the unplaced scaffolds (only 1.74% of the genome) from the mapping files. For each pool, we split mapping files per chromosome using SAMtools v1.16.1 (111) allowing parallel computation in downstream analysis. We removed duplicates using the following functions of SAMtools v1.16.1 (111):collate, fixmate with -m option, sort, markdup with -r option. Unmapped reads were filtered out and only properly paired mapped reads with a mapping quality above 20 were retained using SAMtools v1.16.1 using the following parameters with the SAMtools view function: quality = 20; F = “0×4”; f =“0×2”. We used BCFtools v1.15.1 107to call variant using the following functions: *mpileup* with -a “FORMAT/AD, FORMAT/DP” option and *call*. We filtered out SNPs with a minimum basecall quality below 20 and minimum depth of coverage of 5 for each pool, using the *bcftools filter* function.

### Putative centromere imputation

To account for variation in sequencing depth, SNP counts were normalized within each pool using the autosomal median. The median SNP value per pool was computed from all autosomes. For all genomic windows, normalized SNP counts were calculated as log₂ (SNP count + 1) and smoothed using a rolling mean with a window size of 50. For each chromosome, breakpoint detection was performed on the smoothed log₂-transformed SNP density. A linear regression model of SNP density against genomic position was fitted, followed by segmented regression using the segmented R package (112). The model included up to seven breakpoints per chromosome. For each fitted segmentation, estimated breakpoint positions, segment slopes, and the corresponding Davies tests for changes in slope were extracted. All breakpoints were plotted against the smoothed SNP profiles to allow visual inspection. For each chromosome, a subset of breakpoints was manually selected based on clear inflection points corresponding to pronounced shifts in SNP density. These selected breakpoints were interpreted as boundaries of putative centromeric regions, consistent with expected reductions in polymorphism in pericentromeric domains. When only one breakpoint was selected, the second boundary was set to the end of the chromosome to ensure a contiguous interval. For each chromosome, the smallest and largest selected breakpoint positions were taken as the start and end of the putative centromere.

### Read coverage ratio and SNP count ratio between male and female pools

Overlapping windows of 1,000,000 bp window length and 500,000 bp interval stride across each of 13 chromosomes from the reference genome assembly were generated using bedtools v2.30.0 (113) and its *makewindows* function. For each pool, we then used the filtered mapping files to compute coverage across overlapping window using SAMtools v1.16.1 (114) and its *bedcov* function. Similarly, we used variant calling output files to compute SNP counts across overlapping window using the *coverage* function of bedtools v2.30.0 with the “-counts” command line option (113). We then calculated a normalized coverage value per window by dividing its raw value by the autosomal median value. For computing the autosomal coverage, the putative centromere region was removed. For each window, we computed the log₂ ratio of normalized SNP density and depth of coverage for each male pool relative to the female pool. SNP counts were incremented by one, normalized within each pool using the autosomal median derived from autosomes with centromeric regions excluded, and transformed to log₂ ratios between male and female values. For each chromosome, rolling means of the log₂ SNP ratios were calculated using a fixed-size sliding window of 50. Confidence intervals were obtained exclusively from autosomal windows outside centromeric regions. For each male pool, the 2.5th and 97.5th percentiles of the autosomal rolling-mean log₂ ratios were used as empirical 95% confidence limits, following the percentile-based approach recommended by (120). These confidence intervals were subsequently used to annotate the sex chromosome and autosomes in the final plots. The calculations and plots were performed in R v4.2.1 (115) using custom R scripts. All related scripts and R codes is available in the GitHub upon manuscript acceptance.

### Sliding-window FST scanning analysis

For each pool and for each pairwise comparison between male pools and female pool, we generated pileup files using SAMtools v1.16.1 *mpileup* function from the filtered mapping files. Pileup files were converted into sync files using grenedalf (114) sync function for which we only retained position matching the following criteria: a minimum base quality of 20 (--pileup-min-base-qual), minimum base count of 2 for a nucleotide to be considered as an allele (--filter-sample-min-count), minimum read depth (--filter-sample-min-read-depth) and maximum read depth (--filter-sample-max-read-depth) per sample of 30 and 120 respectively for the pools with 120 individuals (Y_B1_^a^, XX females, XX males). F_ST_ was computed using grenedalf (116) applied to the pileup-derived sync files. For each focal–reference pool pair and each chromosome, the grenedalf *fst* module was executed with the following settings: method = *unbiased-nei*, window type = *interval*, window-average policy = *window-length*, window interval width = 10,000 bp, and window interval stride = 5,000 bp. Pool sizes were provided per chromosome, and computations were performed using the reference genome and pool-size tables as specified in the workflow. The resulting output consisted of per-window F_ST_ estimates for each chromosome. In parallel, the same sync files were supplied to the grenedalf *frequency* function to extract reference and alternate allele counts and allele frequencies genome-wide, using the quality and coverage filters defined in the configuration (minimum base quality = 30, minimum count = 2, minimum coverage = 30, maximum coverage = 120). Similar approach and filtering process and criteria were conducted for pairwise comparison among the four male pools.

### Nucleotide diversity π and pairwise Dxy divergence analysis

Nucleotide diversity (π) was estimated from pool-seq alignments using grenedalf diversity (116). Five pooled BAM files were analysed. Reads were mapped to the *R. temporaria* reference genome from NCBI (accession: GCF_905171775.1). For each pool, sites were filtered using a minimum mapping quality of 20, minimum base quality of 20, minimum sample allele count of 2, minimum read depth of 30, and maximum read depth of 180. Nucleotide diversity was calculated in sliding genomic windows of 100 kb with a 50 kb step, using parameters valid-loci for window-averaging and the unbiased Nei estimator. For visualization, window (100 Kb) genomic midpoints were used. For chromosome-wide profiles, π values were smoothed using a rolling mean over 50 windows (5 Mb), with a step of 20 windows (1 Mb). For zoomed views, π values were smoothed using a rolling mean over 5 windows (500 kb), with a step of 1 window (100 Kb). Statistical comparisons were performed using paired window-wise tests with the XX female pool as the reference. To reduce non-independence among overlapping windows, a non-overlapping subset of windows was used. For each comparison, paired t-tests were performed on finite, positive π estimates, and mean differences, median differences, 95% confidence intervals, paired Cohen’s (d_z), and paired Cohen’s (d) were calculated. Multiple comparisons were corrected by Benjamini–Hochberg false discovery rate.

Pairwise genetic divergence was estimated from whole-genome variant calls using pixy (117). To include an outgroup/reference individual, publicly available PacBio HiFi reads were downloaded (accession: ERR7012640-42), concatenated mapped to the reference with minimap2 using the map-hifi preset and WSI as read group. Alignments were filtered to retain primary alignments with a minimum mapping quality of 20 and sorted with samtools. All-sites genotype calls were generated for the HiFi sample using bcftools mpileup and bcftools call. The pileup and calling were performed separately by chromosome, with the following parameters: -q 20 -Q 20 -X pacbio-ccs-1.20 for mpileup and --ploidy 2 -c -A -a GQ for the call, retaining invariant sites and alternate alleles. The per-chromosome VCFs were concatenated. Haploid Y-linked samples were called separately using bcftools mpileup and bcftools call using these parameters: -q 20 -Q 20 --max-depth 1000 for mpileup and -m --ploidy 1 -A -f GQ for call. Sites were filtered by masking genotypes with low support, using a threshold of QUAL < 10 or total depth < 10 for variant sites using bcftools filter. The diploid HiFi all-sites VCF and the haploid Y-sample all-sites VCF were then merged with bcftools merge to produce a combined all-sites VCF for divergence estimation using pixy. Pairwise sequence divergence was calculated on chromosome 1 using windows of 1 Mb windows for chromosome-wide visualization and 500 kb for the focal region.

### dN/dS and πΝ/πS analysis

To estimate synonymous (d_S_) and non-synonymous (d_N_) substitution rates between X- and Y-linked coding sequences, we used paired-end sequencing reads from doubled haploid YY individuals representing the Y_B1_^0^, Y_B2_^0^, and Y_B1_^a^ haplotypes. Reads were processed following the same quality-control, trimming, and mapping procedures described above. Variant calling was performed with bcftools call using the haploid mode (“--ploidy 1”). For each sample and chromosome, SNP-only BCF files were combined with the reference genome to generate consensus sequences using bcftools consensus. Coding sequences (CDS) were then extracted from each consensus genome with gffread based on the reference GFF annotation. Reference CDS were extracted in parallel using the same procedure. For every chromosome and sample, CDS from the reference and the corresponding sample were split into per-gene FASTA files using a custom Python script (split_cds_per_gene.py). These gene-specific alignments were used as input for pairwise dN/dS estimation.

For each gene, codon-based alignments in PHYLIP format were analysed with the yn00 program from PAML v4.9j (118), executed through Biopython’s PAML interface (119) via the script run_paml_pairwise.py. The script extracted the dN, dS, and dN/dS values for each gene, and returned a summary table for every sample and chromosome. Only coding sequences with valid codon length (multiples of three) and without premature stop codons were retained. For each YY haplotype, dN, dS, and dN/dS values were summarized per gene, and results were restricted to the longest isoform. The final dataset comprised one dN, one dS, and one dN/dS estimate per gene and per Y haplotype.

Synonymous and non-synonymous nucleotide diversity (πS and πN) and their ratio (πN/πS) were estimated for each pool using SNPGenie v3.0 (120). Variant calls were first filtered and split by chromosome. For each chromosome and pool, we generated strand-specific input files consisting of (i) VCF files extracted from per-chromosome SNP BCFs using bcftools view, (ii) FASTA sequences extracted from the reference genome, and (iii) GTF annotations restricted to CDS features whose lengths were multiples of three. For the negative strand, corresponding FASTA, VCF, and GTF files were reverse complemented using custom Perl utilities to ensure correct codon orientation (see SNPGenie manual). SNPGenie was then executed separately on the positive and negative strand datasets, using its recommended settings (VCF format 3 and codon-based CDS parsing). For each pool and chromosome, SNPGenie computed per-gene and genome-wide estimates of π_N_, π_S_, and π_N_ / π_S_. The resulting outputs from both strands were retained for downstream analyses.

### Molecular clock and age of sex chromosomes

As mentioned in the above section, we used paired-end sequencing reads from double-haploid YY individuals representing the Y_B1_^0^, Y_B2_^0^, and Y_B1_^a^ haplotypes to estimate the synonymous (d_S_) substitution rate across the genome. The molecular clock derived from the Tibetan frog *Nanorana parkeri*, to estimate the age of *R. temporaria* sex chromosomes (92). Since the molecular clock was provided as 0.776 × 10^−9^ substitutions per site per year (±1.34 × 10^−12^ SE), we calculated the age as follows: (d_S_)/rate = d_S_/(0.776 × 10^−9^). To encounter possible outlier d_S_ values to bias the age estimate, d_S_ values of all coding genes at the small NRR or large NRR were used after correcting for within-XX male pool polymorphism.

### Defining genomic compartments from FST variation

We used the F_ST_ estimates produced by grenedalf for all focal–reference pool comparisons to infer large-scale genomic compartments along the sex chromosome. Per-chromosome F_ST_ results in CSV format were imported and parsed to extract genomic coordinates, window boundaries, and F_ST_ values. For each pool comparison, the midpoint of each window was computed, and windows with missing or undefined F_ST_ values were excluded. To reduce local noise, we applied a rolling mean with a window size of 1,000 windows. For each male and female pool, change-point detection was applied to the smoothed F_ST_ profiles using the PELT and Binary Segmentation (BinSeg) algorithms implemented in the changepoint R package 116. The number of expected change points (Q) was specified per pool based on graphical inspection of the underlying profiles. For each pool, the genomic positions of inferred change points were converted from window midpoints to the nearest window start coordinate in the original data. These change points defined the boundaries of successive genomic compartments along the sex chromosome. For each pool, compartment start and end coordinates were defined by combining the first genomic position on the chromosome, the inferred change-point positions, and the annotated end coordinate of the sex chromosome. Each interval was recorded as a distinct genomic compartment specific to that pool. Autosomal compartments were defined separately. For all pools, autosomal regions were taken directly from the annotated genomic intervals in the reference GFF file. Each autosome was assigned a unique compartment label per pool without further segmentation. All sex-chromosome and autosomal compartments were then combined into a unified BED file, which was exported for downstream analyses.

### TE insertion and abundance distribution analysis

A *de-novo* TE library was generated using RepeatModeler2 on the *Rana temporaria* reference genome aRanTem1.1 through the Dfam TE tools 1.88.5 container (https://github.com/Dfam-consortium/TETools) (121). Later, Cd-hit was used to reduce redundancy on the TE library with the following settings: -d 0 -aS 0.8 -c 0.8 -G 0 -g 1 -b 500 118. PoPoolationTE2 was used to detect TE insertion abundance on Poolseq data of one female and four male pools with varying Y chromosome differentiation (113). Briefly, the newly generated TE library and aRanTem1.1 were used as reference TE and reference genome respectively. PoPoolationTE2 requires a TE masked genome that was created through RepeatMasker masking TE sequences of the *de-novo* generated TE library (122, 123). Later, the masked genome and TE library were merged. Reads were reordered using BBMap repair and later mapped against the merged genome using bwa and processed using PoPoolationTE2 in separate mode with default parameters (124).

### RNA-seq data generation, processing and major allele ratio analysis

Adult samples from the same Swiss Alpine population were previously collected in the breeding season of 2013 for transcriptome sequencing. In total, 13 adult males and 3 adult females were captured and sacrificed following standard euthanasia methods previously described (76, 90). Brain and gonad tissues were removed and flash frozen in liquid nitrogen, followed by RNA extraction using the Trizol Method. To determine the haplotype of the male Y chromosome, males were genotyped with the 12-16 sex-linked markers we described in the above section on genotyping. Three males were identified as undifferentiated, five as semi-differentiated, and five as fully differentiated Further investigation into their specific haplotype was not undertaken. Paired-end sequencing producing 100 bp reads was performed by the University of Lausanne Genomic Technologies Facility.

Quality was assessed using FASTQC v0.11.9 (www.bioinformatics.babraham.ac.uk/projects/fastqc). Adapters were removed and reads were trimmed in Trimmomatic with the parameters: ILLUMINACLIP:TruSeq2-PE.fa:2:30:10:2:keepBothReads LEADING:20 TRAILING:20 SLIDINGWINDOW:4:20 MINLEN:50 123. Trimmed reads were then aligned against the *Rana temporaria* reference genome using the 2-pass method in STAR (125), in which the first pass collects the splice junction information to be included in the genome index, and the second pass uses this information to refine the alignment. As the chromosomes are *R. temporaria* are particularly large and exceed the capability of GATK to index using the typical .bai format, all indexing steps were performed using the .csi format. We then processed the according to GATK best practices for RNA-seq data (126), including adding read groups, marking duplicates, and splitting reads. We then used Haplotype Caller to call variants and combined (using GATK CombineGVCFs) and genotyped (using GATK GenotypeGVCFs the resulting vcf files. These were then stringently filtered (QD < 2.0, FS > 30.0, SOR > 3.0, MQ < 40.0, MQRankSum < -12.0, ReadPosRankSum < -8.0, DP < 10) to produce an initial, high-confidence variant file. This variant file was then using in a second iteration of aligning reads using STAR in WASP mode (127), to further improve alignments. Again, a two-pass alignment method of STAR was performed with trimmed sequences, followed by the same pipeline of GATK processing (adding read groups, marking duplicates, and splitting reads). Again, variants were called and resulting vcfs were combined and genotyped as above. During the GATK VariantFiltration step, we included the previous high-quality variant file produced in our first round of STAR alignments, and filtered our raw vcf from our second alignment with slightly less stringent parameters (QD < 2.0 || FS > 60.0 || MQ < 30.0 || SOR > 3.0 || ReadPosRankSum < -10.0 || MQRankSum < -10.0 || DP < 8). We extracted biallelic SNPs that passed all previous filters using bcftools (138). We implemented GATK ASEReadCounter, which produces a .tsv file for each file with the number of reads for the reference and alternative alleles (128).

We next aimed to detect evidence of allele-specific expression. First, we removed SNPs from our filtered variant site if there were 5 in a 100bp window, as this can indicate bias towards the mapping allele during the mapping step (129). If the two alleles have equal expression, we expect a major allele ratio of 0.5. Deviations from 0.5 indicate a bias towards one allele or the other. As our filtering criteria above likely resulted in the inclusion of mapping or genotyping errors in our dataset, we then selected for polymorphisms with a minimum depth of 30, with at least 5 reads representing the reference and the alternative genotype each. Then or each SNP, we conducted a two-tailed binomial test and corrected or multiple using the Benjamin-Hochberg method. To assess significance for allelic specific expression, we permutated Major Allele Ratio (MAR) values within each genotypic class 10,000 and compared the difference between the median MAR for sex chromosomes and autosomes. P-values were computed as the proportion of replicates which exceeded the median difference in MAR between sex chromosomes and autosomes. Further, to detect evidence of allele-specific expression in differentiated regions of chromosome 1, we computed the major allele ratio in sliding windows of 100 SNPs using the rollmean function in the R package zoo (130).

### Differential gene expression and enrichment on sex chromosome

Meitreile RNA-seq datasets of *R. temporaria* including gonadal (testis and ovary), and brain tissues sampled from males with three Y-differentiation levels and females were used in differential expression analysis. Raw sequencing reads were assessed using FastQC v0.11.9 both before and after trimming to evaluate read quality metrics (www.bioinformatics.babraham.ac.uk/projects/fastqc). Adapter removal and quality trimming were performed with Trimmomatic v0.39 in paired-end mode, using the following parameters: ILLUMINACLIP (adapter clipping), HEADCROP:12 (removal of sequencing bias at the start), SLIDINGWINDOW:4:15 (quality filtering), and MINLEN:36 (minimum read length). Both paired and unpaired reads were retained for downstream analysis.

Transcript-level quantification was performed using Salmon *v1.10.3* (131). To prepare a decoy-aware transcriptome index, the *R. temporaria* reference transcriptome and genome (RefSeq assembly GCF_905171775.1, aRanTem1.1) were concatenated to create a “gentrome” file. Decoy sequences were extracted from the genome FASTA headers, and the Salmon index was built using the --decoys and --validateMappings options to ensure accurate alignment. Salmon quantification was run in quasi-mapping mode using paired-end trimmed reads. For each sample, the following parameters were used: -l A for automatic library type detection, --validateMappings for selective alignment, and --gcBias correction. Transcript-level abundances were quantified as TPM (Transcripts Per Million), which normalizes for both transcript length and sequencing depth. These normalized values were used for gene-level expression comparisons, while raw read counts were used for differential expression analyses.

Gene annotation was manually curated from the *R. temporaria* reference genome assembly (GCF_905171775.1, aRanTem1.1). Custom annotation tables were constructed to support downstream gene-level analyses. Two annotation sets were generated: i) Annotation_CDS: based on coding sequence, incorporating gene name, product, chromosome, coordinates, length, and effective length, merged with Salmon quantification results using the CDS reference. ii) Annotation_Transcriptome: based on mRNA and non-coding transcript identifiers, excluding rRNA, snRNA, snoRNA, and tRNA, and similarly merged with Salmon outputs to extract accurate transcript length and effective length estimates. Salmon RNA-seq quantifications were used to refine effective lengths, accounting for transcript-specific sequence characteristics. Duplicate FASTA entries were filtered out to ensure consistency between quantification and annotation. Chromosome names were cleaned and standardized. The resulting annotation tables were integrated with expression data enabling positional and functional analyses of differentially expressed genes.

Differential expression analysis was performed using the EdgeR *v4.2.2* package in R (132, 133). Genes with low expression were removed through a two-step filtering process: first, genes with average log₂-counts per million (CPM) ≤ 0 were excluded; second, genes were retained only if at least two samples exhibited CPM ≥ 1. Read counts were normalized using the trimmed mean of M-values (TMM) method to correct for differences in library size and composition. Gene-wise dispersion estimates were obtained using quasi-likelihood methods (*glmQLFit*), and differential expression was tested using quasi-likelihood F-tests (*glmQLFTest*). Experimental contrasts were specified via a custom contrast matrix, and p-values were adjusted for multiple testing using the Benjamini-Hochberg false discovery rate (FDR) procedure. Exploratory analyses were conducted prior to model fitting. Multidimensional scaling (MDS) plots based on log_2_ fold change (log₂FC) and biological coefficient of variation (BCV) were used to assess sample clustering and detect potential outliers. Mean-variance plots of log2 CPM values were used to evaluate expression dispersion. Sex-biased gene expression was defined using a combined threshold of FDR < 0.05 and |log₂FC| ≥ 1, corresponding to at least a two-fold expression difference.

Differential expression results obtained from EdgeR *v4.2.2* were used to investigate whether sex-biased gene expression was non-randomly distributed across chromosomes (134). For each chromosome and each level (undifferentiated, semi-differentiated, and fully differentiated), whether the distribution of log₂FC values differed from the rest of the genome was tested using permutation tests. Comparisons involving sex chromosome (Chr1) were performed against all autosomes (Chr2-13), while autosomes were compared to all other autosomes excluding the focal one. Resulting p-values were adjusted using the Benjamini-Hochberg procedure to control the false discovery rate.

To examine the distribution of sex-biased genes across chromosomes, gene count summaries were generated stratified by chromosome, differential expression status, and direction of expression bias. Using DE gene lists identified by *EdgeR* (FDR < 0.05) across each contrast, we defined sex-biased genes as those with |log₂FC| > 1 and classified them as male-biased or female-biased. Genes not meeting the DE criteria were labelled as non-differentially expressed (non-DE). For each chromosome and contrast, we recorded the total number of tested genes, the number of DE genes, and the number of sex-biased genes. To determine whether sex chromosome was significantly enriched for sex-biased genes, we performed permutation tests. Similarly, we also performed permutation tests to assess compartment enrichment on Chromosome 1 for PAR and NRR regions compared to each other and autosomes. P-values were adjusted for multiple testing using the Benjamini-Hochberg method.

### Gene expression analysis in the sex-determining region

The expression profiles of 29 putative candidate genes located within the semi-differentiated region of the proto-Y chromosome were examined to investigate their potential involvement in sex determination. Differential expression results from the EdgeR pipeline were used to extract log₂FC values and associated false discovery rates for each gene across the three Y differentiation levels. Genes were classified as sex-biased if they showed a significant expression difference (FDR < 0.05) in either direction (log₂FC ≠ 0); genes without significant differential expression were labelled as “Not significant,” and those with missing log₂FC values due to expression below detection thresholds were labelled as “Not expressed.” To visualize these patterns, absolute log₂FC values were plotted for each gene, ordered by genomic position, with separate panels for each Y haplotype. Additionally, six genes exhibiting strong sex bias (|log₂FC| > 4) were selected for detailed visualization of expression levels. Transcripts per million (TPM) values were plotted across gonadal (ovary and testis) and somatic (brain) tissues. This approach enabled the assessment of tissue specificity in expression patterns, supporting the identification of candidate master sex-determining genes.

### Dendrogram clusters along the sex chromosome

For all the RNA-seq datasets, filtered VCF files from the Swiss Alpine population Metreille were further processed using VCFtools (v0.1.16) to generate region-specific VCFs corresponding to previously defined pseudoautosomal and nonrecombining regions of chromosome 1 in 10 Mb and 50 Mb windows, respectively, and specifically the region including *Dmrt1*, *Dmrt3* and *Dmrt2*. Trimmed VCF files, genome assembly, and annotation were imported into RStudio (v.4.5.0) using the vcfR package (135). Genotypes were extracted and recoded into numeric format to represent genotype classes (homozygotes coded as 0 and heterozygotes as 1). Additional filtering removed monomorphic sites and sites with missing data. As RNA seq data can only be genotyped for samples with gene expression, we examined two different datasets: one with all individuals and one limited to males (as females did not express some genes of interest, including *Dmrt1*). Dendrograms were generated by hierarchical clustering of samples using Euclidean distance calculated from genotypic data using the pheatmap package (https://github.com/raivokolde/pheatmap). To examine SNP patterns of homozygosity and heterozygosity, the dplyr package was used to extract sites where females and/or XX males show homozygosity and all other samples showed heterozygosity. These sites were then referred to the annotated Rana temporaria genome to record corresponding gene names and locations.

## Author contributions

W-JM, PV, NP designed the study. PV, W-JM collected frog samples, conducted genotyping analysis and performed all androgenesis and related experiments. FC, PV, W-JM, AH, EU, KLD, RF and MAT performed bioinformatics analyses and conducted initial figure visualization. AB designed and generated RNA-seq dataset. NP and W-JM obtained the fundings for the project. W-JM and PV supervised the work. AH, W-JM, and PV largely revised the figure visualization. W-JM drafted the manuscript with the help of FC, with significant contribution from PV, which was improved and commented by all co-authors. All authors agreed on the final version of the manuscript.

## Capture and ethical permits

Capture permits were delivered by the division Biodiversité et Paysage (DGE Vaud), and ethical permit delivered by the Veterinary office of the Canton Vaud (authorization 2287).

## Acknowledgments

We are grateful for Mark Kirkpatrick for discussions and advice throughout the development of this project, for Chengde Chang for insightful suggestions on the pool-seq experimental design, for Glib Mazepa for continuous help with androgenesis experimental design. We thank Karim Ghali, Guillaume Lavanchy, Glib Mazepa, Yvan Vuille, Charlotte Karsegaard, Nathalie Jollien, Maud Baudraz and Kim Schalcher for their help during the Meitreile fieldwork. We are grateful for Roberto Sermier for his significant help in the wet lab. We thank Nicolas Rodrigues for sharing some additional frog samples from this population. We thank James Hagan for consultation with various statistical aspects including helping a more reasonable way of computing the 95% confidence interval. We appreciate helpful and constructive feedback and discussion with Julien Joseph. We are thankful for two anonymous reviewers for constructive comments to clarify and improve an earlier version. The computations were performed at the Vital-IT (http://www.vital-it.ch) Centre for high-performance computing of the SIB Swiss Institute of Bioinformatics, the VSC (Flemish Supercomputer Centre), funded by the Research Foundation - Flanders (FWO) and the Flemish Government, and Louisiana Optical Network Infrastructure (LONI: http://www.loni.org). This work was supported by the Swiss National Science Foundation (Sinergia grant CRSII3_147625 to NP), and by the European Union (ERC starting grant, FrogWY, 101039501), a starting grant from research council of Vrije Universiteit Brussel (OZR4049), and an FWO junior research grant (G0A3B26N) to W-J M.

## Data and code availability

Raw WGS sequencing reads have been deposited in the NCBI Sequence Read Archive under BioProject PRJNA1399088 (BioSample accessions SAMN54473431 – SAMN54473446), raw RNA-seq reads under BioProject PRJNA1400602 (SRR36758587 - SRR36758609). All code used for the analyses is available at https://github.com/TheWMaLab/poolseq and has been archived on Zenodo (doi: 10.5281/zenodo.17599170). All datasets and code will be made publicly available upon acceptance of the manuscript.

## Supplementary files

Supplementary File 1 contains Figures S1–S28. Supplementary File 2 contains Tables S1–S22. All supplementary materials are provided as separate figure and table files, compiled into two documents.

