## Supplementary figures for "Coexisting Y haplotypes reveal cyclic sex chromosome differentiation around a shared sex-determining region"

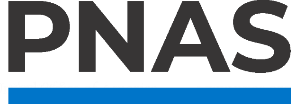

### Supporting Information for:

Coexisting Y haplotypes reveal cyclic sex chromosome differentiation around a shared sex-determining region

Fantin Carpentier^1^, Antoine Houtain^1^, Ezgi Unal^1^, Kelsey L. Doucette^2^, Ricard Fontserè^1^, Alan Brelsford^3^, Melissa A. Toups^2,4^, Nicolas Perrin^4^, Paris Veltsos^1,4†^, Wen-Juan Ma^1,4†^

1. Research group of Ecology, Evolution and Genetics, Biology Department, Vrije Universiteit Brussel, Brussels, Belgium

2. Department of Biology, University of Louisiana at Lafayette, Lafayette, LA, USA

3. Evolution Ecology and Organismal Biology Department, University of California, Riverside, CA, USA

4. Department of Ecology and Evolution, University of Lausanne, Lausanne, Switzerland

^*^Equal contribution:

The two authors contribute to the work equally.

^†^Correspondence:

This PDF file includes:

Supporting text

Figures S1 to S28

Tables S1 to S22

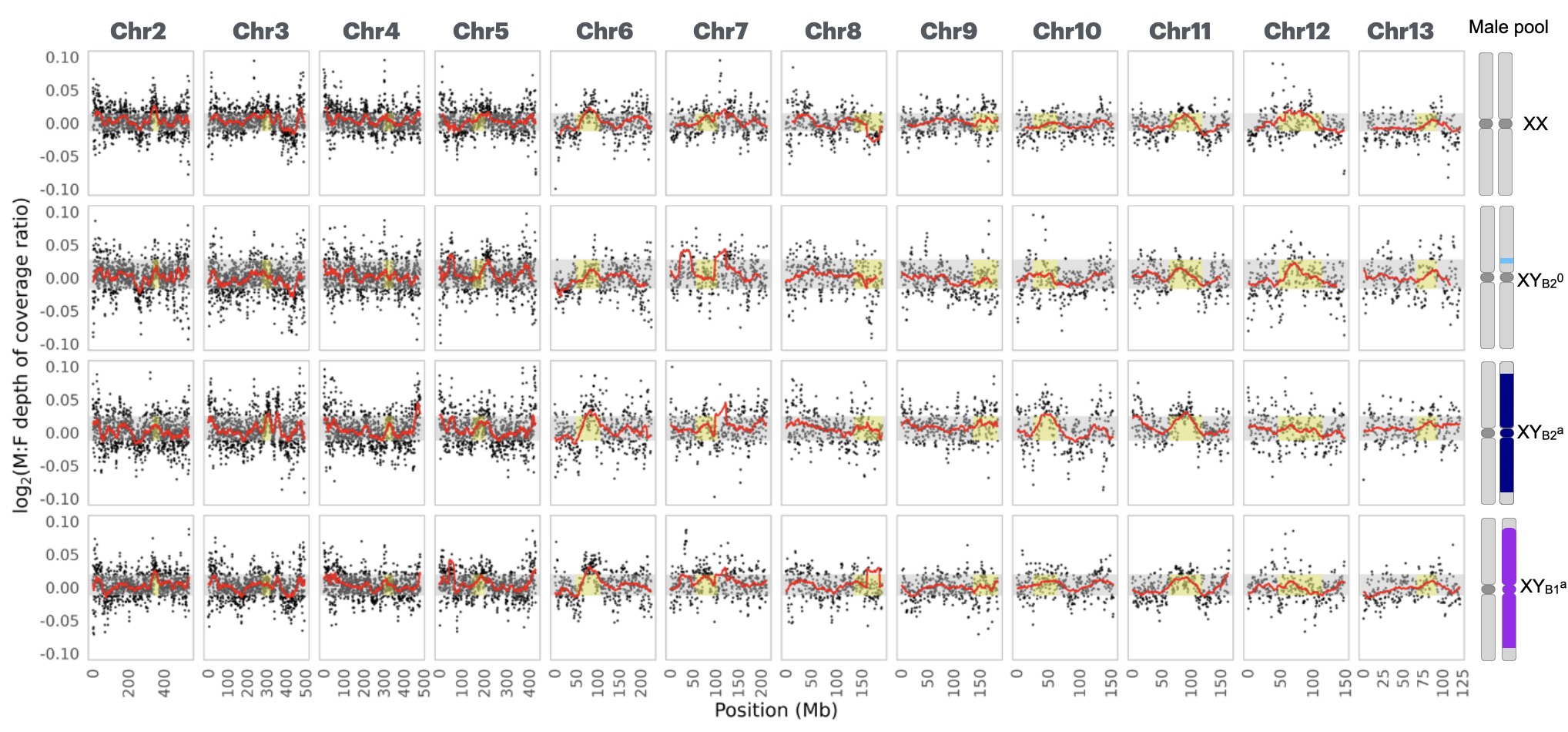

**Figure S1**: Log_2_ male-to-female read coverage ratio between each male pool and the female pool. Each male pool (XX, Y_B2_^0^, Y_B1_^a^, and Y_B2_^a^) was compared to the female pool along each of the 12 autosomes (chr2 to chr13). The dots represent average read coverage ratio between each male and female pools within 1 Mb overlapping window (stride = 500 kb). The red line indicates the rolling mean (window size = 50 Mb). The grey rectangle marks the central 95% of the rolling mean of autosomal values. Yellow zones indicate tentative centromeric regions.

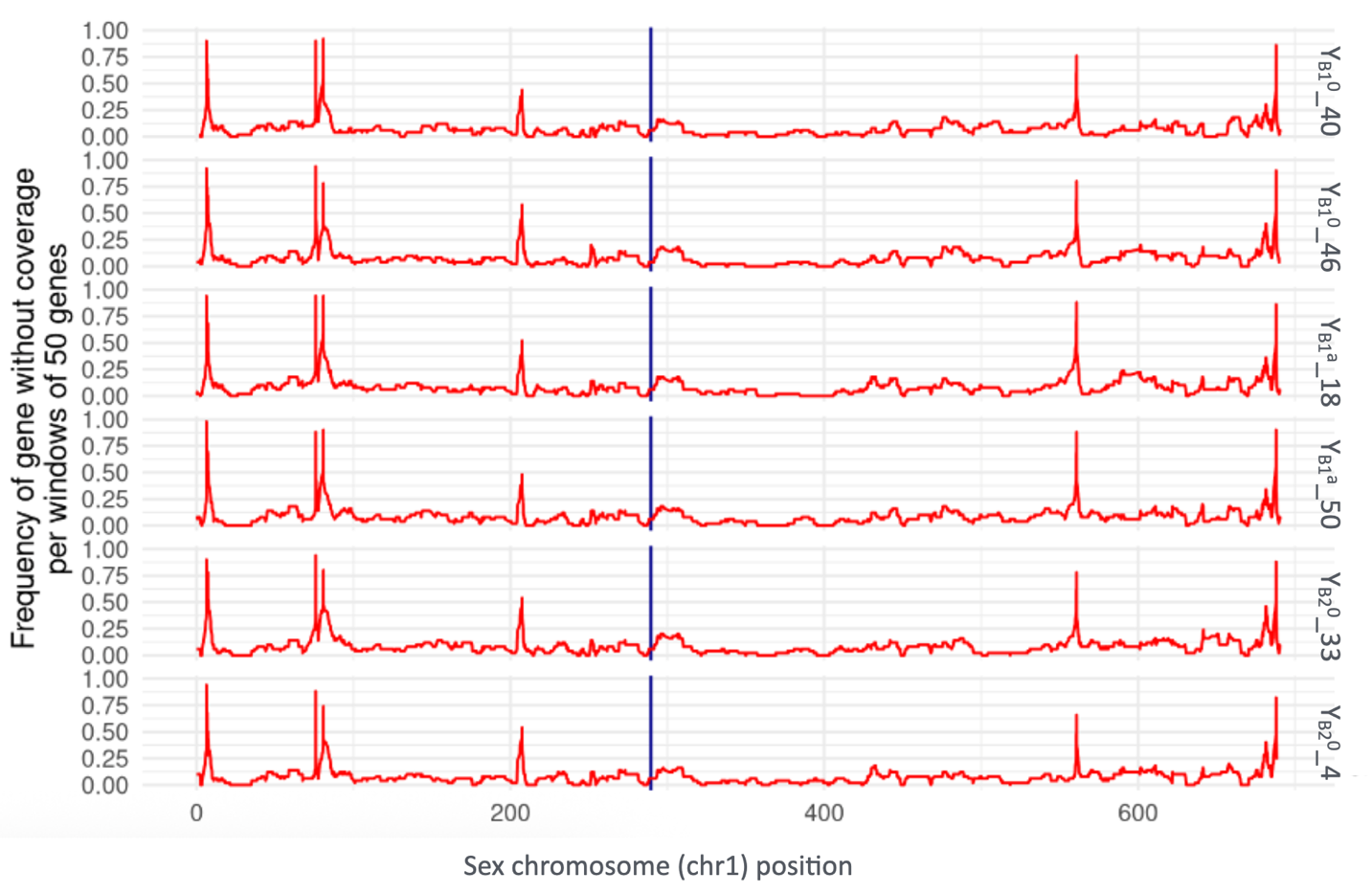

**Figure S2**: Frequency plot of genes in a non-overlapping window of 50 genes with zero coverage support when mapping to the XX reference genome, along the sex chromosome for the 6 double haploid YY individuals. The blue vertical line is the region for sex-linked marker at genes of *Dmrt1* and *Dmrt3*.

(a)

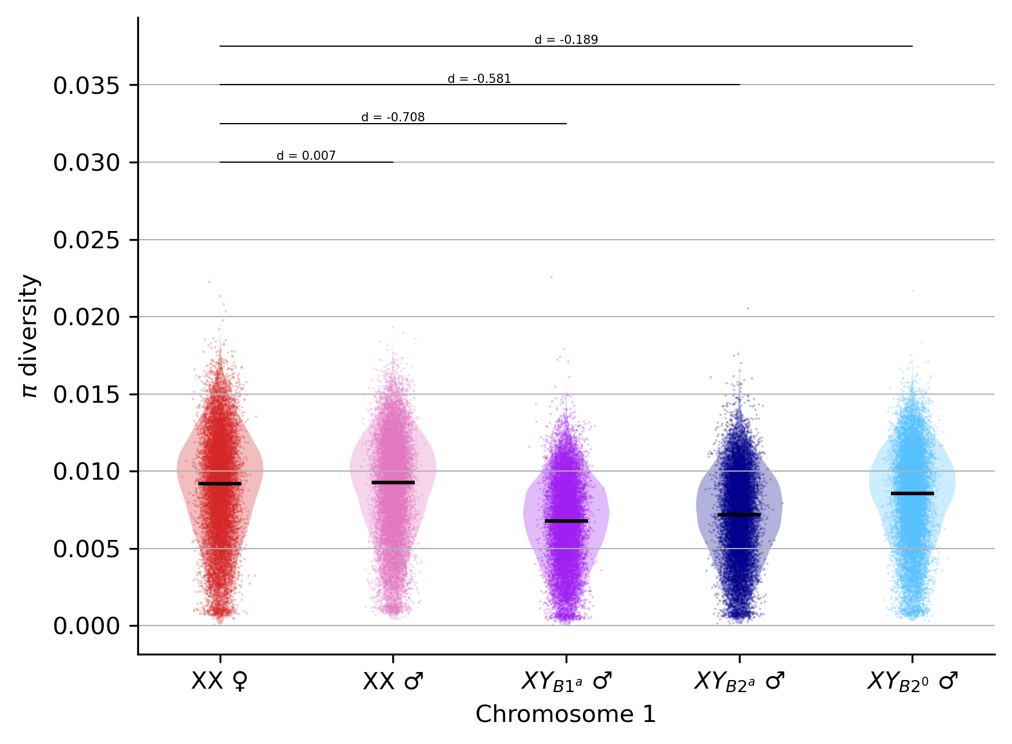

(b)

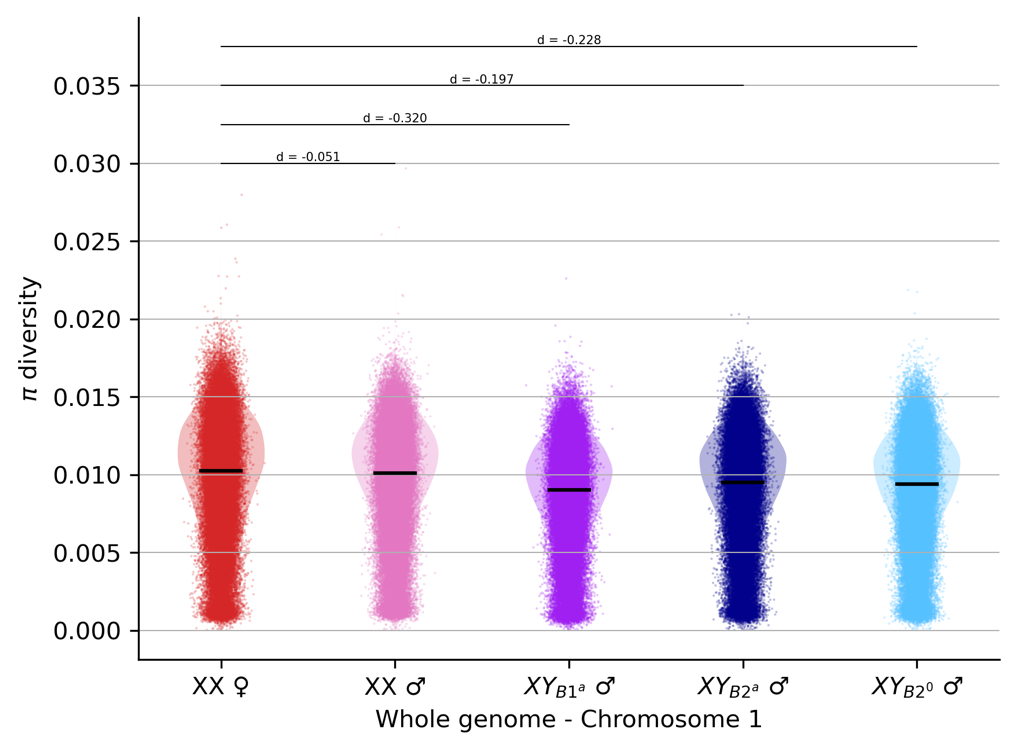

(c)

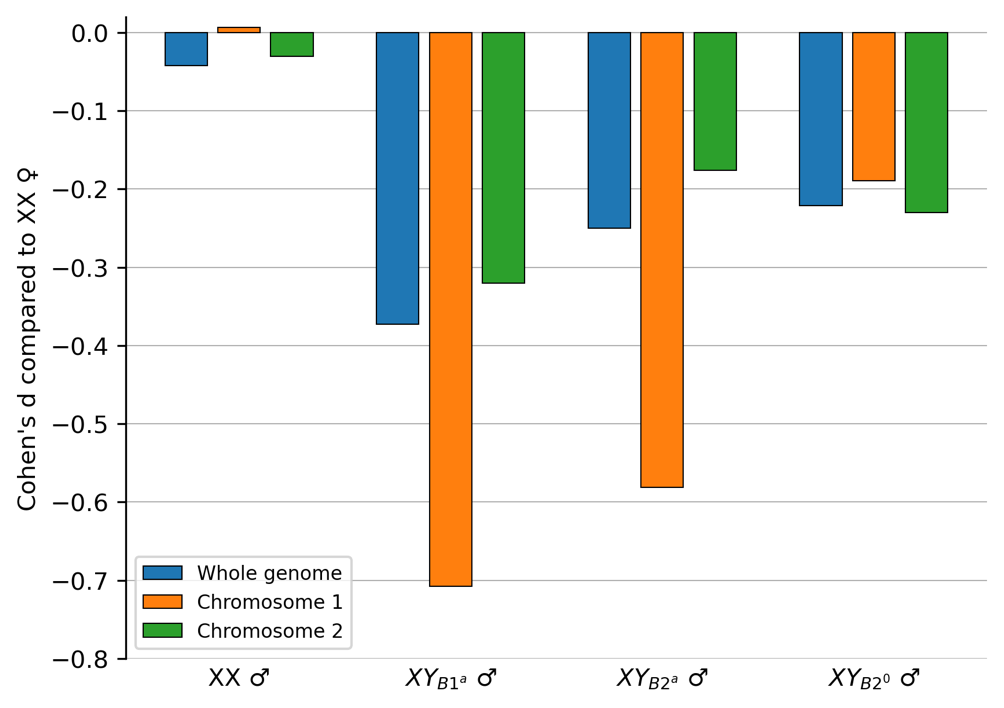

**Figure S3**. Nucleotide diversity (π) calculated in chromosomes and whole genome for the XX female pool and four male genotype pools: XX, XY_B1_^a^, XY_B2_^a^, and XY_B2_^⁰^ males. Black horizontal bars indicate the median π for each pool. Pairwise effect sizes shown above the plots represent Cohen’s d relative to the XX female pool. (a) shows π of the sex chromosome (chromosome 1) across 5 pools. (b) shows π diversity of autosomes across 5 pools, excluding chromosome 1. (c) summarizes Cohen’s d for each male pool compared with the XX female pool across the whole genome, sex chromosome, and chromosome 2. Negative values indicate reduced nucleotide diversity relative to the XX female pool.

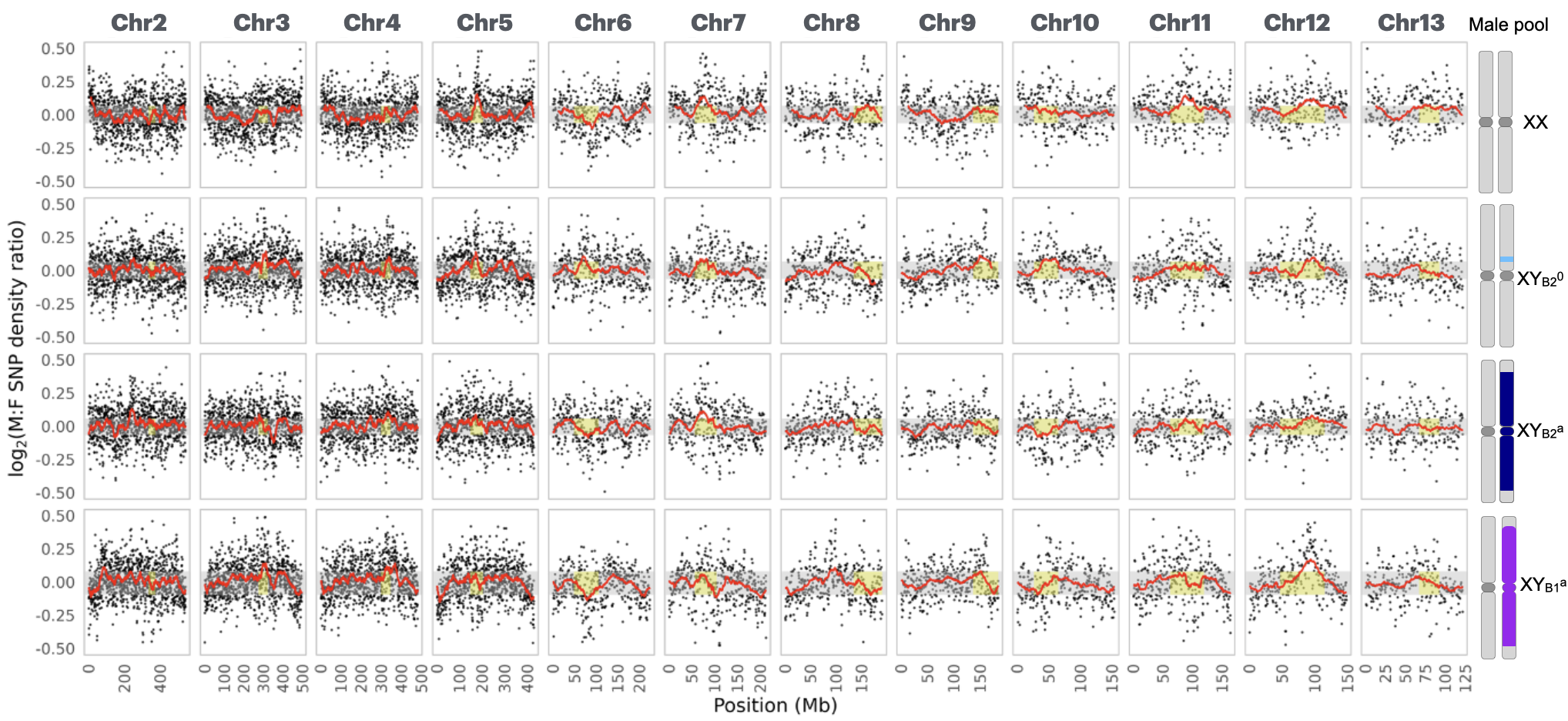

**Figure S4**: Log_2_ male-to-female SNP density ratio between each male and the female pool. Each male pool (XX, YB_2_^0^, YB_1_^a^, and YB_2_^a^) was compared to the female pool along each of the 12 autosomes. The dot represents the average SNP density ratio within 1 Mb overlapping window. The red line indicates the rolling mean (window size = 50 Mb). The grey rectangle marks the central 95% of the rolling mean of autosomal values. Yellow zones indicate tentative centromeric regions.

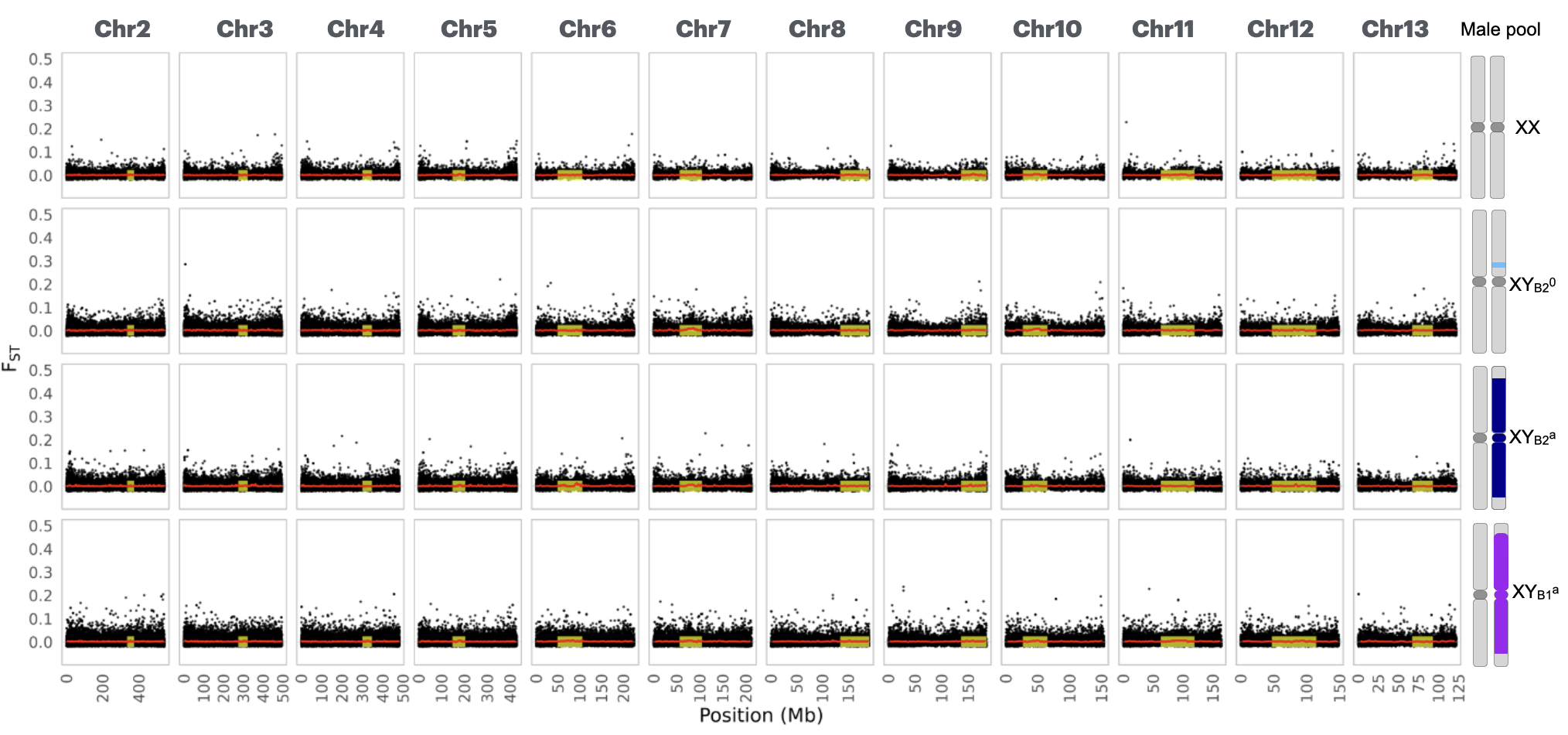

**Figure S5**: F_ST_ between each male pool and the female pool. Each male pool (XX, YB_2_^0^, YB_1_^a^, and YB_2_^a^) was compared to the female pool along each of the 12 autosomes. The dot represents the average F_ST_ within a 10kb overlapping window (stride = 5kb) between each male and female pools. The grey rectangle marks the central 95% of the rolling mean of autosomal values. Yellow zones indicate tentative centromeric regions.

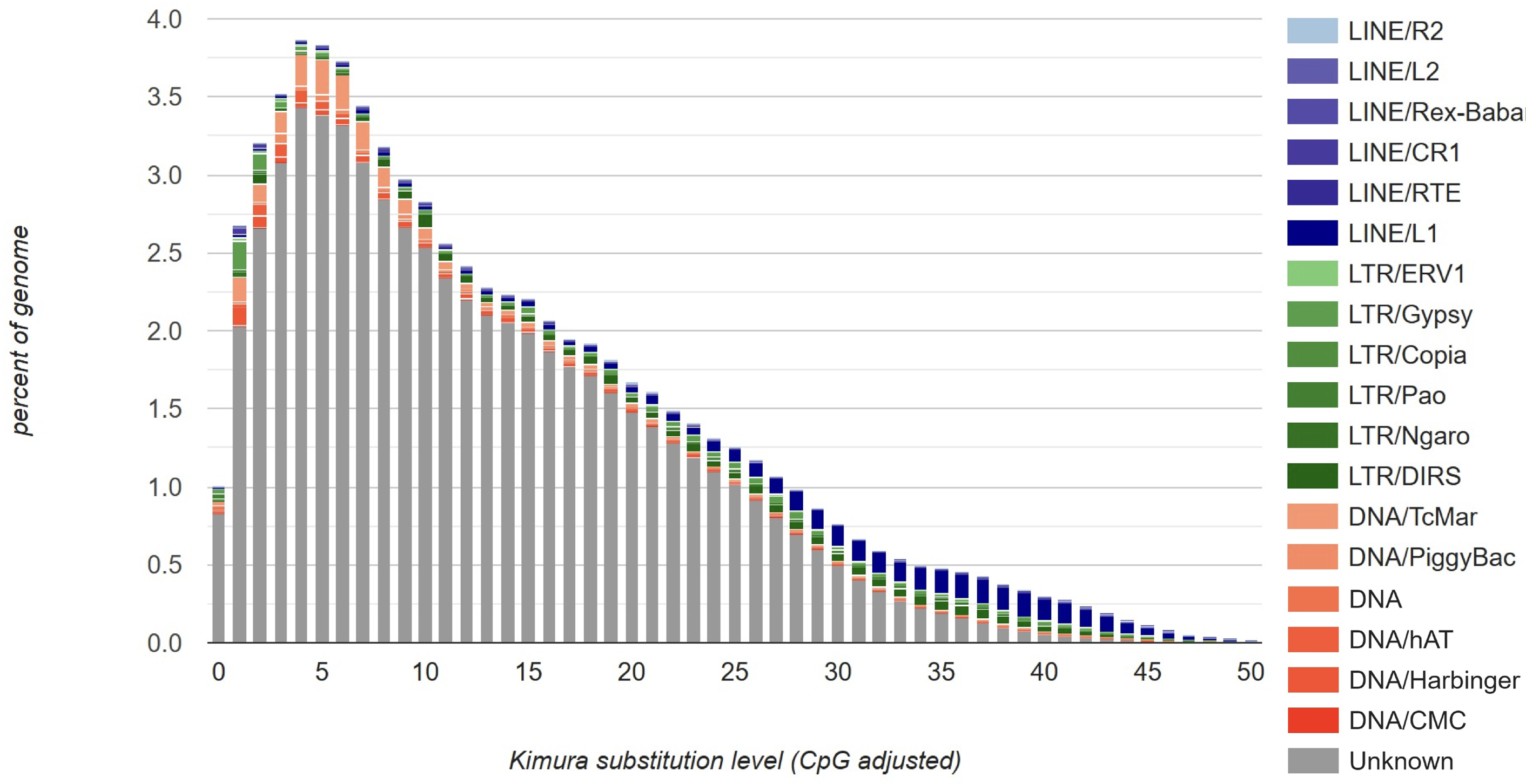

**Figure S6.** TE divergence landscape inferred by the kimura substitution level, and the smaller values indicate low divergence. The TE library is conducted with *de novo* annotation of the reference genome *R. temporaria*.

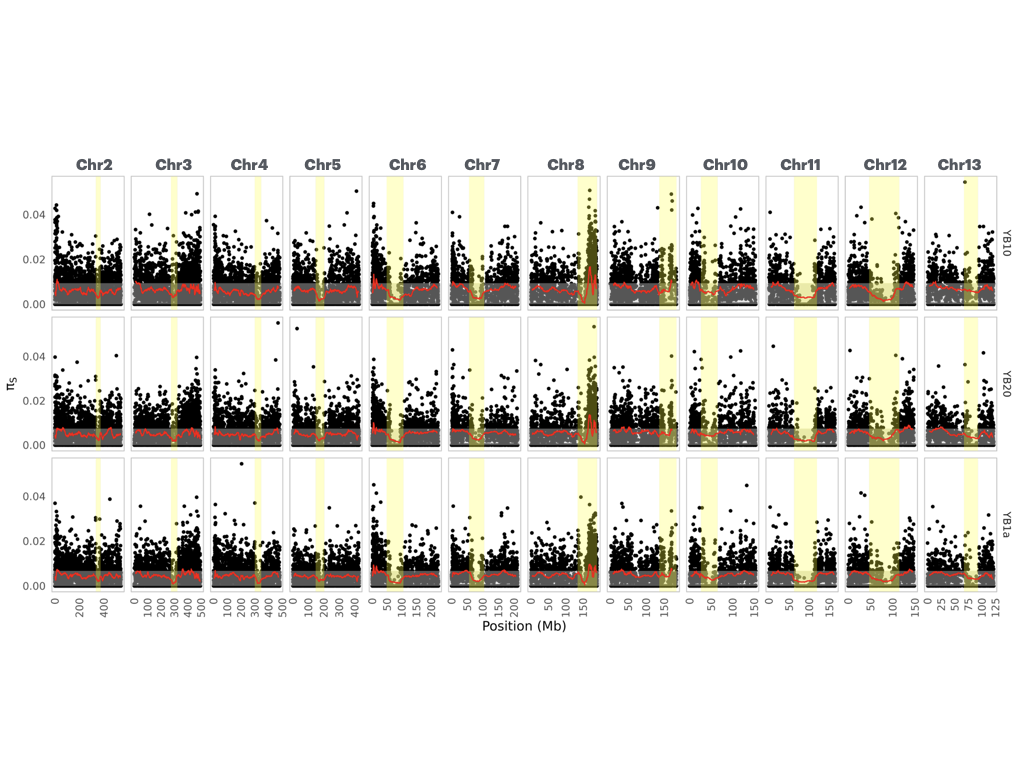

**Figure S7**. Computed π_S_ along each of 12 autosomes between each of the three type of Y haplotypes and the reference XX genome. The grey rectangle marks the central 95% of the rolling mean of autosomal values. Yellow zones indicate tentative centromeric regions.

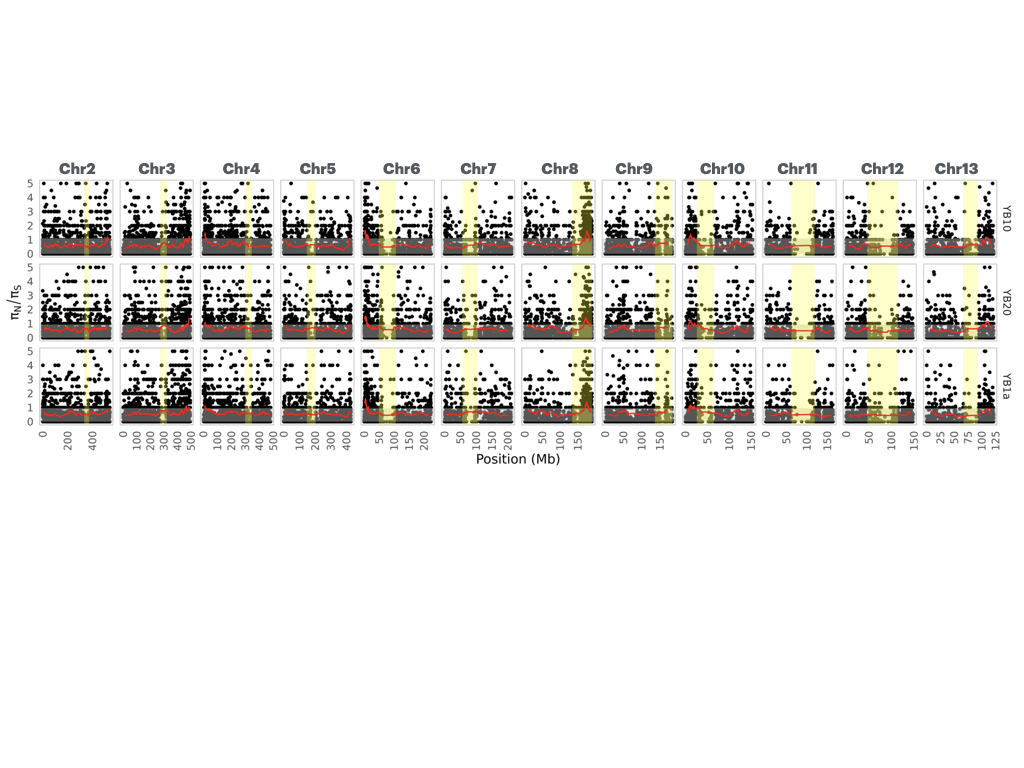

**Figure S8**. Computed π_N_/π_S_ along each of the 12 autosomes between each of the three type of Y haplotypes and the reference XX genome. The grey rectangle marks the central 95% of the rolling mean of autosomal values. Yellow zones indicate tentative centromeric regions.

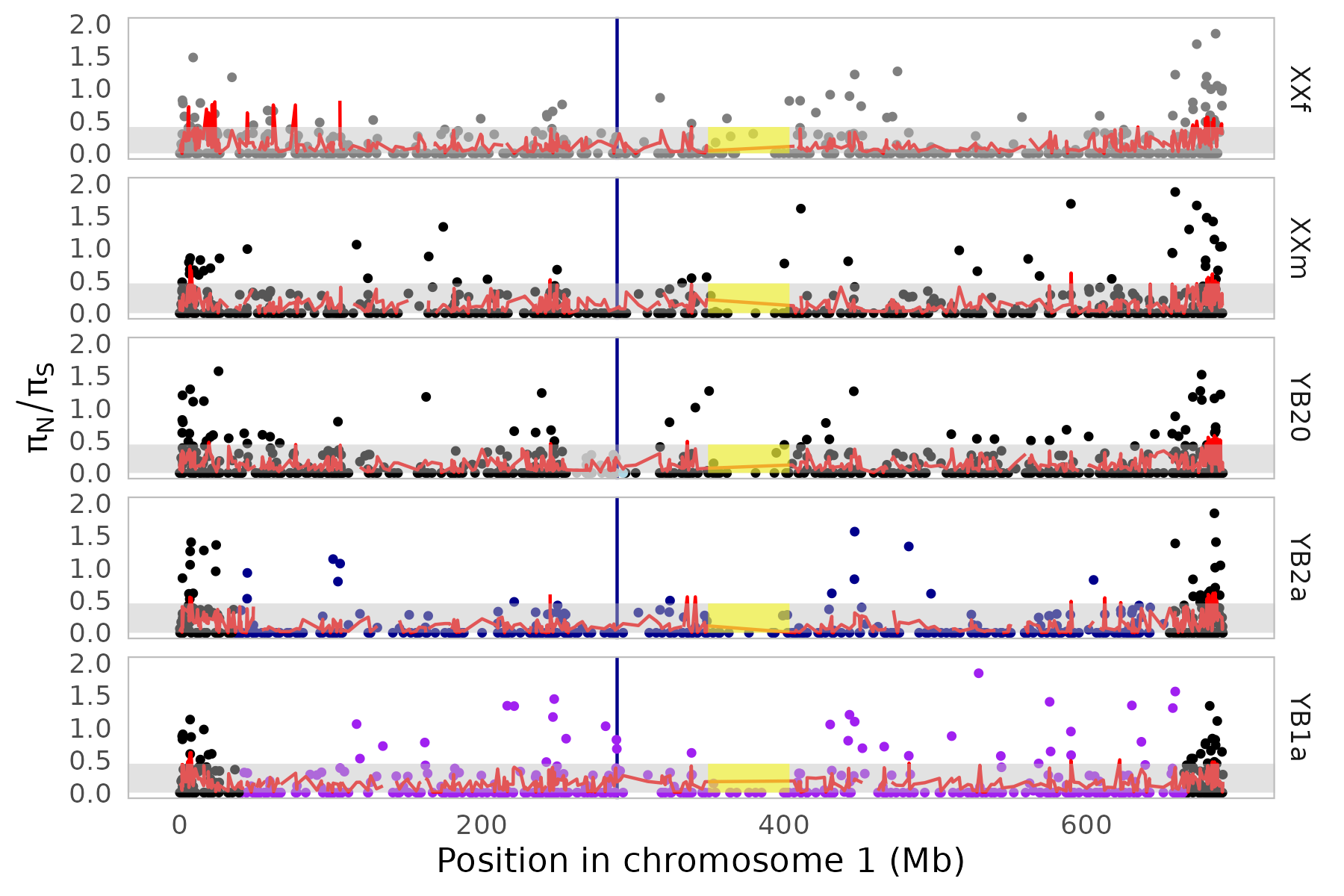

**Figure S9**. Computed π_N_/π_S_ between each of the five pool and the reference XX genome. The vertical blue line is the region including *Dmrt1* and *Dmrt3* genes.

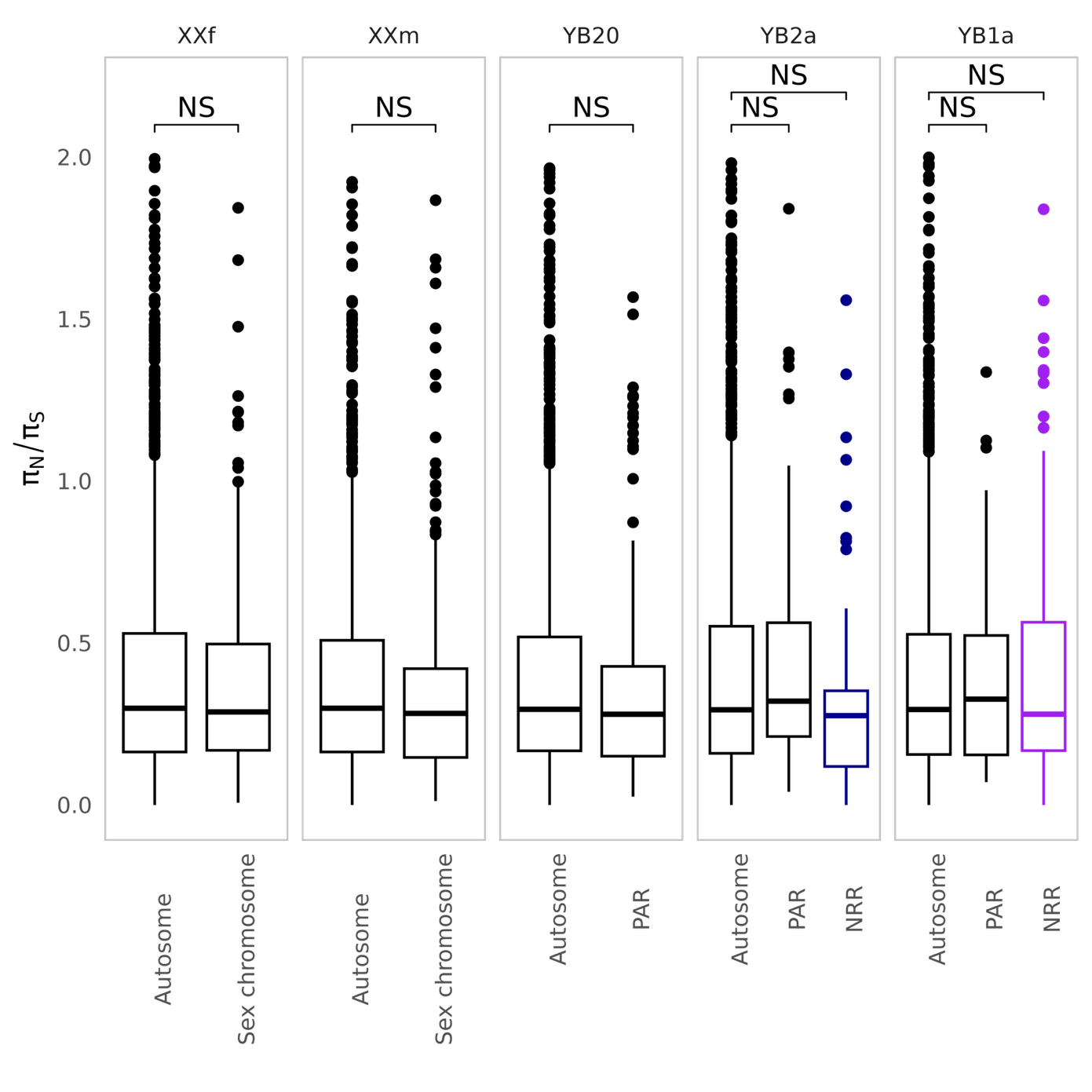

**Figure S10**. Boxplot of π_N_/π_S_ values between genomic compartments with significance indicated from permutation test.

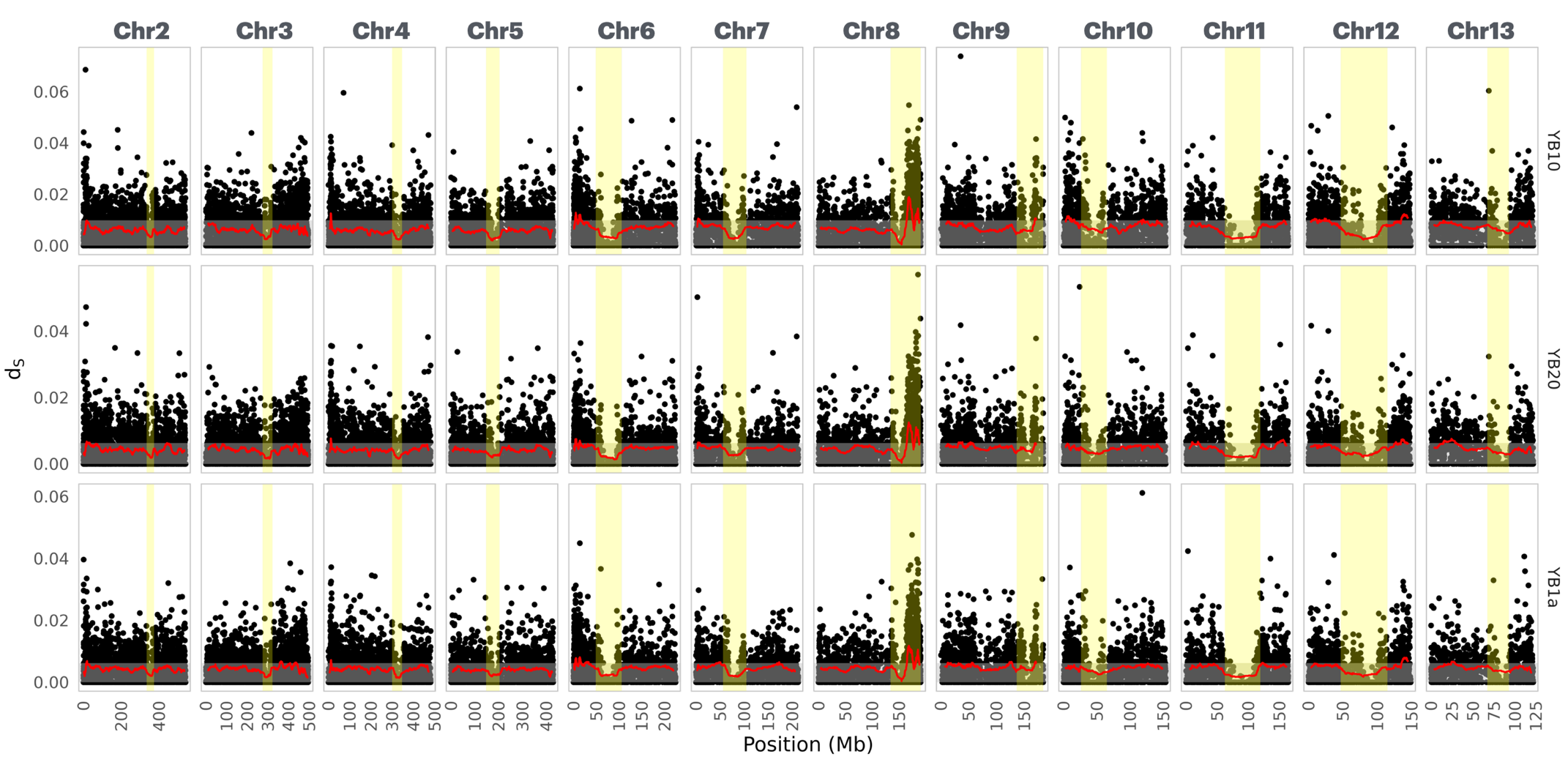

**Figure S11**. Computed d_S_ along each of 12 autosomes between each of the three type of Y haplotypes and the reference XX genome. The grey rectangle marks the central 95% of the rolling mean of autosomal values. Yellow zones indicate tentative centromeric regions.

(a)

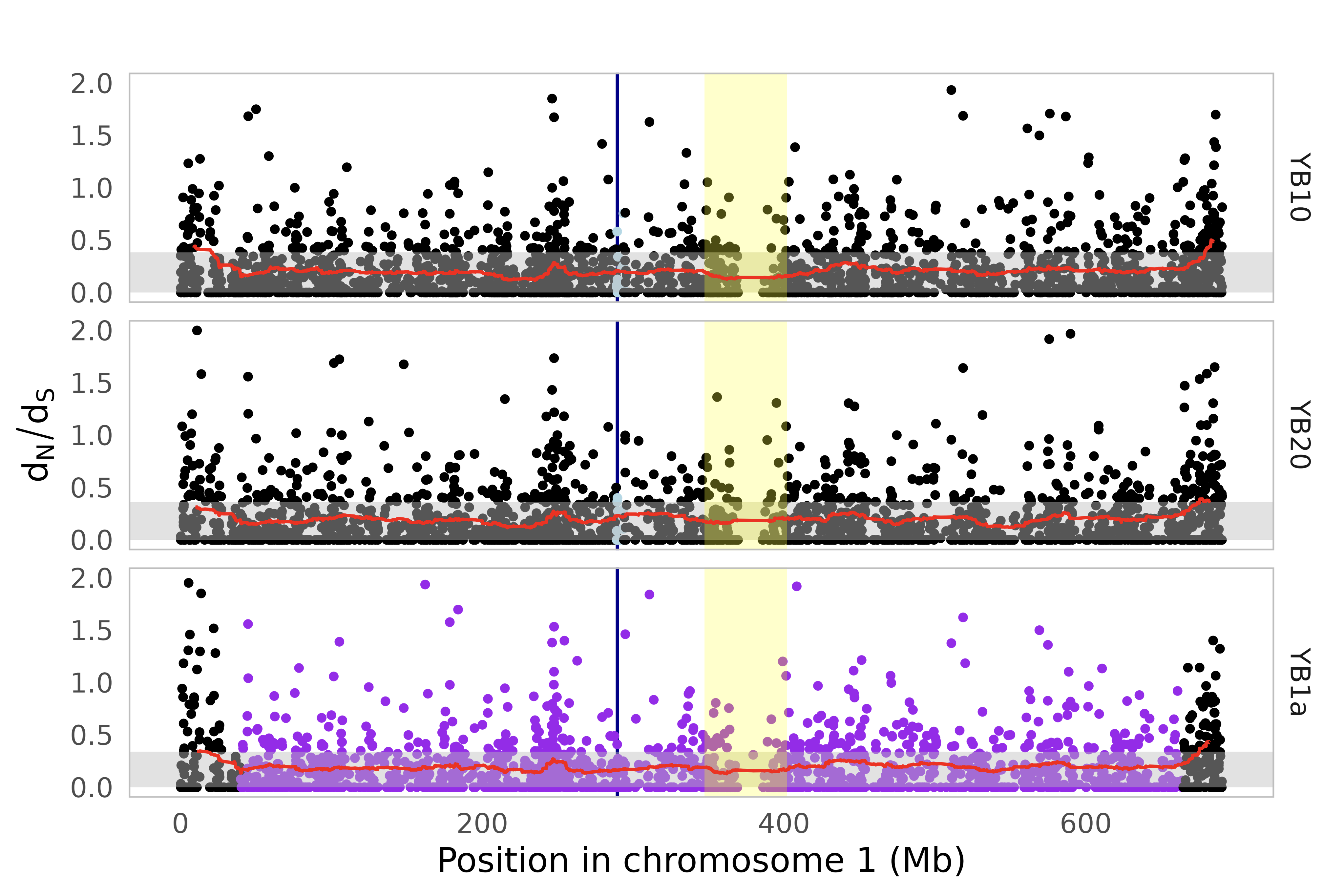

(b)

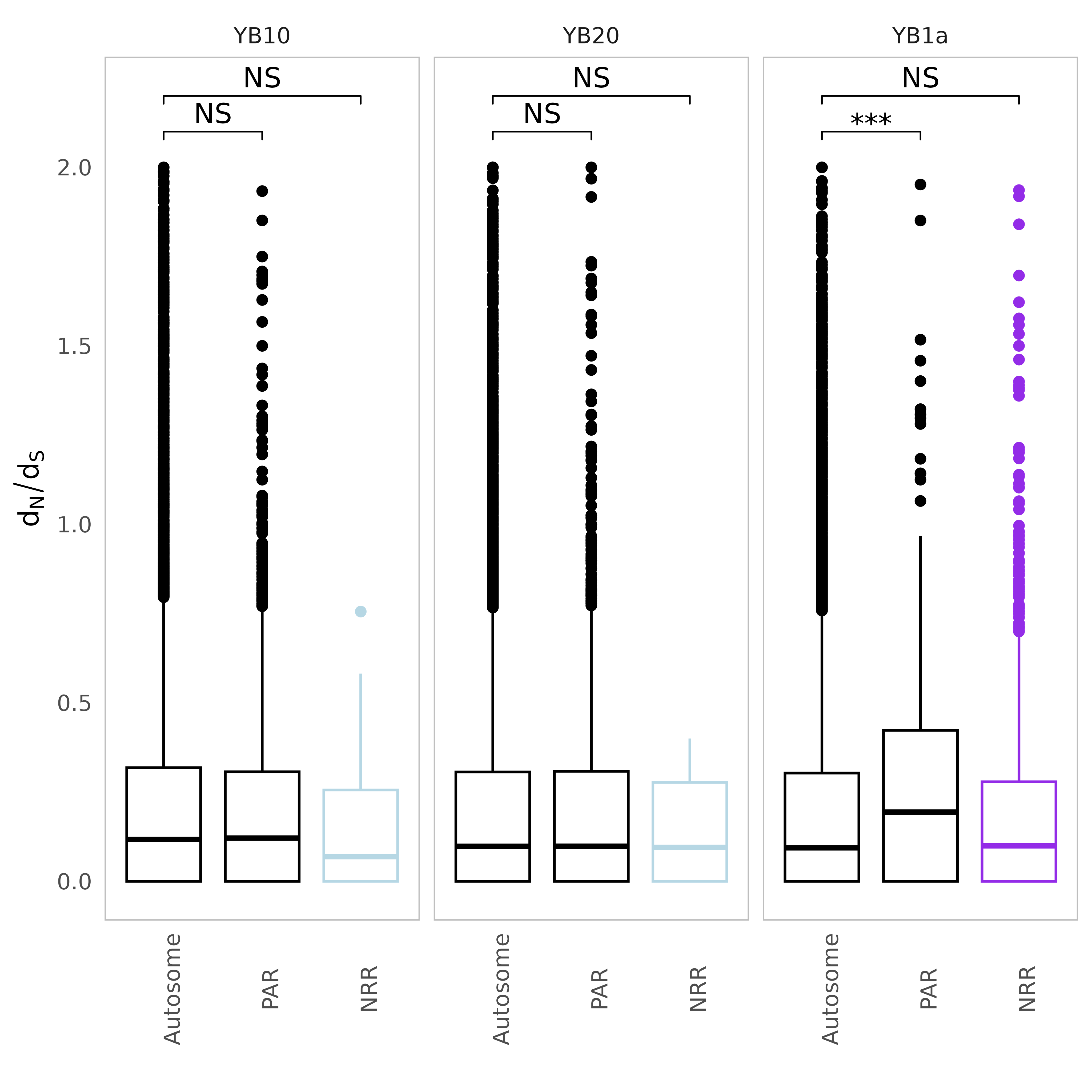

**Figure S12**. Computed d_N_/d_S_ between each of the three types of Y haplotypes and the reference XX genome. (a) d_N_/d_S_ along the sex chromosome. The vertical blue line is the region including *Dmrt1* and *Dmrt3* genes. (b) Boxplot of d_N_/d_S_ values between genomic compartments with significance indicated from permutation test.

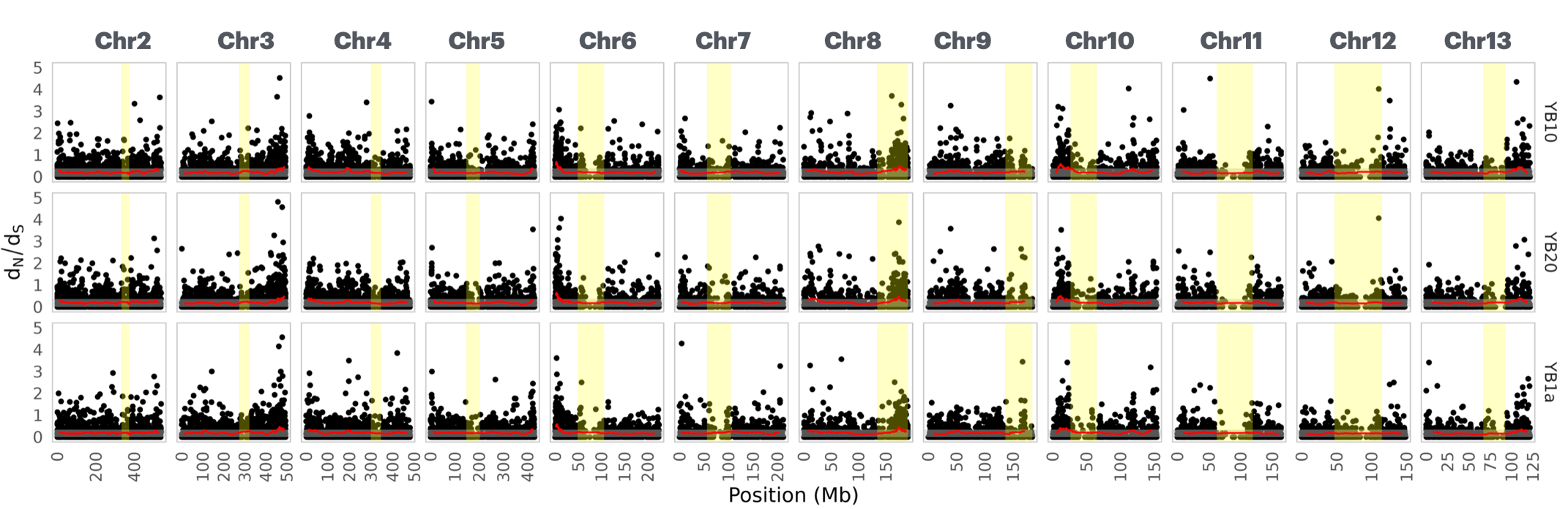

**Figure S13**. Computed d_N_/d_S_ along each of the 12 autosomes between each of the three type of Y haplotypes and the reference XX genome. The grey rectangle marks the central 95% of the rolling mean of autosomal values. Yellow zones indicate tentative centromeric regions.

a)

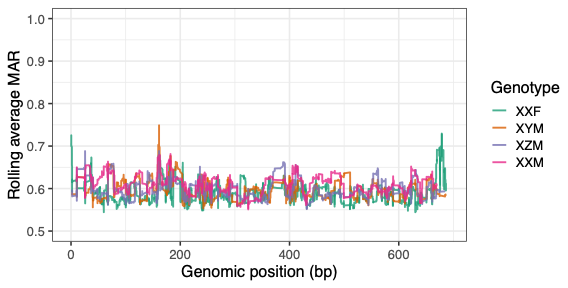

b)

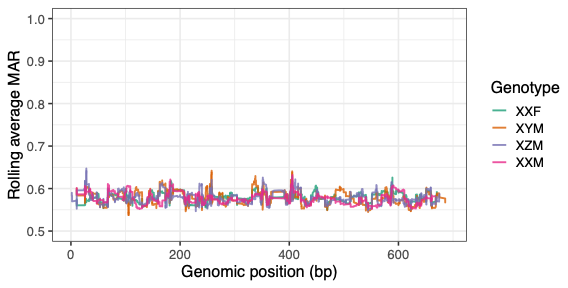

**Figure S14**. Major allele ratio along the sex chromosome for the female, and three types of males in both gonad (a) and brain tissues (b) from the Meitreile population. We computed the major allele ratio in sliding windows of 500 SNPs using the rollmean function in the R package zoo.

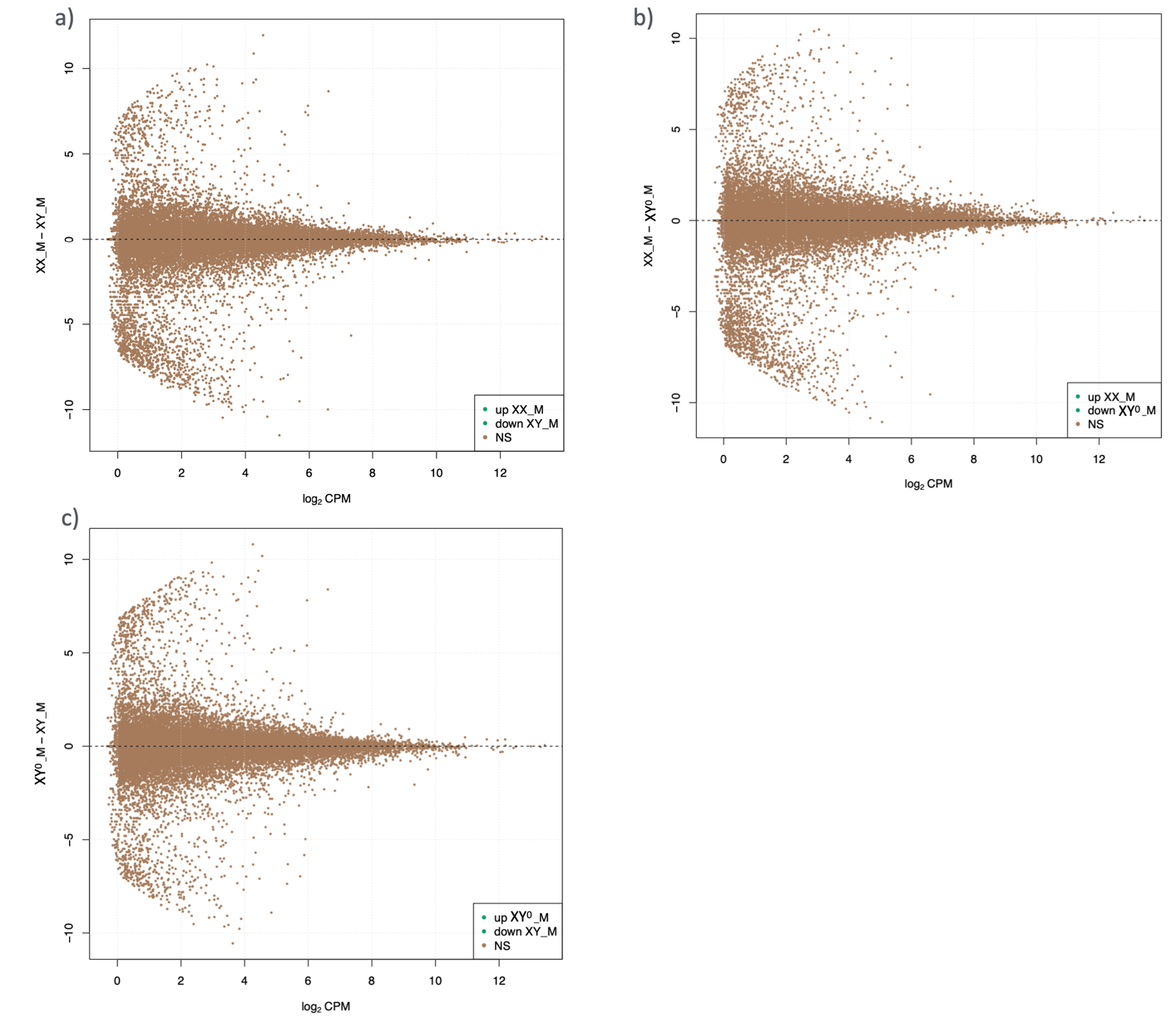

**Figure S15**. Differential expression analyses in testes for pairwise male-male comparison among all three genotypes with varying Y differentiation levels. (a) XX male vs XY male; (b) XX male vs XY^0^ male; and (c) XY^0^ male vs XY male.

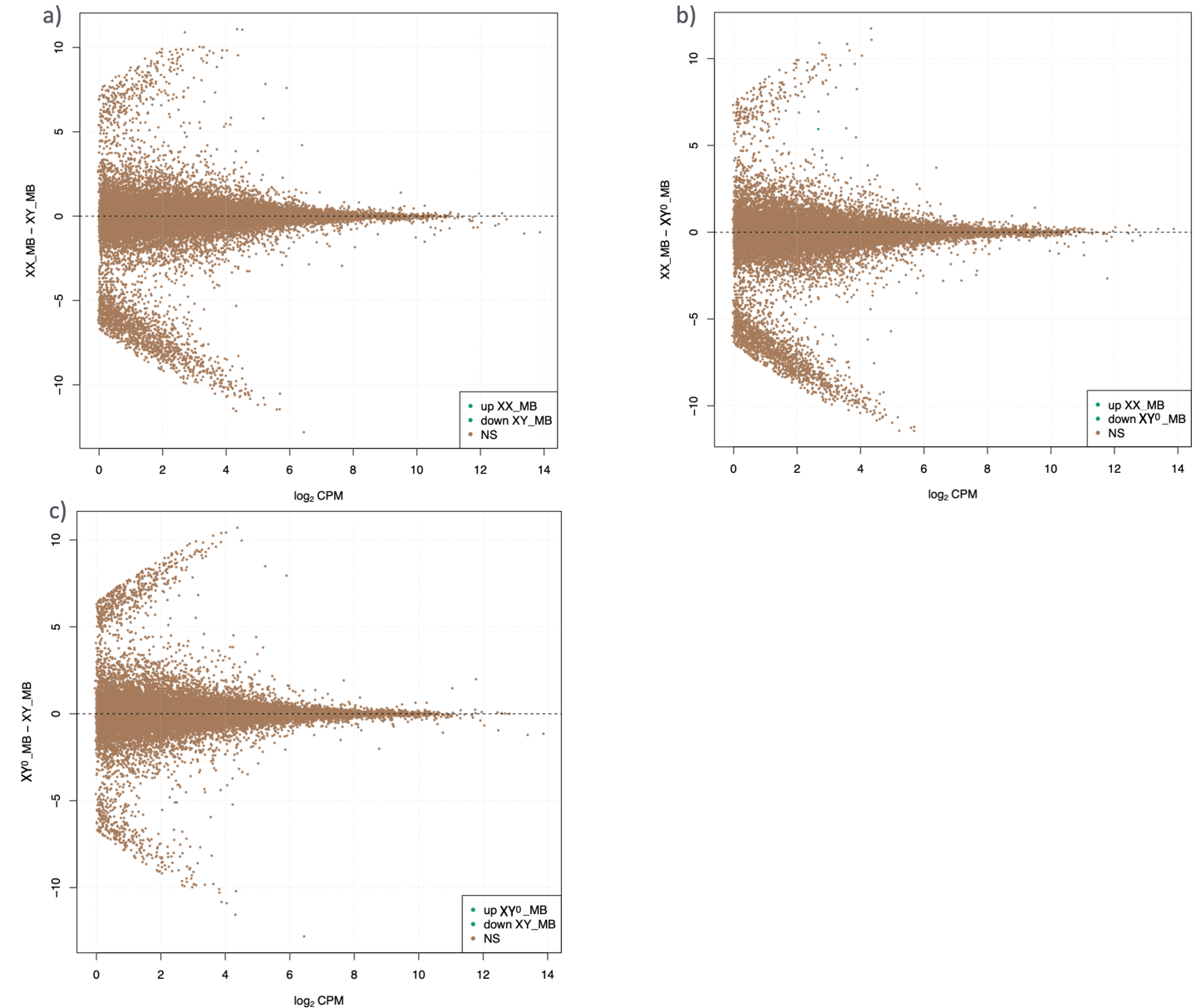

**Figure S16**. Differential expression analyses in brain tissues for pairwise male-male comparison among all three genotypes with varying Y differentiation levels. (a) XX male vs XY male; (b) XX male vs XY^0^ male; and (c) XY^0^ male vs XY male.

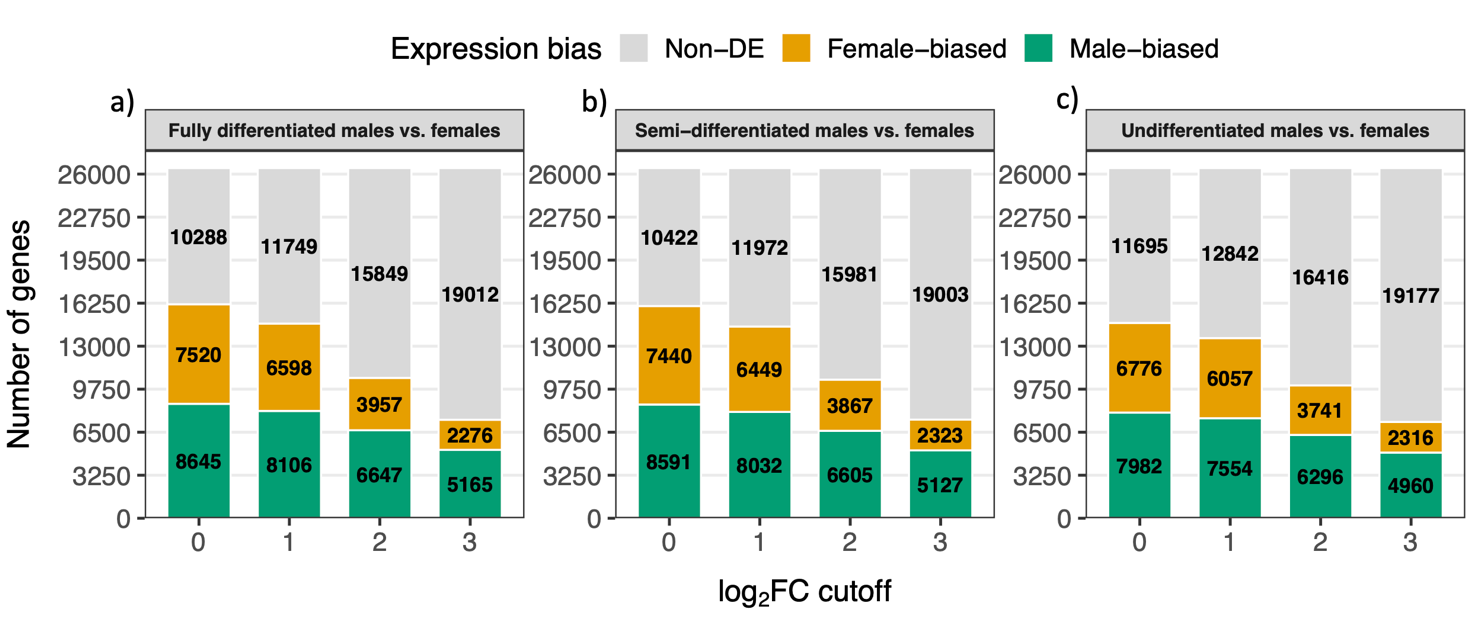

**Figure S17**. Summary number of differential expression analyses in gonad tissues between females and males with all three genotypes with varying Y differentiation levels, with FDR < 0.05 and |log2(male/female)| cutoff of >= 0, >= 1, >= 2 and >= 3 respectively. (a) XY male vs XX female; (b) XY^0^ male vs XX female; and (c) XX male vs XX female.

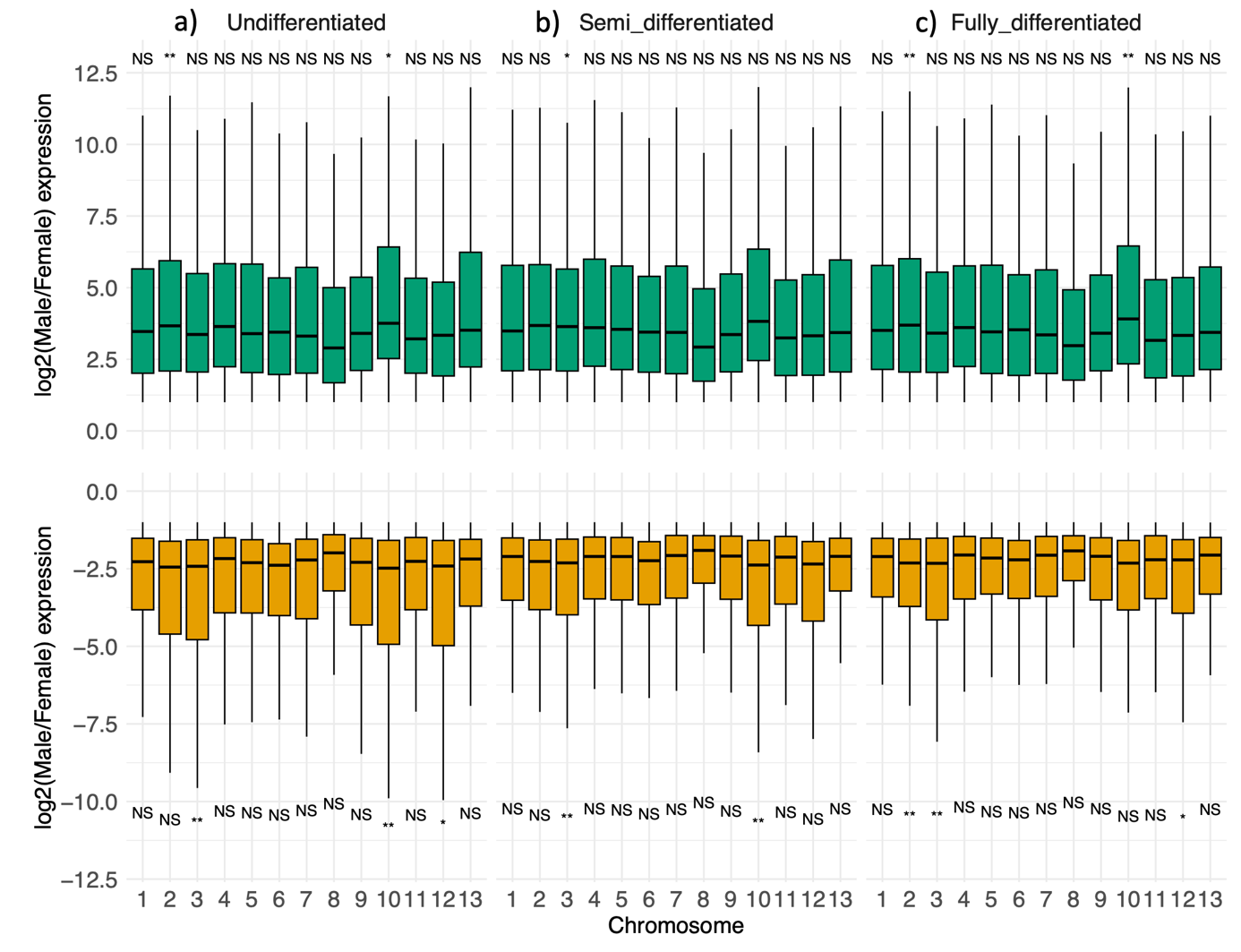

**Figure S18**. Boxplot for distribution of differentially expressed genes in gonad tissues between females and males with all three genotypes with varying Y differentiation levels, as well as the permutation tests, with FDR < 0.05 and |log2(male/female)| cutoff >= 1. Permutation test (FDR-corrected): * < 0.05, ** < 0.01, *** < 0.001. (a) XX male vs XX female; (b) XY^0^ male vs XX female; and (c) XY male vs XX female.

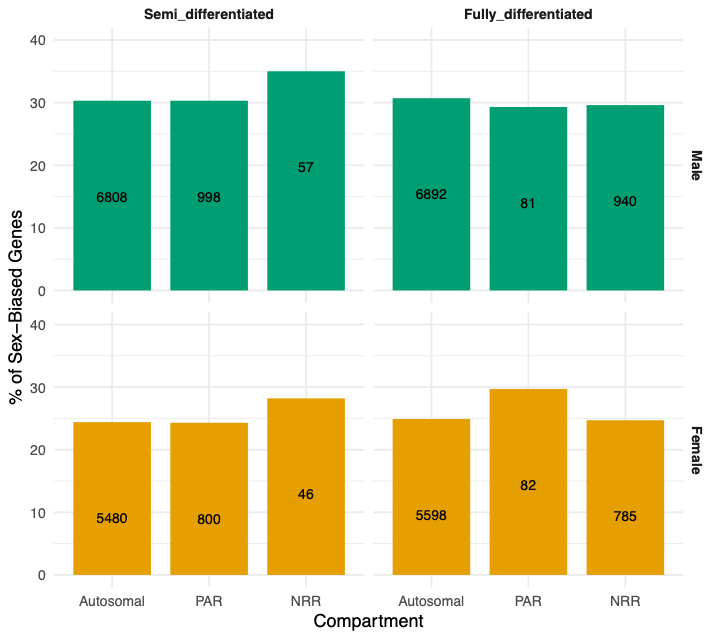

**Figure S19**. Barplot for the number of sex-biased genes in gonad tissues between females and males across genomic compartments with full-differentiated Y and semi-differentiated Y. Orange are female-biased genes, green are male-biased genes.

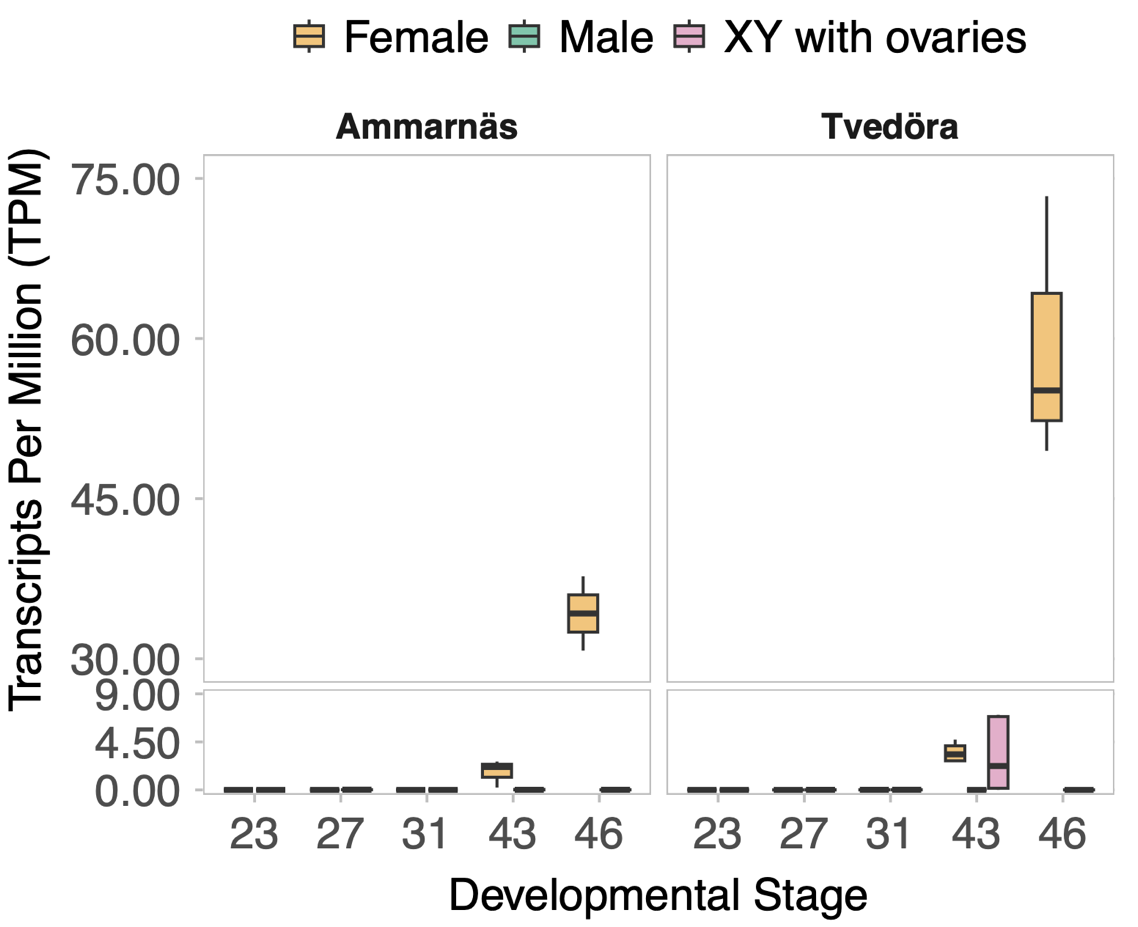

**Figure S20**. Loc120943612 gene expression in TPM (transcript per million) across developmental stages (G23, 27, 31, 43 and 46) from the published RNAseq data of the whole individual, in populations with fully-differentiated XY in Ammarnäs (north Sweden), with semi-differentiated XY^0^ males in Tvedöra (south Sweden).

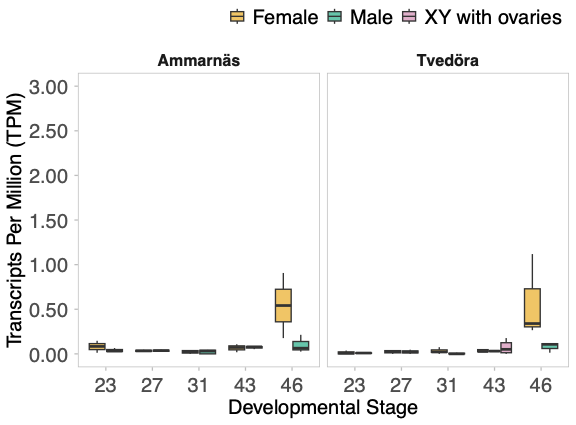

**Figure S21**. *Dmrt1* gene expression in TPM (transcript per million) across developmental stages (G23, 27, 31, 43 and 46) from the published RNAseq data of the whole individual, in populations with fully-differentiated XY in Ammarnäs (north Sweden), with semi-differentiated XY^0^ males in Tvedöra (south Sweden).

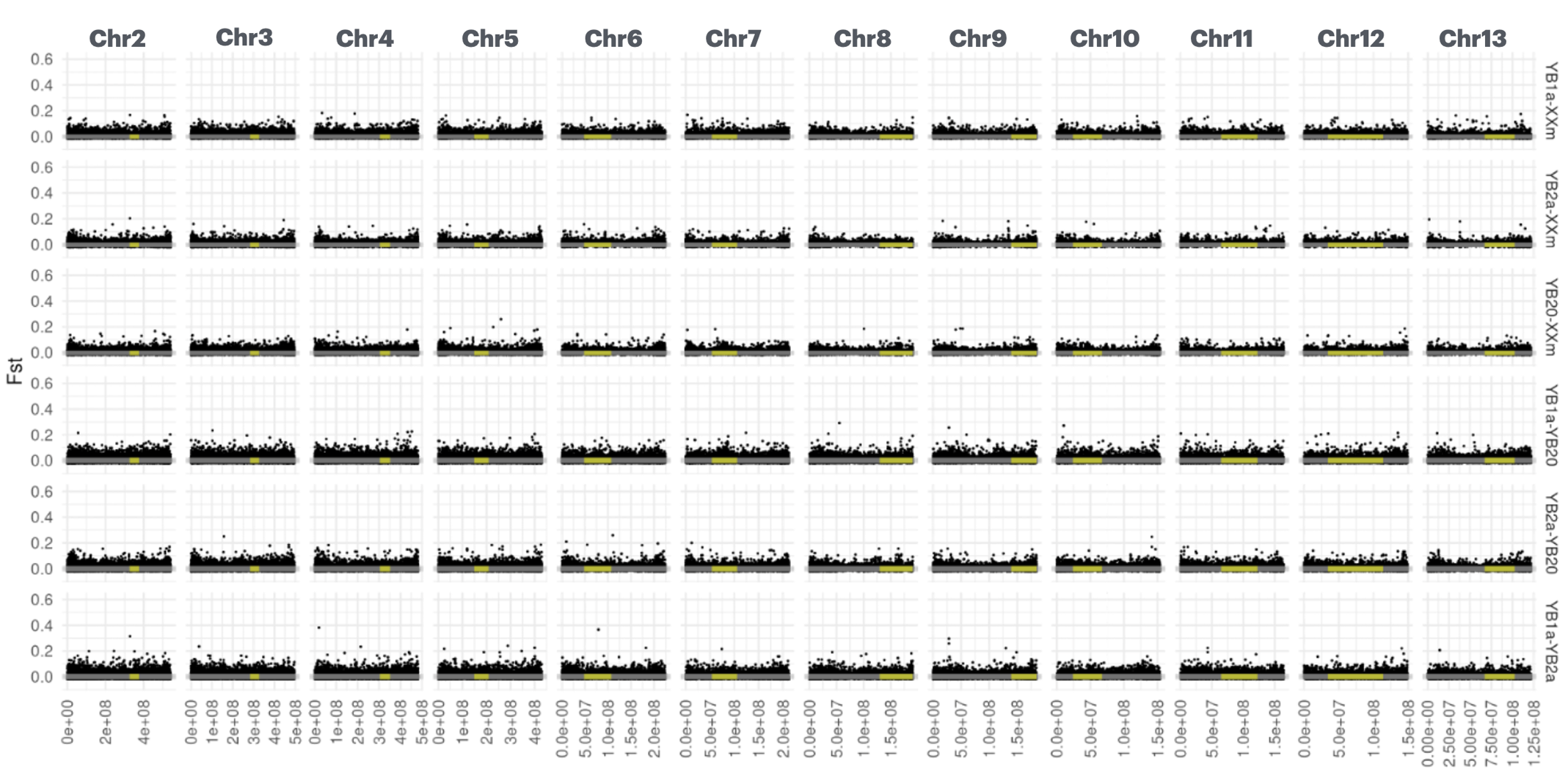

### **Figure S22**. Autosomal F_ST_ among male pools. For each of the 12 autosomal contigs and among male pools (XXm, YB20, YB1a, and YB2a), each dot represents the average F_ST_ between the male pools within a 10kb overlapping window (stride = 5kb). The grey rectangle marks the central 95% of the autosomal value distribution. Yellow intervals show centromeric regions.

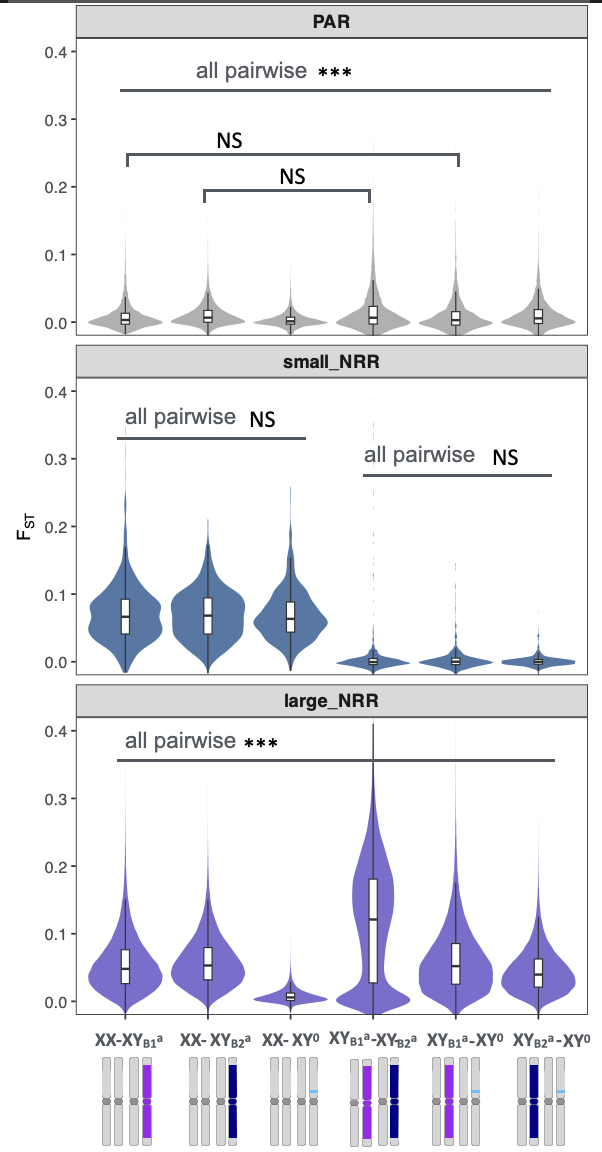

**Figure S23.** Distribution of genetic differentiation, F_ST_, across sex-chromosome compartments including pseudoautosomal region (PAR), small non-recombining region (small NRR), and large non-recombining region (large NRR), for pairwise male–male comparisons. F_ST_ values are shown as violin plots with embedded boxplots. No significant differentiation was detected among male pools in small NRR, whereas differed significantly among comparisons in the large NRR. Statistical significance was assessed using pairwise Wilcoxon tests with Benjamini–Hochberg correction (*** P < 0.001; NS, not significant).

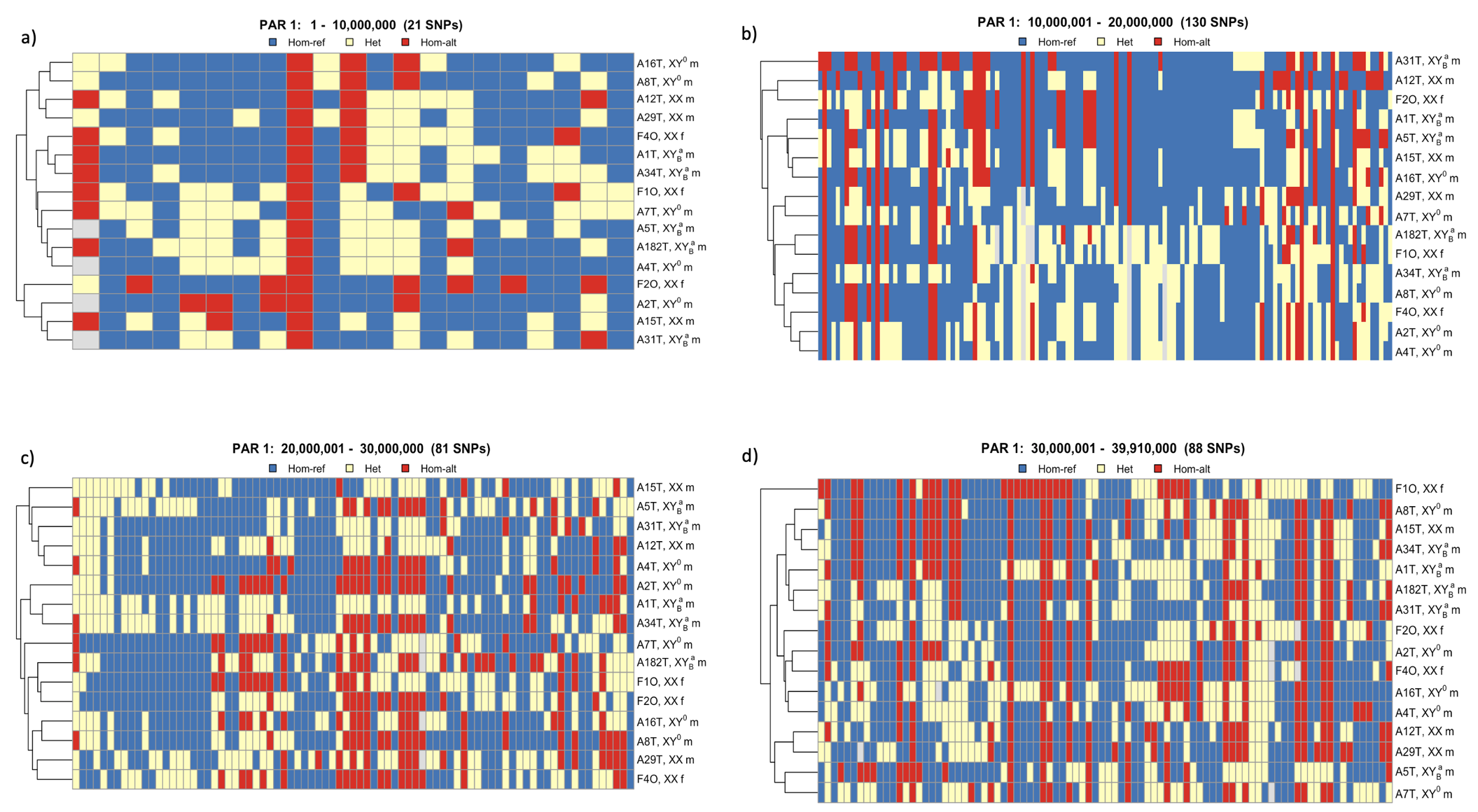

**Figure S24**. Clustering dendrograms based on SNPs derived from RNA-seq read mappings of male and female gonad and brain tissues to the reference genome, illustrating patterns of homozygosity and heterozygosity along PAR1 region. Each dendrogram was generated from SNP sets extracted using a 10-Mb sliding window (a, b, c and d accordingly).

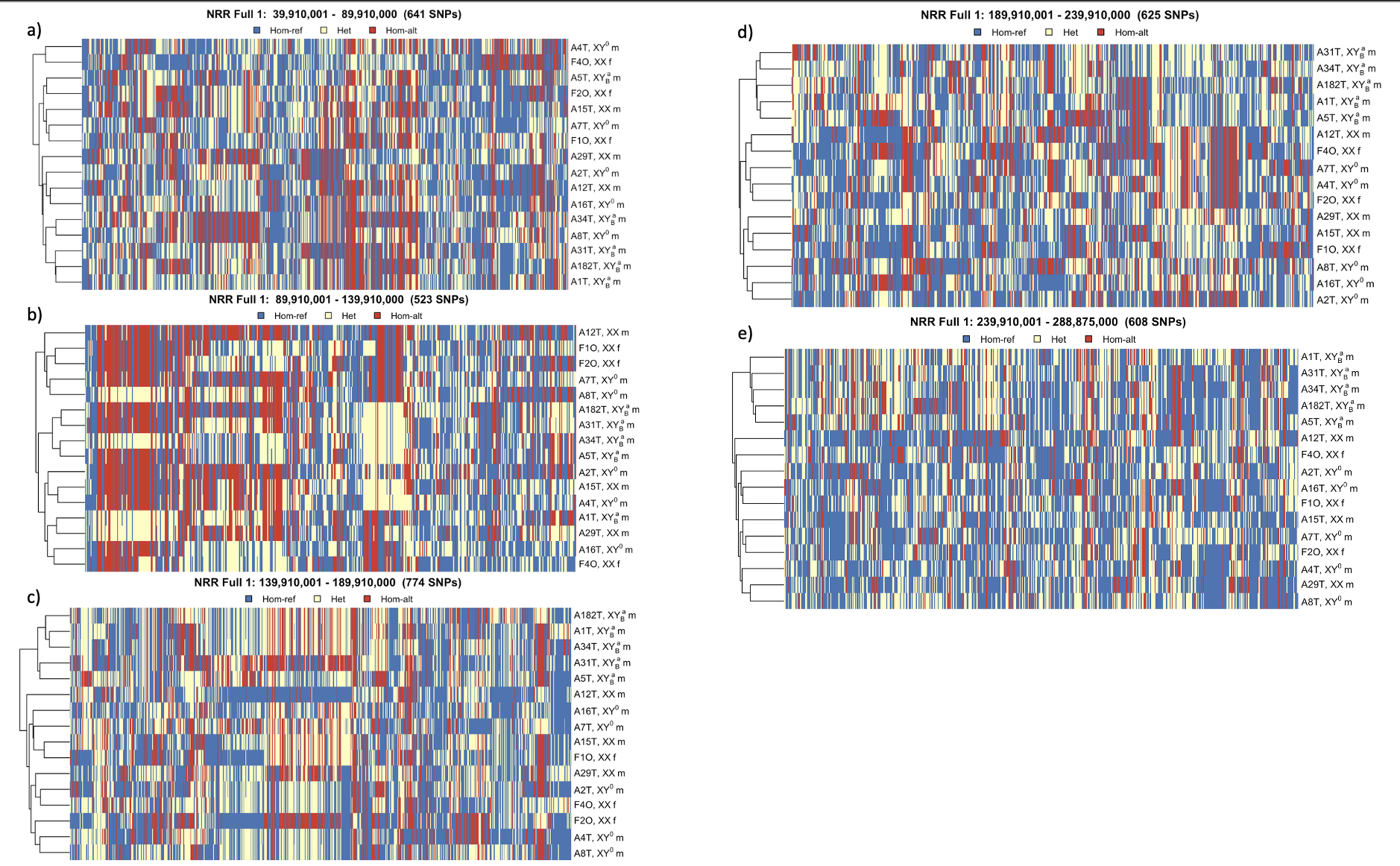

**Figure S25**. Clustering dendrograms based on SNPs derived from RNA-seq read mappings of male and female gonad and brain tissues to the reference genome, illustrating patterns of homozygosity and heterozygosity along large NRR part1. Each dendrogram was generated from SNP sets extracted using 50 Mb sliding window (a, b, c, d accordingly).

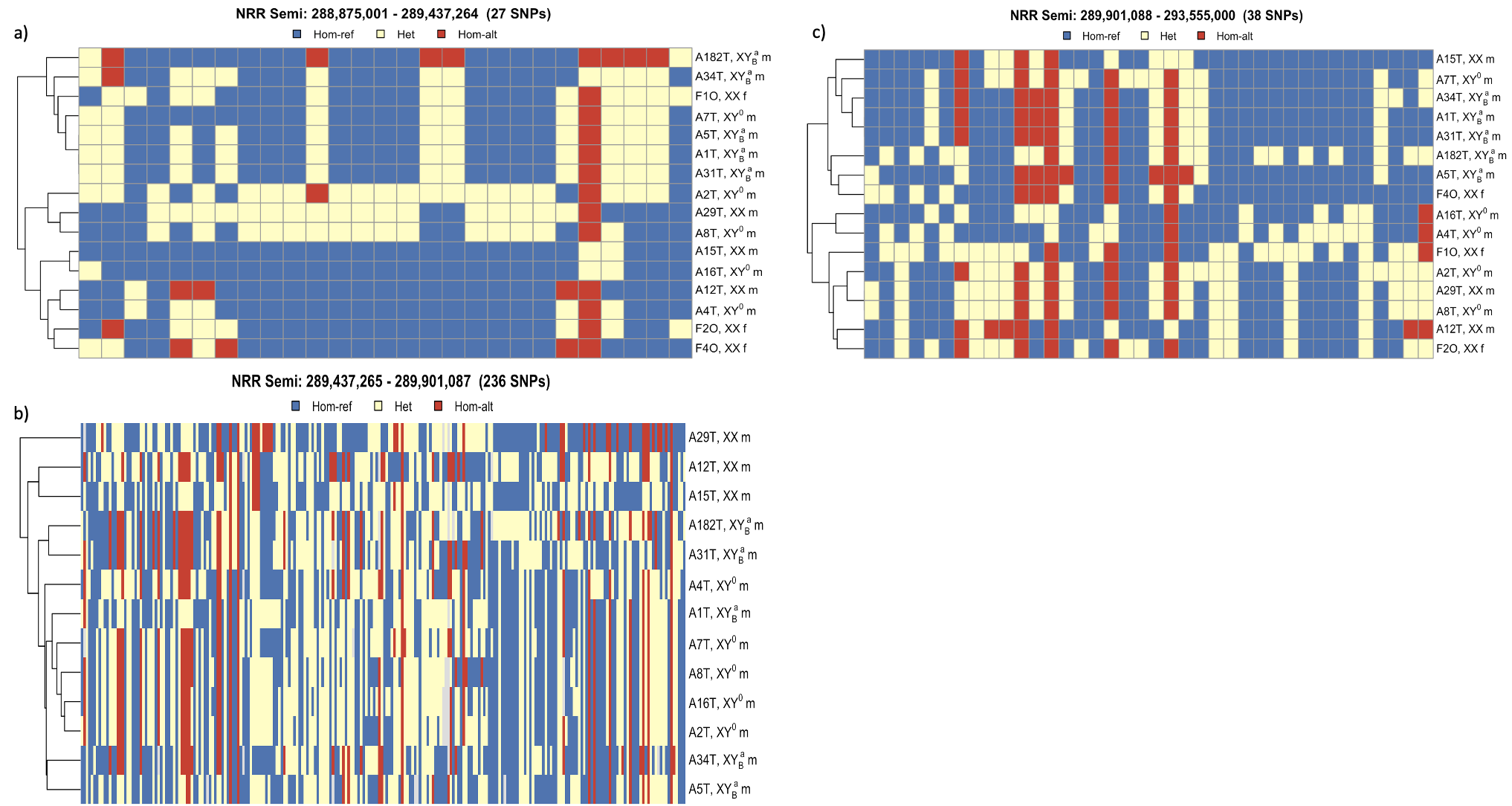

**Figure S26**. Clustering dendrograms based on SNPs derived from RNA-seq read mappings of male and female gonad and brain tissues to the reference genome, illustrating patterns of homozygosity and heterozygosity along PAR1. Each dendrogram was generated from SNP sets extracted using 3 divided window, genes upstream of *Dmrt1* (a), *Dmrt1, Dmrt3* and *Dmrt2* (b), and genes downstream of *Dmrt2* (c).

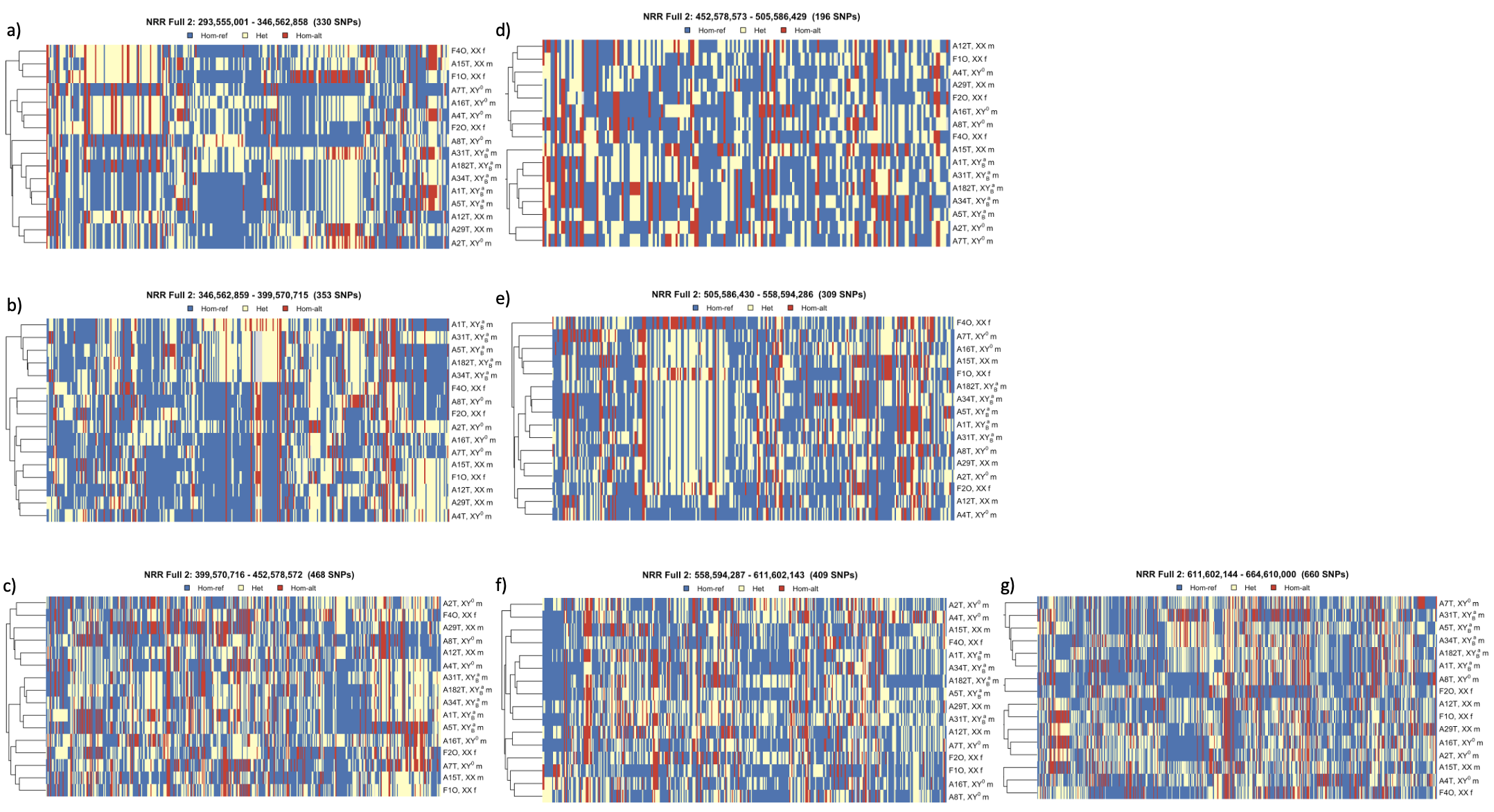

**Figure S27**. Clustering dendrograms based on SNPs derived from RNA-seq read mappings of male and female gonad and brain tissues to the reference genome, illustrating patterns of homozygosity and heterozygosity along large NRR part2. Each dendrogram was generated from SNP sets extracted using 50 Mb sliding window (a, b, c, d, e, f, and g accordingly).

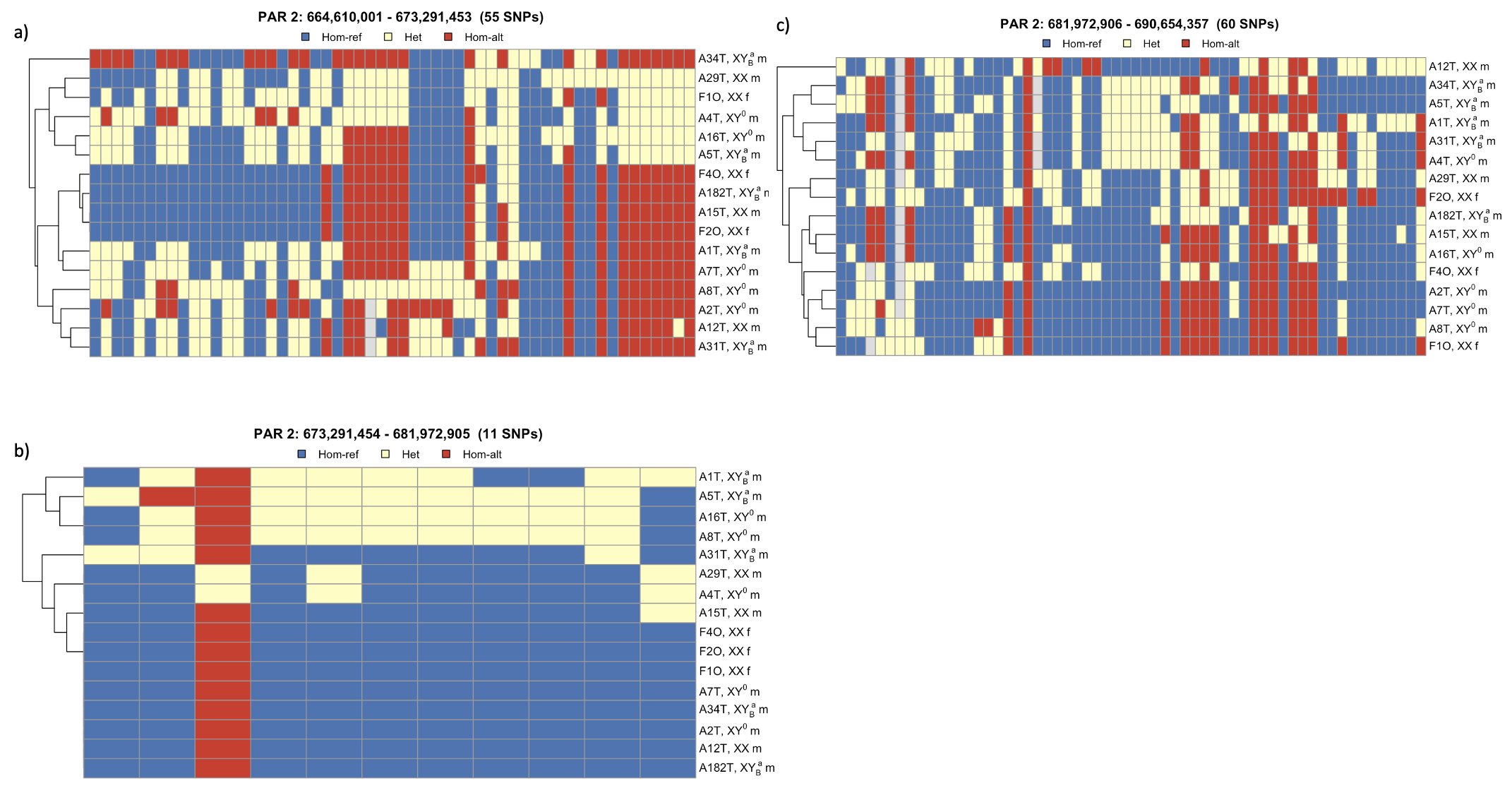

**Figure S28**. Clustering dendrograms based on SNPs derived from RNA-seq read mappings of male and female gonad and brain tissues to the reference genome, illustrating patterns of homozygosity and heterozygosity along PAR2 region. Each dendrogram was generated from SNP sets extracted using a 10-Mb sliding window (a, b, c accordingly)
